# Identification of Covalent Cyclic Peptide Inhibitors Targeting Protein-Protein Interactions Using Phage Display

**DOI:** 10.1101/2024.11.08.622749

**Authors:** Sijie Wang, Franco F. Faucher, Matilde Bertolini, Heeyoung Kim, Bingchen Yu, Li Cao, Katharina Roeltgen, Scott Lovell, Varun Shanker, Scott D. Boyd, Lei Wang, Ralf Bartenschlager, Matthew Bogyo

**Author notes:** Correspondence: Matthew Bogyo.

## Abstract

Peptide macrocycles are promising therapeutics for a variety of disease indications due to their overall metabolic stability and potential to make highly selective binding interactions with targets. Recent advances in covalent macrocycle peptide discovery, driven by phage and mRNA display methods, have enabled the rapid identification of highly potent and selective molecules from large libraires of diverse macrocycles. However, there are currently limited examples of macrocycles that can be used to disrupt protein-protein interactions and even fewer examples that function by formation of a covalent bond to a target protein. In this work, we describe a directed counter-selection method that enables identification of covalent macrocyclic ligands targeting a protein-protein interaction using a phage display screening platform. This method utilizes binary and ternary screenings of a chemically modified phage display library, employing the stable and weakly reactive aryl fluorosulfate electrophile. We demonstrate the utility of this approach using the SARS-CoV-2 Spike-ACE2 protein-protein interaction and identify multiple covalent macrocyclic inhibitors that disrupt this interaction. The resulting compounds displayed antiviral activity against live virus that was irreversible after washout due to the covalent binding mechanism. These results highlight the potential of this screening platform for developing covalent macrocyclic drugs that disrupt protein-protein interactions with long lasting effects.

## INTRODUCTION

Over the past two decades, macrocycles including cyclic peptides, have gained attention as potentially viable therapeutic drug modalities.^1,2^ Compared to conventional small-molecule drugs and large antibody-based biological macromolecules, macrocycles have unique advantages. Their size (500-2000 Da), flexibility, and structural properties provide unique capabilities to interact with large and flat protein interfaces that are typically challenging for small molecules. Furthermore, macrocycles can present binding surfaces that are similar in size, affinity and specificity to antibodies,^3,4^ while still offering many of the advantages of small-molecules such as oral bioavailability, synthetic accessibility and potential to be stockpiled for future use. Macrocyclic peptides can also be optimized for high metabolic stability, cell-membrane permeability, and conformational rigidity.^5–8^ These benefits have led to a rapid expansion in the use of macrocycles across various therapeutic areas, including antibacterial, antiviral, and anticancer applications.

Macrocycles have been reported to target a wide range of biological molecules, including cell surface receptors,^9–11^ enzymes,^12–15^ protein-protein-interactions (PPIs),^16–19^ and protein-RNA interactions.^19–24^ Because PPIs play a fundamental role in cell signaling pathways related to initiation and progression of diseases, they are attractive targets for therapeutics.^25,26^ The three main classes of PPIs^27^ include a) interactions between short peptide motifs and small protein domains (e.g. SH2 and PDZ domains b) interactions between structured epitopes (e.g. single α-helix) and downstream binding partners and c) interactions between large binding surfaces formed by multiple distal and discontinuous motifs. Small molecules can be effective for targeting small, deep binding pockets such as the interactions between peptides and small protein domains (type a) but often are ineffective against large protein domain interactions (types b and c) due to the size of the binding surfaces and lack of specific interactions at defined sites. Macrocyclic peptides are well-suited for targeting these more challenging PPIs due to their ability to mimic a protein fold^3^ and present an extended surface area for binding.

While possessing many advantages for use in therapeutics, the *de novo* design of peptide macrocycles is technically challenging. Structural and computational approaches^28–30^ can be used to engineer molecules to disrupt PPIs but structural deconvolution and the success of computational prediction is not always guaranteed. As an alternative, shotgun-scanning^31^ and *in vitro* selection methods (e.g. display technologies) have been developed to make it possible to screen the high diversity of ligands required to identify effective inhibitors of PPIs.^32–34^ Recent progress in the discovery of synthetic macrocycles targeting protein-protein interactions through phage display,^34,35^ mRNA display,^36–38^ DNA-encoded libraries,^39,40^ as well as solid-phase-based diversity oriented libraries,^41^ has demonstrated the potential power of these approaches to produce effective inhibitors of PPIs.^42,43^ Furthermore, continuing advances in methods to incorporate non-natural amino acids and post-translational modifications using variable linkers in display methods have further expanded the chemical space available for macrocycles, enhancing both diversity and functionality.^44–47^ This includes multiple strategies that enable direct screening of peptide macrocycle libraries containing a reactive electrophile.^48^ This has enabled directed screens to identify covalent binding ligands for diverse targets. However, examples of covalent ligand selection using phage display or mRNA display have only focused on enzymatic targets with defined binding sites that contain a reactive nucleophile.

Covalent targeting strategies have experienced a resurgence in interest due to the recent success of several irreversible binding drugs. There are many benefits of the covalent binding mechanism, including reduced dosing frequency, simplified pharmacokinetic and pharmacodynamic considerations and reduced resistance generation.^49–51^ However, in order to avoid issues of off-target toxicity, it is essential to use electrophiles that have overall low reactivity and which require tight binding to the target of interest to facilitate covalent bond formation. To address this requirement for balancing reactivity with target selectivity, covalent electrophiles can be attached to larger complex molecules and proteins. In particular, recent examples incorporating a weakly reactive electrophile in existing protein-based^52–54^ or peptide-based^55–59^ ligands targeting large binding surfaces and PPIs have shown great promise as a way to achieve highly selective and irreversible binding to a target protein *in vivo*. However, approaches for the *de novo* development of covalent PPI inhibitors by direct screening of libraries of small, chemically accessible peptide structures has so far remained elusive.

We describe here a phage display platform integrating covalent chemistry with diverse libraries of macrocycles that can be screened using a combination of binary and ternary selection methods to identify optimal irreversible binding inhibitors of a target PPI. As a proof-of-concept, we targeted the interaction of SARS-CoV-2 spike protein with angiotensin II converting enzyme 2 (ACE2). Due to its importance for viral entry, replication and virus spread, this interaction has been the focus of structure-based design and screening efforts to identify potent and selective inhibitors.^60–64^ In addition, the use of weakly reactive SuFEx electrophiles on antibody scaffolds has already demonstrated the potential value of using covalency to enhance potency and duration of activity of inhibitors of the spike-ACE2 PPI.^53^ Therefore we reasoned that by combining a validated SuFEx electrophile with our phage screening platform to generate large libraries of cyclic peptides, we could create a platform for identification of small, chemically accessible covalent inhibitors of the spike-ACE2 PPI. By developing a screening workflow in which spike binders are enriched and then negative selection screening is performed using proteins that block the ACE2 binding site, we demonstrate that it is possible to identify compounds that covalently bind to spike resulting in irreversible anti-viral activity. This platform has not only identified a promising new class of anti-viral agents but also has the potential to be applied to identify covalent inhibitors of other therapeutically relevant PPIs.

## RESULTS AND DISCUSSIONS

### Development of A Selection Method to Screen for Protein-Protein Interaction Inhibitors

Given the inherent challenges of identifying inhibitors capable of blocking specific PPIs, we reasoned that using a covalent electrophile would help to facilitate the binding of relatively small ligands such as cyclic peptides at a site that may lack defined binding pockets. We therefore chose to use the phage display method that we have previously reported in which a chemical linker is used to cyclize peptides on the surface of the phage through the reaction with two engineered cysteine residues.^48^ This approach allows an electrophile of choice to be installed directly in the cyclic peptide scaffold through a highly efficient chemical reaction performed on the phage prior to selection. The resulting library of electrophile-containing cyclic peptides can then be screened for binding to the target and covalent binders can be enriched using stringent washing to remove reversibly bound peptides.

The major drawback of using target binding for selection rather than a functional assay, is that the final set of hits includes many non-productive binders that dilute the pool and make it difficult to focus on molecules that bind at a site of interest on the protein target. Therefore, to identify molecules capable of disruption of a specific PPI by binding at a known site on the protein target, it is crucial to develop strategies that can be used to exclude off-site binders during the analysis of the final data sets. To achieve this, we developed a strategy that uses a combination of binary (target + peptides) and ternary (target + blocker + peptides) screenings to filter and enrich for cyclic peptides that bind to a defined location on the target (Figure 1A). This approach uses traditional binary selections, referred to as positive selection hereafter, in which libraries are incubated with the target protein alone through all rounds of selection. In a second arm of selections after the first three rounds of positive selection, we use ternary selections, or counter-selection, in which the site of the target PPI is blocked with a high affinity or irreversible binding ligand prior to screening with the libraries. This approach allows the final data to be filtered to select site-specific binding hits that are both enriched throughout the rounds of positive selection but also lost in subsequent later rounds of negative selections.

**Figure 1.**
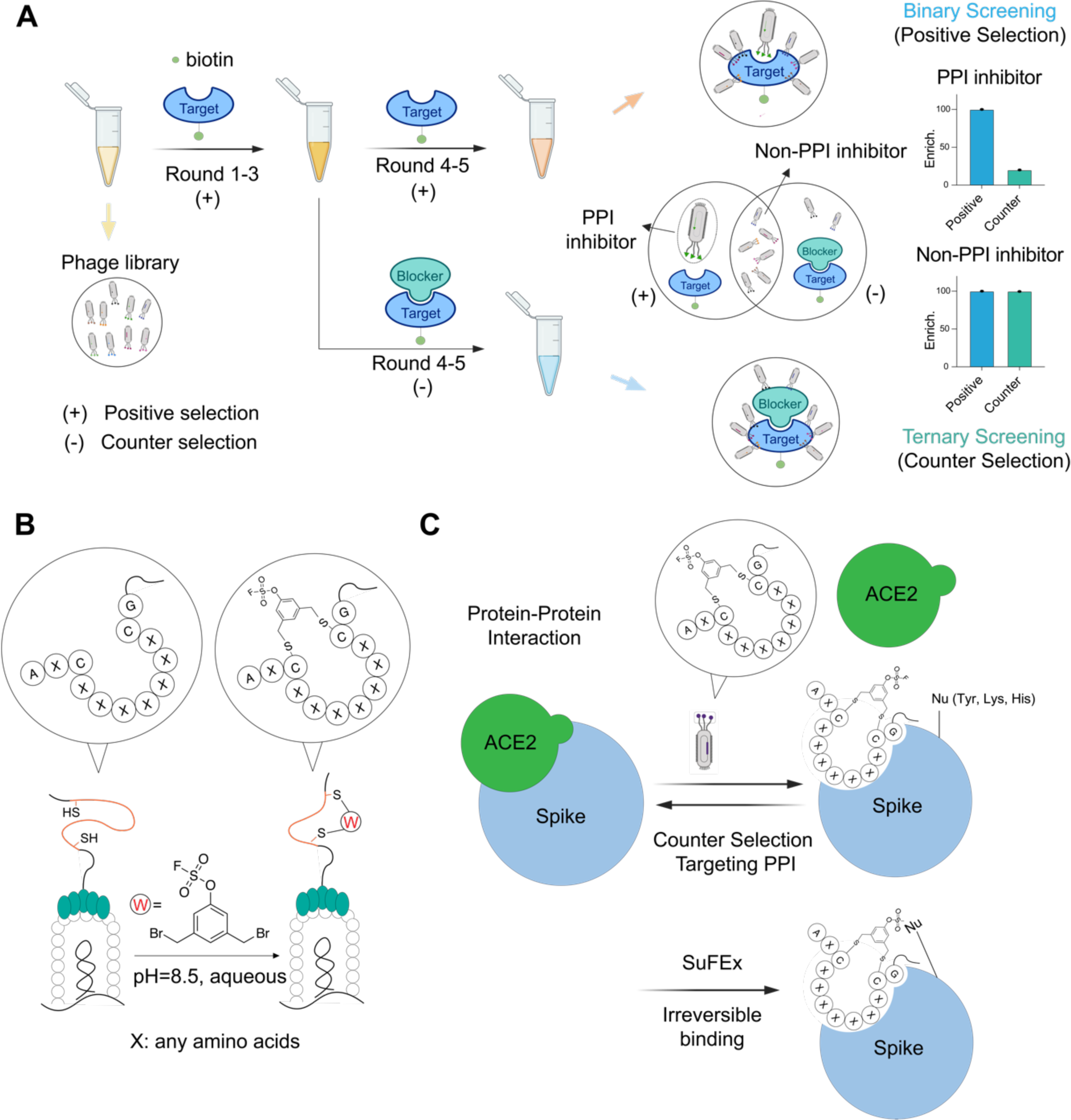
Selection strategy for covalent targeting protein-protein interaction using phage display. (A) General workflow of discovering inhibitors targeting protein-protein interactions using counter selections. Chemically modified phage libraries are panned against target protein (positive selection) for 3 rounds. The enriched phage pool after round 3 is divided equally and used for two additional rounds of either positive selection or negative selection in which the target is incubated with a known binder of the interface region. Sequencing results of elutes from both positive and negative selections in all rounds are compared to identify binders targeting protein-protein interaction interface. (B) Strategy of post-translational modification of phage displayed peptides. An aryl fluorosulfate warhead linker was used to chemically modify diverse peptides containing two conserved cysteines to generate a cyclic peptide–warhead library on the phage pIII surface. (C) Strategy for selecting inhibitors targeting protein-protein interaction of spike/ACE2. Cyclic peptide interface binders on phage compete with ACE2 for binding to the SARS-CoV-2 spike protein during the selection process. The fluorosulfate electrophile forms a covalent bond with a nucleophilic residue (e.g., Tyr, Lys or His) near the PPI interface resulting in irreversible inhibition of the native PPI.

### Design, Synthesis and Screening of Chemically Modified Phage Displayed Libraries Targeting the Spike-ACE2 Interaction

To validate our general selection approach, we used the SARS-CoV-2 spike protein as a target. This protein is an ideal candidate for several reasons. First, it has a defined PPI with its receptor target ACE2 that is required for host cell entry by the virus. Thus, selection for molecules that block this PPI have the potential to be novel inhibitors of viral infection. Second, prior efforts using phage display methods for this target have predominantly identified peptides that bind at sites outside the spike-ACE2 interface and therefore fail to have anti-viral activity.^65^ Therefore, the use of both a covalent binding electrophile and an optimized screening approach has the potential to overcome some of the existing pitfalls for identifying anti-viral agents that block spike-ACE2 interactions. Third, there are multiple antibodies and nano-bodies that have been identified that block the spike-ACE2 binding interaction, including several that have been engineered to contain covalent binding electrophiles.^53^ These molecules are ideal for use in our negative counter screens to help us to select for covalent cyclic peptides that bind at a site that will block subsequent ACE2 binding.

Given the pitfalls of using the N-terminal receptor binding domain (RBD) as a surrogate for the full-length spike protein in screens,^65^ we decided to use the full-length Omicron BA.2 variant spike trimer for our selections (Figure 1A, Supplementary Information). For our counter selections, we leveraged the native ACE2 receptor (Supplementary Information) as well as an engineered nanobody, Nb70 (56FFY)^53^, which covalently inhibits the ACE2-spike interaction and mimics the function of larger antibodies (Supplementary Information). This provided us with multiple negative selections using two ligands with distinct modes of binding to the spike protein.

To generate the library of covalent cyclic peptides displayed on the phage surface, we employed a previously described cyclization strategy using two conserved cysteine residues expressed between a variable sequence on the P3 surface protein.^48^ Given the abundance of tyrosine and lysine residues on protein surfaces and at protein-protein interaction (PPI) interfaces,^66,67^ we selected the fluorosulfate as the optimal reactive electrophile for chemical modification of the phage libraries (Figure 1B). This electrophile predominantly reacts with Tyr, Lys, and His residues through SuFEx chemistry,^68,69^ (Figure 1C). In addition, it is ideal for use in this application since it is highly stable *in vivo*, has overall low reactivity and has been used for targeting of PPIs by others.^53,70^ Therefore, we synthesized a linker in which the OSF electrophile was directly attached to the core aromatic ring of the linker to allow direct placement near the core of the cyclic peptide. We found that cyclization did not significantly impair phage infectivity, as confirmed by titer measurements (Figure S1), and high cyclization efficiency was verified through a pulldown assay (Figure S1).

For selection screening, we used five progressive rounds of phage panning beginning with three rounds of positive selections using full-length biotinylated BA.2 spike trimer followed by two parallel rounds of positive and counter-selections (Figure 2A, see Table S1 for selection details). We monitored overall phage recovery at each round of selection via titer measurements (Table S2). To eliminate non-binders and low-affinity binders, we progressively increased selection stringency by adjusting protein concentration, panning time, and washing conditions throughout the rounds. Based on the estimated nanomolar concentrations of the phage libraries (calculated from titer and copy numbers), we used 200 nM of spike trimer (∼5 µg) in round 1 to enhance recovery of bound phage, and then reduced the amount to 100 nM in round 2 and 25 nM in round 3. We also used a long 2.5-hour incubation period in rounds 1- 3 to promote formation of covalent labeling by the OSF electrophile and select as many possible covalent binders as possible. Total recovery of phage and fold change (sample/bead control) were relatively constant in the first two rounds, but increased significantly in round 3, suggesting enrichment for target binders (Table S2, Figure S2). In the later rounds 4 and 5, we further increased selection stringencies by decreasing the incubation time with the libraries to 1.5 hours and reduced the concentration of the spike protein to 10 nM in the final round 5 of selections. As an additional step to reduce non-specific binding, we used 0.1% BSA in the selection buffer and alternated between neutravidin and streptavidin beads as negative controls to minimize the selection of bead-specific binders. We also used stringent washing conditions for all rounds of selections. This included a non-denaturing wash (PBS with 0.1% Tween-20), a more stringent wash (PBS with 1% Tween-20), and finally a denaturing wash (5 M guanidinium in PBS) to eliminate low-affinity or non-covalent binders. After the initial three rounds of positive selection, we equally divided the eluted phage for positive and counter-selections in rounds 4 and 5 (Figure 2A). For these counter-selections, we pre-incubated the spike protein with ACE2 or the covalent nanobody Nb70 (56FFY) before panning with the phage libraries. We used a Mesoscale Discovery (MSD) assay to confirm single digit nanomolar binding affinity for both ACE2 protein and the Nb70 (56FFY) nanobody for full-length BA.2 spike trimer consistent with previously reported activities for these proteins (Figure S3).^53,71^ Furthermore, the potency of binding of only the nanobody increased with increasing incubation time confirming its covalent mode of binding. Based on the relatively slow kinetics of Nb70 (56FFY) binding to the spike protein, we pre-incubated the nanobody and spike for 4 hours prior to panning. We also confirmed by BLI binding assays that this 4-hour of incubation did not alter the ability of spike to bind ACE2 (Figure S4), ensuring protein stability during the selection process.

**Figure 2.**
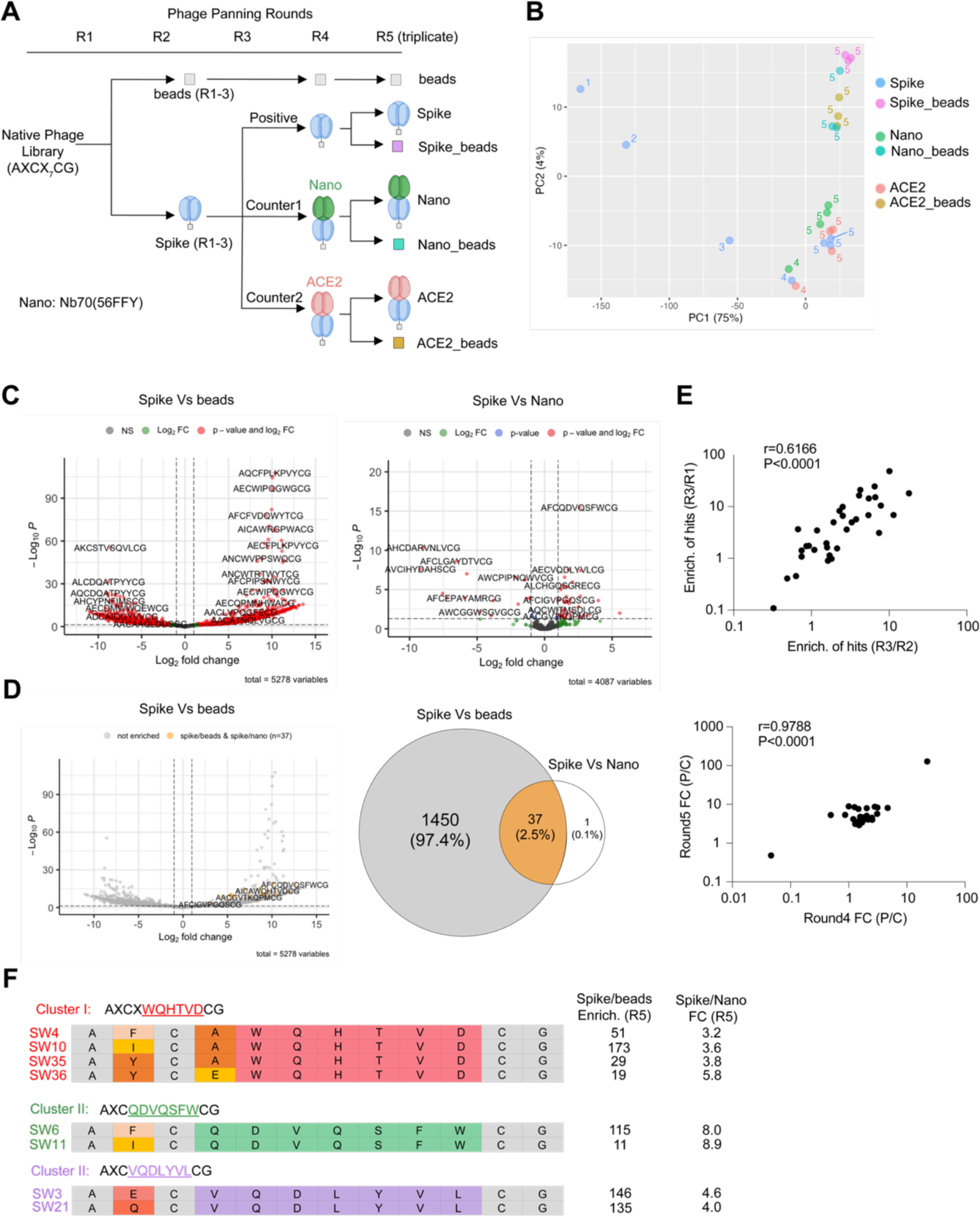
Bioinformatic analysis for deconvolution of hits targeting protein-protein interactions using counter selections. (A) Workflow of five rounds of phage panning. Names of seven samples in round 5 were annotated. (B) Principal component analysis of selection samples in five rounds. Samples are colored coded and selection rounds are coded by numbers. (C) Left: Volcano plot of enriched hits comparing positive selection to empty beads selection in round 5. Right: Volcano plot of enriched hits comparing positive selection to counter negative selection using covalent nanobody Nb70 (56FFY) in round 5. All peptides were detected at least once in at least one of the three replicates in round 5. (Cutoff: Fold change >2, p-value < 0.05) (D) Left: re-color all the peptides in grey for the left volcano plot of (C), highlighting the 38 enriched hits in yellow from right panel of (C). Right: Venn diagram of overlapping enriched hits between both left and right panels of (C), showing number/percentage for each intersection. (Significance cutoff: Fold change >2, FDR < 5%) (E) Enrichment correlation of 37 hits in the first three rounds. Enrichment of hits between different rounds were calculated using the avgCPM numbers of each hit in those rounds. Enrichment of hits in round 3 over round 2 was plotted against the enrichment of hits in round 3 over round 1. Pearson score r = 0.6166. p-value < 0.0001. Fold changes of hit enrichment in positive selection and negative selection. Fold changes of hits were calculated using the enrichment of those hits in positive selection and counter negative selection using nanobody in both round 4 and round 5. Fold changes in two rounds were plotted for correlation. Pearson score r = 0.9788. p-value < 0.0001. (F) Family cluster analysis of hits. Three clusters sharing conserved motifs are shown. Conserved binding motifs were highlighted using the same color for the same cluster members. Variable amino acids were coded by color. Conserved amino acids are colored grey. Enrichment of each hit calculated using the positive selection and empty beads selection in round 5 is shown along with fold changes of enrichment in positive selection and counter negative selection (nanobody) in round 5.

### Bioinformatic Analysis to Deconvolute Hits that Disrupt the Protein-Protein Interaction

#### Pre-analysis of NGS Data Quality

After sequencing of the eluted samples from all five rounds of selection, we evaluated the data quality before proceeding with analysis. The phage library was designed with a semi-random peptide sequences AXCX7CG displayed on the PIII region of the filamentous phage fd,^72^ as previously described.^48^ To assess the quality of peptide sequences in the NGS data, we counted all detected peptides across all datasets, focusing on those with the correct 12-amino acid length. After selecting all peptides containing 12 amino acids within the semi-random sequences as we designed the library format as AXCX7CG, the majority of peptides displayed the expected amino acids at the constant positions, and those with mutations in these positions were discarded. A sequence logo analysis confirmed the expected amino acid composition after truncation (Figure S5). In total, we detected approximately 542,000 unique peptide species across all samples. The phage library has a diversity of 1x10⁹, with around 50 copies of each specific peptide in the initial library. Peptides from round 1 had relatively low counts, mostly ranging from 10 to 1,000 (Figure S6). As panning rounds progressed, the distribution of peptide counts skewed towards higher values, reflecting the enrichment of binders and elimination of non-binders under the selective pressure of multiple rounds. The number of unique peptide species was tracked across all five rounds (Figure S6), showing a gradual decrease in both beads and spike groups, with only ∼10,000 unique peptides remaining in round 5. After filtering out low-count peptides, 644 peptides (with counts >10) remained in round 5 for the spike group (Figure S6).

To assess the overlap of peptides across samples, we analyzed how many peptides detected in round 5 were exclusive to one sample or shared among multiple samples (Figure S7). This analysis is crucial to determine if the enrichment of peptides in each sample is meaningful and to guide downstream enrichment analysis using DESeq2^73^ (see Methods in Supplementary Information), a method for conducting differential analysis of count data, which utilizes shrinkage estimation for dispersion and fold changes to enhance the clarity of the estimates.

When considering all detected peptides, most were unique to a single sample (Figure S7A). When restricting the analysis to peptides consistently detected across replicates (i.e., >1 count in all replicates of each sample), 45% were found exclusively in the beads sample (Figure S7B). Notably, only 160 peptides were reliably detected across all four samples (positive selection: spike only; counter selection 1: spike + nanobody; counter selection 2: spike + ACE2; beads selection: empty beads only). From the Venn diagrams of detected peptide distribution among positive and two counter selections (Figure S7C), abundant intersection (45%) shared by three groups suggests large amounts of potential non-PPI binders were excluded with the counter selection.

#### Principal Component Analysis

To assess the data quality and reproducibility of our panning process, we performed principal component analysis of the sequencing data in all five rounds (Figure 2B, Figure S8A). Our PCA analysis shows the expected evolution of selected peptides throughout panning rounds: while panning against beads causes separation along the second principal component (which explains only 4% of variance in the data, Figure S8A), panning against Spike had the most notable impact on the population of selected peptides, causing gradual shifts away from beads and along the first principal component (which explains ∼93% of variance), with the first three panning rounds having the largest impact. To zoom in and evaluate round 4 and round 5 data sets, we generated a new PCA (Figure 2B) excluding round 1 to 5 beads data, because they are dominant data outliers in the full PCA analysis (Figure S8A). Due to the maturation of phage binder selection profile in the later rounds, round 4 and round 5 showed mild changes. In addition, because the majority of the hits in the pools of selection are non-PPI binders, samples of positive selections and counter selections in round 4 and 5 did not show too much variance in the PCA analysis map, as expected (Figure 2B). We performed three replicates for each selection group in round 5, which also showed good reproducibility based on their tight clusters in the PCA.

#### Comparison of Positive and Counter Selection

In the 5th round of selection, we analyzed seven groups of selection datasets: negative control (selection using empty beads starting from round 1, named as “beads”), positive selection (selection using spike only in all 5 rounds, named as “spike”), round 5 beads control for positive selection (empty beads selection using round 4 positive selection elute, named as “spike_beads”), counter selection 1 (selection using spike and nanobody, named as ”nano”), round 5 beads control for counter selection 1 (empty beads selection using round 4 counter selection 1 elute, named as “nano_beads”), counter selection 2 (selection using spike and ACE2, named as “ACE2”), round 5 beads control for counter selection 2 (empty beads selection using round 4 counter selection 2 elute, named as “ACE2_beads”). Using DESeq2 analysis, we calculated the differential enrichment of hits for three key comparisons: positive selection vs. empty beads, positive selection vs. counter selection 1, and positive selection vs. counter selection 2 (Figure 2C, Figure S8B-D). The correlation matrix indicates robust correlation between the positive and counter selections (Figure S9), while a few significantly enriched peptides could be found in the positive compared to negative volcano plots. Given that ACE2 and the nanobody blocked the protein-protein interface, the peptides significantly enriched in positive over counter selections are considered PPI inhibitors (Figure 1A). The high correlation implies that the majority of the enriched hits in positive selection over beads are non-PPI inhibitors, since PPI interfaces occupy only a small portion of the entire protein surface. Comparison of positive selection and counter selection 1 using covalent nanobody identified 38 hits as potential interface binder (Figure 2C). One of these 38 peptides was also significantly enriched when comparing positive and counter selection 2 using ACE2, using the same cutoff adjustment (Figure S8B). Of the 38 enriched hits, 37 were also enriched in positive over beads, but they accounted for just 2.5 % of the total peptides enriched in the positive selection (Figure 2D). Notably, these 38 hits were not the most highly enriched but were distributed across various fold changes and p-values (Figure 2D). This highlights the stark difference between using simple binary positive selections compared to ternary screenings with defined negative controls, and explains why results from affinity selections often do not correlate well with functional screening results.

#### Enrichment Correlation of Hits in Sequencing

We used normalized DNA copy numbers (avgCPM) to calculate enrichment between rounds (R2/R1, R3/R1, and R3/R2) for the 38 enriched hits from counter selection 1. Enrichment correlation using Pearson score (r=0.6041, P=0.0002; r=0.6166, P=0.0001) indicated that enrichment in each round is highly correlated through panning propagation for those hits (Figure 2E). For rounds 4 and 5, fold changes between positive and counter selections were calculated using DNA copy numbers. High correlation (r=0.9788, P<0.0001) between two rounds indicated that enriched samples in round 4 with higher fold changes (positive/counter) enrich further with even higher fold changes in round 5, identifying those candidates as promising inhibitors for protein-protein interactions (Figure 2E).

#### Family Cluster Analysis of Hits

Among the full set of 37 hits, we identified three distinct sequence clusters (Figure 2F), suggesting that defined groups of compounds with similar structures emerged from the selection process. Enrichment profiles of those hits within each cluster based on the DNA copy numbers (AvgCPM) for each round, exhibited similar enrichment patterns across the five rounds of selection (Figure S10), reflecting comparable behaviors of hits sharing the same binding motif, and success of the *in vitro* evolution. The significant enrichment of these hits in the round 5 positive selection over the empty beads (Figure 2F), coupled with high fold changes in positive selection compared to the counter selection, strongly supports their potential as potent inhibitors of the Spike/ACE2 protein-protein interaction.

### Functional Validation of Phage Display Hits

#### Synthesis, Testing and Identification of Top Screening Hits

We selected the top 37 peptide sequences identified from bioinformatic analysis for synthesis using rink amide resin following solid phase peptide synthesis using a semi-automated peptide synthesizer. After synthesizing the crude linear peptides and performing global deprotection with TFA, we formed the final cyclic peptides using the dibromo-aryl fluorosulfate linker used in the phage panning (Figure 3A, detailed in synthesis scheme 1). The crude products exhibited high yields and purity (50-95%, Table S3), primarily consisting of the desired final product with minor residual side products, such as disulfide-mediated cyclized compounds without the warhead linker, as confirmed by LC/MS.

**Figure 3.**
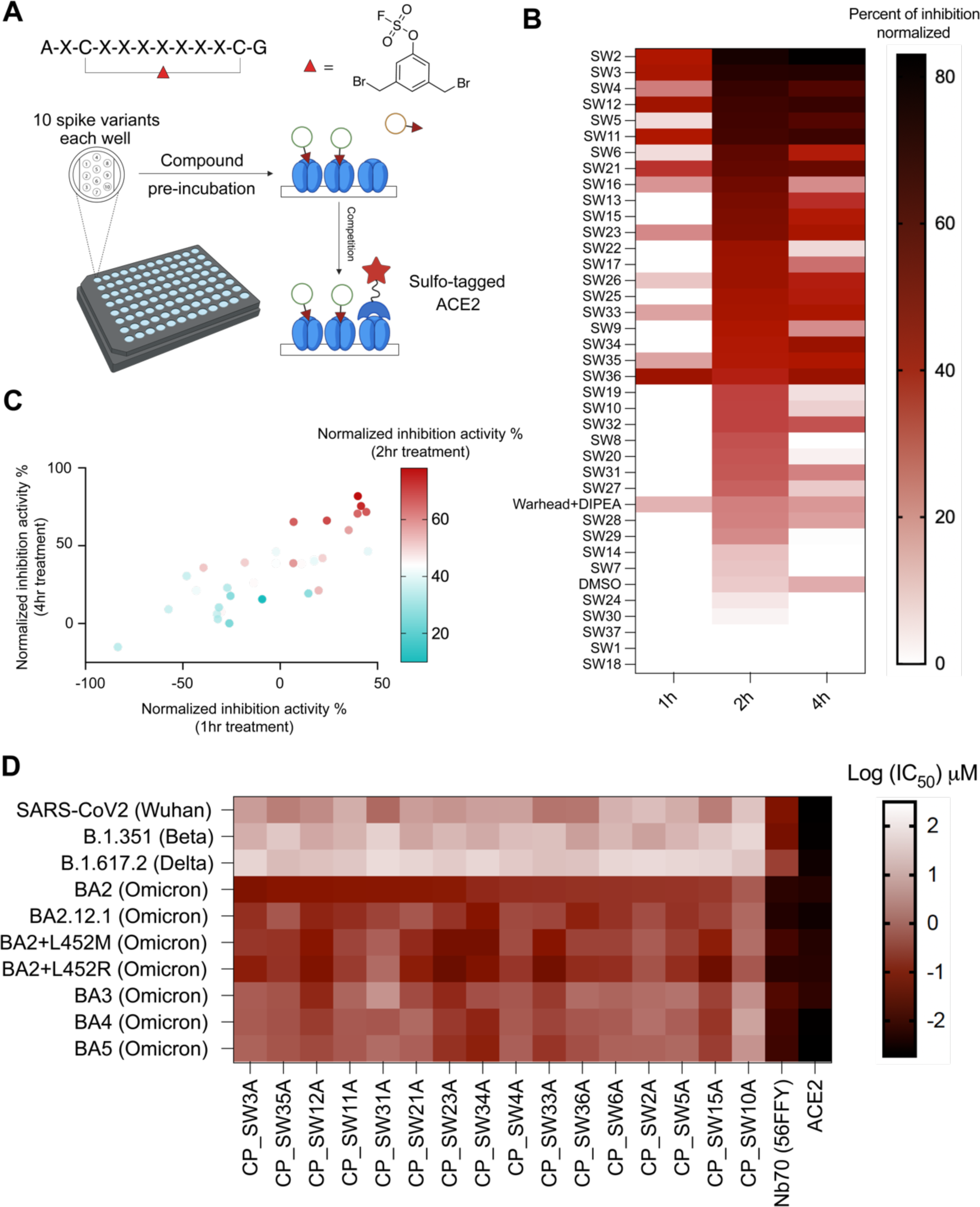
Activity evaluation and cluster analysis of hits. (A). Time-dependent activity evaluation of hits using Meso Scale Discovery (MSD) electrochemiluminescence assays. The top 37 high-yield crude cyclic peptides (AXCXXXXXXXCX, X=any amino acid in hit sequences) were dissolved in DMSO as compound stocks. Stock compounds were pre-incubated at 10 µM in well pre-coated with 10 different SARS-CoV-2 full length spike variants with different incubation times (1/2/4 hrs) on an MSD plate, before performing a competition binding assay with Sulfo-tagged ACE2. the dibromomethyl aryl fluorosulfate linker and DIPEA (used for cyclization) was used as a negative control, and DMSO was used as a solvent background control. (B) Compound inhibition activity using 10 µM to Omicron BA2 variant with three different incubation times (1/2/4 hrs) was normalized using a calibration reagent provided with the MSD assay and plotted as a heatmap using Graphpad prism 10. The order of compounds in the heatmap was arranged based on the 2 hr incubation time point inhibition data, from the highest activity to the lowest. (C) Correlation of the inhibition activity of hits at three different time points (1/2/4 hrs) were plotted using a multi-variable correlation map in Graphpad Prism 10. Normalized inhibition activity at 2hr incubation was color coded as a third variable in the map. (1h/4h: r= 0.8136, p= 1.5027e-008; 1h/2h: r=0.6105, p= 0.0002; 2h/4h: r= 0.8241, p= 6.7976e-009). The 1hr and 4hr data are shown as normalized inhibition activity. The 2hr activity data is shown as color coded. (D) Full dose inhibition activities of annotated purified compounds, nanobody (Nb70 (56FFY)) and ACE2 to 10 SARS-CoV2 spike variants in MSD assay were shown. IC50 values after 4 hours of compound incubation to disrupt the interaction between annotated spike and Sulfo-ACE2 using MSD assay are plotted as a heatmap.

Encouraged by previous studies demonstrating the use of crude compounds for rapid screening, with subsequent validation of purified compounds after selection of hits,^74–76^ and considering the high yield and purity of crude products, we decided to directly measure activities of cyclic peptides in the biochemical assays without further purification. Crude peptides were lyophilized to dryness and dissolved in DMSO to prepare 10 mM stock solutions, based on resin loading capacity. We then utilized the Meso Scale Discovery (MSD) multi-spot assay system to directly measure the ability of each peptide to block binding of ACE2 to the spike protein. This assay is a well-established high-throughput screening platform in which each well of a 96-well plate is coated with up to 10 spike variants and binding of a sulfonated ACE2 protein is measured after pretreatment with a test agent (Figure 3A). This assay enabled the simultaneous quantitative measurement of the binding of each of our hit peptides to 10 spike variant antigens, including the Omicron BA2 used for selection (Figure 3A).

Compounds were pre-incubated with spike antigens on the plates, followed by the addition of human ACE2 conjugated with MSD SULFO-TAG to assess competition and electroluminescence detection. Compounds were tested at two concentrations (1 µM and 10 µM) for three increasing incubation times (1 hr, 2 hrs, and 4 hrs) at 37°C. The activity of each compound was normalized to a standard calibrator provided in the assay kit. Remarkably, 31 out of 37 hits exhibited time-dependent activity (2 hrs vs. 4 hrs) to Omicron BA2 spike at the lower 1 µM concentration (Figure S11), while 26 out of 37 hits showed time-dependent activity across all time points (1 hr, 2 hrs, and 4 hrs) at 10 µM, suggesting that they are likely covalent inhibitors (Figure 3B). Correlation analysis of the MSD assay results revealed significant correlations in both dose and time dependencies (Figure 3C), supporting the statistical robustness of using high-yield crude compounds. Based on both time-dependent MSD activity assays and sequence cluster analysis, 17 hits with either potent time-dependent activity or belonging to a cluster, were selected for resynthesis introducing a clickable handle and purification for further evaluations.

#### Full-dose Activity Evaluation and Direct Labeling of Spike Trimer Protein using Purified Peptide Hits

Those 17 hits were resynthesized with a propargyl glycine substitution for the original N-terminal alanine and purified for MSD assay and protein labeling. Full-dose characterization of the purified hits in MSD assays revealed that CP_SW3A, and several other promising hits, had high potency to Omicron BA2 (IC50 = 100-500 nM) (Figure 3D, Figure S12). In addition, activities to three other Omicron BA2 sublineages (BA2.12.1, BA2+L452M, BA2+L452R) showed comparable submicromolar activities, while significant loss of activities (5-10 fold) were observed for BA3, BA4 and BA5 Omicron lineages, and even more loss of activities (100-1000 fold) for previous viral strains (Wuhan strain, B1.351 (Beta strain) and B.1.167.2 (Delta strain). As positive controls, ACE2 and Nb70 (56FFY) showed potent nanomolar activities to all viral lineages. For protein labeling, compounds were directly incubated with spike protein before undergoing a CuAAC click reaction with an azide-TAMRA for fluorescence quantification. Among the eight labeled hits from the three clusters, the extent of labeling correlated with enrichment for cluster II and cluster III (Figure 2E and Figure S13). Out of the 17 compounds tested, CP_SW3A (cluster III) and CP_SW35A (cluster I) demonstrated the highest gel labeling intensity (Figure S13) and strong activity in MSD assays (Figure S12). Combining enrichment (positive selection), fold changes (positive/counter), MSD assay results, and gel labeling efficiency, CP_SW3A (Figure 4A) emerged as the optimal lead molecule for further validation.

**Figure 4.**
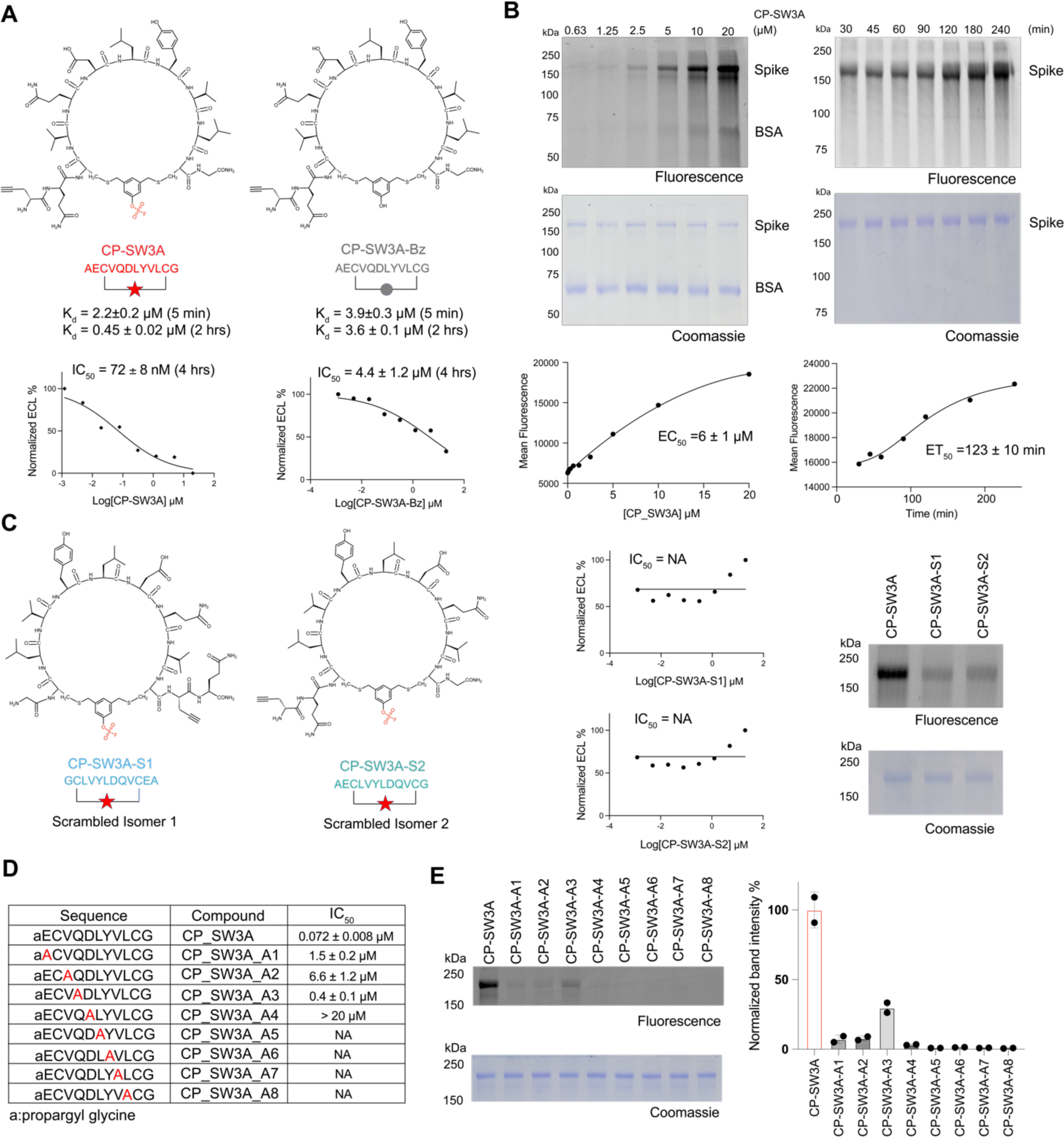
**Biochemical validation of optimal hit CP-SW3A**. (A) Structures of the optimal hit CP-SW3A and the negative control compound CP-SW3A-Bz lacking the electrophile. The time dependent dissociation equilibrium constant Kd of the two compounds characterized using BLI at two different association time points (t=5min and 2 hr) are shown. IC50 values for inhibition activities after 4 hours of compound incubation to disrupt the interaction between Omicron BA2 spike and ACE2 using MSD assay are shown below. (B) Dose- and time-dependent labeling of the full-length spike Omicron BA2 variant by CP-SW3A. The alkyne labeled peptide was added at the indicated concentrations for 2 hrs or at the concentrations of 10 µM for the indicated times followed by CLICK labeling with azide-TAMRA. Labeled protein was resolved by SDS-PAGE followed by scanning of the gel for fluorescence (C) Structure of two scrambled isomers of CP-SW3A. Sequence of isomer S1 is fully reversed compared to the parental CP-SW3A, and sequence of isomer S2 is only reversed for the variable regions between the two cysteines. Inhibition activities of the two isomers using MSD assay are shown. NA: Not available. Labeling of the two isomers, along with the parental compound CP-SW3A, to spike protein is shown. For labeling, 0.25 µM spike was incubated with 10 µM compounds in PBS, 37 °C, for 2 hours. Labeled mixtures were clicked with azide-TAMRA via CuAAC click reaction, followed by fluorescence scanning and coomassie staining. (D) Evaluation of structure dependent labeling activity of CP-SW3A using positional alanine scanning mutants. Alanine was scanned across the variable amino acids except the first alkyne and last glycine, to yield 8 alanine mutants (annotated as A1-A8 in table). IC50 values for inhibition activities after 4 hours of compound pre-incubation to disrupt the interaction between Omicron BA2 spike and ACE2 using MSD assay are shown below. (E) Each mutant and the parental peptide was incubated at 10 µM with 0.3 µM spike in PBS for 2hr at 37C. Labeled mixtures were clicked with azide-TAMRA via CuAAC click reaction, followed by fluorescence scanning and coomassie staining. Band intensities in fluorescence imaging were quantified by two technical replicates using ImageJ. Quantification data was normalized to the parental molecule CP-SW3A.

To again corroborate the feasibility of using high-yield crude compounds in the primary screening to accelerate DMTA cycle, we assessed whether crude compounds display similar patterns to their purified counterparts by comparing protein labeling between crude and purified forms (Figure S14). Due to the comparable peptide yields, five of six compounds demonstrated similar labeling patterns between crude and purified forms, except for CP_SW6A, due to its lower crude yield.

#### Spike Labeling and Structure Activity Relationship Analysis of Top Hit Cyclic Peptide

To assess the dose-dependency and kinetics of labeling, we performed direct labeling studies of the spike protein with, CP_SW3A, over a range of concentrations and time points. CP_SW3A, along with other promising hits (Figure S15), demonstrated clear concentration and time-dependent labeling of the spike protein, achieving an EC50 of 6 ± 1 µM for concentration- dependent labeling and around two hours for half-maximal protein labeling (Figure 4B). In addition, the probe showed only faint labeling of BSA at the highest concentration tested suggesting that the covalent reaction was mediated by specific binding interaction between the cyclic peptide and the spike target protein.

To further confirm that the labeling of spike by CP_SW3A was due to reaction of the electrophile and binding by specific interactions between the probe and the target, we synthesized two control compounds lacking the warhead, CP_SW3A_Bz (Figure 4A) and CP_SW3A_DS (Figure S16), along with two scrambled isomer peptides, CP_SW3A_S1 and CP_SW3A_S2 (Figure 4C). Using a Biolayer Interferometry (BLI) assay, we determined that CP_SW3A and CP_SW3A_Bz exhibited comparable apparent binding affinities during a 5-minute incubation in the association step (Figure 4A, Figure S17). However, the dissociation constant (Kd) of only CP_SW3A improved five-fold to 0.45 µM due to slower off-rate, when the association time was extended to two hours, confirming that the electrophile confers time-dependent covalent inhibition of the target (Figure S17).

We next compared the activities of the two scrambled isomers, CP_SW3A_S1 and CP_SW3A_S2, to the parental compound using in-gel fluorescence labeling and MSD assays. Both scrambled isomers exhibited dramatically reduced labeling and loss of inhibition activities (Figure 4C) compared to the parent compound. This data demonstrates the importance of the peptide sequence of CP-SW3A for its activity against spike. To further assess the contribution of each residue in the CP_SW3A sequence to its binding activity, we performed an alanine scan (Figure. 4D). All mutants demonstrated a significant loss of inhibition activities (Figure 4D, Figure S18), as well as protein labeling (Figure 4E), with only limited residual activity observed when alanine was used at the A1-3 positions. This suggests that the activity of CP_SW3A is highly dependent on the specific sequence makeup of the selected cyclic peptide.

### Irreversibly Anti-viral Activity of the Top Covalent Spike Inhibitor

While our result in biochemical assays confirmed that our top hit binds covalently and irreversibly to the spike protein, in order for this binding to be of potential value for a therapeutic, we needed to demonstrate that the peptide blocks productive infection of a host cell. To evaluate the ability of our top compound CP_SW3A (along with other promising compounds) to disrupting the Spike-ACE2 protein-protein interaction in a cellular model, we conducted infection assays using authentic SARS-CoV-2 originally isolated from nasopharyngeal swabs of infected patients (Figure 5A). We also used Sotrovimab, a monoclonal antibody targeting the spike protein of SARS-CoV-2, as a positive control. In the assay, we pre-incubated the peptides with the virus (Omicron variants BA2, BA1, and BA5; Figure 5A, Figure S19) at 37°C for 1 hour. We tested compounds at a maximum concentration of 150 µM with three-fold serial dilutions to achieve five concentrations. The peptide-virus mixtures were then added to VeroE6 cells and after 24 hrs, viral replication was quantified by measuring the amount of SARS-CoV-2 nucleocapsid protein in infected cells. We determined the effective concentration (EC50) of the inhibitors against cell infection using non-linear regression dose response analysis. The top lead molecule, CP_SW3A, demonstrated single-digit micromolar inhibition activity against the two closely related variants, with EC50 values of 6 µM for Omicron BA2 and 5 µM for Omicron BA1 (Figure 5B). In contrast, it exhibited substantially weaker activity against Omicron BA5 (EC50 = 47 µM). This decreased efficacy is likely due to the high mutation rate which leads to significant changes in the spike protein which impact our inhibitor binding This is consistent with the reduced activity of the control antibody Sotrovimab (EC50 = 7 µM) in the BA5 antiviral assay (Figure 5B). The negative control scrambled compounds CP_SW3A_S1, CP_SW3A_S2, and CP_SW3A_Bz displayed only weak background activity against all three variants consistent with their inability to covalently target spike (Figures 5B).

**Figure 5.**
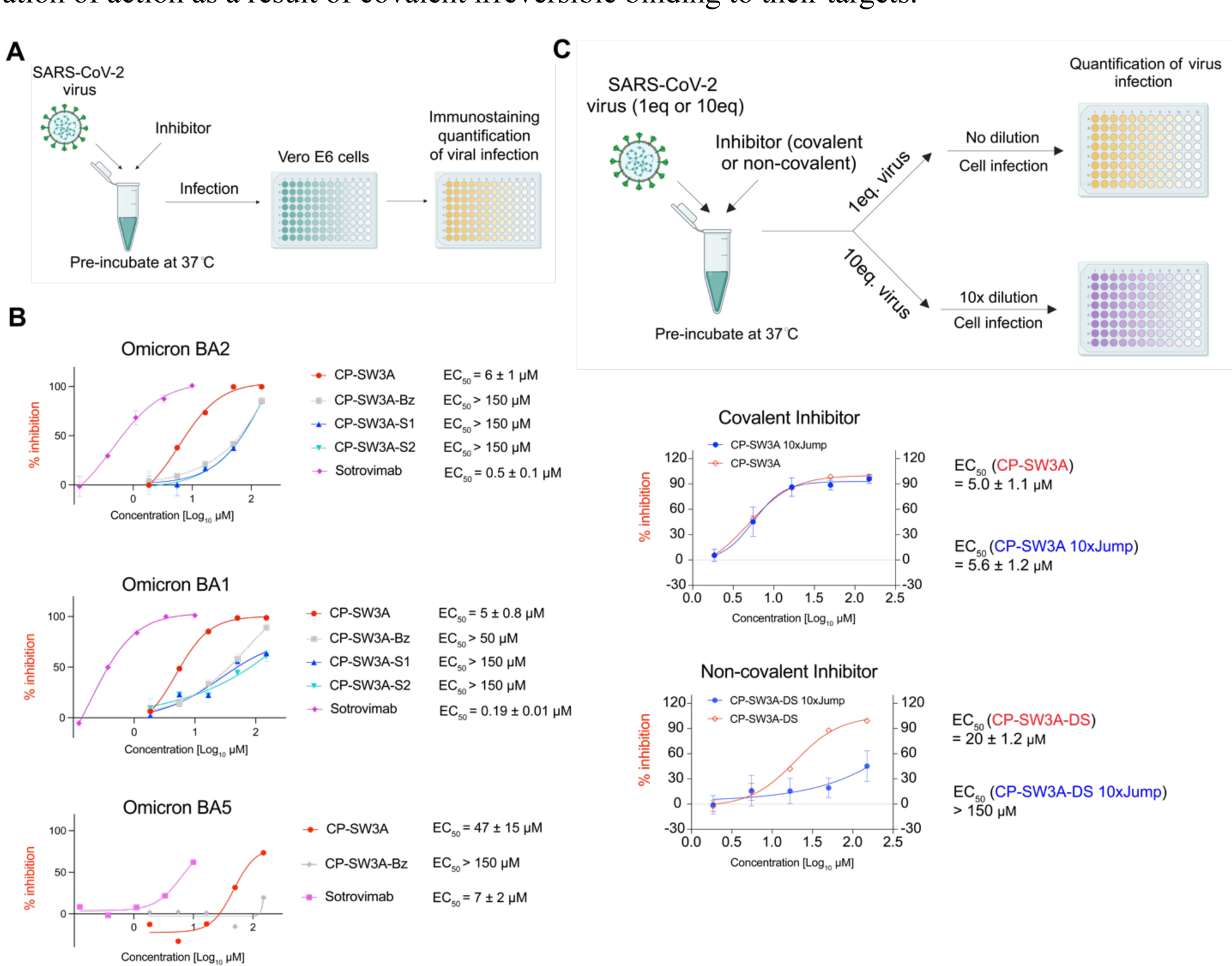
Cellular antiviral activity of hits. (A) scheme of viral infection assay. SARS-CoV-2 particles, collected from clinical samples and amplified in cell culture, were incubated with compounds (or controls: sotrovimab as positive control, and DMSO as blank) in PBS for 1 hour in 37°C before infecting VeroE6 cells. Viral replication was measured at 24h post infection by immunostaining of the viral nucleocapsid in infected cells. (B) Dose response neutralization activity of annotated compounds and positive control sotrovimab in the antiviral assay using Omicron BA2, BA1 and BA5 variants. Values were normalized to solvent control (1% DMSO) with infection (100% infection) and without infection (0% infection, assay background). EC50 values were calculated using non-linear regression dose response analysis by GraphPad Prism version 8.4.3 (GraphPad Software, San Diego CA, USA). (C) scheme of jump dilution in the viral infection assay. Test compounds (covalent compound: CP-SW3A; non-covalent compound control: CP-SW3A-DS) were prepared in PBS containing 3% DMSO and incubated with either 1eq (5x10^4^ TCID50) or 10eq (5x10^5^ TCID50) of SARS-CoV-2 omicron BA.2 for 1 hour at 37°C. The 1eq viral particle sample was directly used for cell infection. The 10 eq sample mixture was diluted 10-fold with 3% DMSO containing PBS and 1eq of diluted viral particles was inoculated to VeroE6 cells. The virus replication was determined at 24h post infection using immunostaining of viral nucleocapsid. Values were normalized to solvent control (1% DMSO) with infection (100% infection) and without infection (0% infection, assay background). EC50 values were calculated using non-linear regression dose response analysis by GraphPad Prism version 8.4.3 (GraphPad Software, San Diego CA, USA).

Given that one of the most significant benefits of a covalent inhibitor is the lack of an off rate which translates into a long-duration of activity, we wanted to confirm that the activity of our selected inhibitor was in fact irreversible. Therefore, we performed an assay in which virus was pre-incubated with the active CP_SW3A inhibitor as well as with the weakly active CP-SW3A-DS control that lacks the electrophilic warhead (Figure 5C), followed by a rapid (jump) dilution to mimic a washout, prior to infection of the cells. We pre-incubated viral particles at a ratio of 10 eq and 1 eq with the same concentrations of CP_SW3A or the no-warhead control compound CP_SW3A-DS for 1 hour at 37°C. The 10 eq virus-compound mixture was then diluted to 1 eq before cell infection, while the 1 eq mixture was used directly for infection. CP_SW3A exhibited no substantial loss of inhibitory potency even after the 10-fold dilution, whereas CP_SW3A-DS demonstrated a dramatic loss of activity after dilution. These results confirm the covalent binding mode of CP_SW3A in the cellular viral infection assay and suggest that this approach could be used to generate anti-viral inhibitors that have prolonged duration of action as a result of covalent irreversible binding to their targets.

## CONCLUSION

In this study, we employed a directed phage display selection method with a combination of positive selection and counter selections to identify hits that bind specifically to a target protein at a site that disrupts a protein-protein interaction. We validated selected hits through structural, biochemical, and cellular assays to assess their ability to bind the spike protein and disrupt its interaction with ACE2. The covalent electrophile effectively prolonged the apparent binding efficacy, as validated by Biolayer Interferometry (BLI) assays. In authentic antiviral assays, our top hit exhibited significant antiviral activity, supporting its potential to target the spike protein on real viral particles and block entry via ACE2 interaction.

Bioinformatic analysis established in this work can potentially serve as a standard pipeline for hit deconvolution of hits of any other PPIs using genetically encoded library screenings. Considering the relative non-competitive environment in the initial round of selections and the selection pressure for covalent hits, we also evaluated if there is correlation between selection enrichment (on phage) and synthesized hit activity (off phage). Protein labeling consists of two major steps. The first step involves reversible binding between the compound and the protein, which brings the warhead into close proximity with nucleophiles located near the binding pocket. The second step is an irreversible process in which the electrophile forms a covalent bond with its nucleophile target (such as Tyr, Lys, or His). The outcomes of direct protein labeling reflect the combinatorial effects of these two steps, paralleling the positive selection panning process. In the first three rounds of positive selection, the protein concentration (ranging from 25 to 200 nM) is significantly higher than the total phage library concentration (less than 2 nM), often exceeding it by 10-100 fold. This relatively low stringency creates a non-competitive environment for phage-displayed binders, facilitating maximal recovery of all potential protein binders with varying affinities. The recovery or enrichment of individual binders is primarily driven by the affinity between the protein and each binder displayed on the phage, which is a result of both the reversible binding step and the irreversible step that forms the covalent bond. Given the similar nature of direct protein labeling and the initial three rounds of positive selections, we conducted a correlation analysis of hits from both processes. This analysis yielded a correlation score of r = 0.5787 (P = 0.0327), indicating a fair degree of correlation between the two datasets.

While our strategy effectively identifies specific binders targeting protein-protein interactions, optimizations of platform design and validations can be further elaborated. Our selection platform can potentially generate both orthosteric and allosteric inhibitors for protein-protein interactions. In order to further elaborate the platform to generate hits for both site specificity and functionality, site-directed mutagenesis can be considered as an additional counter selection to separate orthosteric and allosteric inhibitors. In our later stage validation, we had challenges in localizing a robust and quantifiable inhibitor binding site on the spike protein using mass spectrometry as also reported in another study.^65^ Due to the highly heterogenous post translational modifications (glycosylation) on the spike protein, it is challenging to generate sufficient peptide fragment coverage after trypsin digestion for mass spec analysis. We also used PNGase treatment to remove the glycosylation before analysis, but due to other modifications, we were unable to obtain substantial peptide coverage by mass spectrometry. This may be partially due to the fact that the cyclic peptide is a large adduct to the protein and may be partially digested under strong proteolysis conditions used to generate peptides. Alternatively, the cyclic peptide modification may also preclude ionization of the labeled peptide. Regardless of these challenges in molecularly characterizing the binding site, our data using time-dependent MSD assays, in-gel fluorescence protein labeling, time-dependent BLI assays, and jump dilution in cellular infection assays all strongly support a covalent mechanism of binding.

Our selection and validation models serve as a pilot study for discovering covalent macrocyclic inhibitors of protein-protein interactions using phage display. This selection platform integrates covalent chemistry, macrocycles, binary/ternary screenings, and phage display, creating a versatile toolbox for functional screenings targeting PPIs. It can be envisioned that other genetically encoded libraries or high throughput screening platforms can leverage this concept for covalent PPI inhibitor discovery as well. This strategy could also potentially be adapted to discover probes for intracellular targets, leveraging available tools for designing cell-penetrating peptides. Furthermore, binary/ternary screenings used in this work were also previously used for the discovery of bifunctional molecules, such as molecular glues and PROTACs,^77^ while the discrepancy is the variable ways to compare and analyze the selection data. Molecular glue and PROTAC molecules can stabilize the ternary complex with expected enhanced enrichment in ternary screening, while PPI inhibitors destabilize the ternary complex, thus showing decreased enrichment. Notably, Sulfur(VI) Fluoride Exchange (SuFEx) chemistry has been used successfully for targeting of established binding molecules (antibodies and nanobodies), but it has until now not been used for selection screens with phage or other display methods. The SuFEx electrophile enables macrocycles to covalently bind and, importantly, disrupt protein function. Macrocycles can mimic the action of targeting large binding domains, akin to antibodies, while offering advantages such as stability, ease of accessibility and distribution, and low production costs.

## ASSOCIATED CONTENT

### Supporting Information

Supplementary tables and figures, materials and methods for biological evaluation, detailed synthesis procedures and compound characterization, LC-MS, and NMR spectra (PDF)

### Corresponding Author

\***Matthew Bogyo** − Department of Pathology, Stanford University School of Medicine, Stanford, California 94305, United States; Department of Microbiology and Immunology, Stanford University School of Medicine, Stanford, California94305, United States;

### Funding Sources

This work was funded by an NIH grants R21 AI191151 (to M.B.) and R01 CA258300 (to L.W.). This work was supported by the Project “Virological and immunological determinants of COVID-19 pathogenesis – lessons to get prepared for future pandemics (KA1-Co-02 “COVIPA”)”, a grant from the Helmholtz Association’s Initiative and Networking Fund and by the Deutsche Forschungsgemeinschaft (DFG, German Research Foundation) – Projektnummer 240245660 – SFB 1129 (to R.B.). This work was supported by the National Science Foundation Graduate Research Fellowship under grant no. DGE- 1656518, Stanford ChEM-H O’Leary-Thiry Graduate Fellowship, NIH Stanford Graduate Training Program in Biotechnology T32GM141819, International Alliance for Cancer Early Detection (ACED) Graduate Fellowship (to F.F.).

### Notes

The authors declare no competing financial interest.

## ACKNOWLEDGMENT

The authors thank Dr. Daniel Abegg and Dr. Alex Adibekian for their support and helpful discussions. The authors thank Prof. Peter Kim at Stanford University for support. The authors thank Dr. Marta Barniol-Xicota for helpful discussions. The authors thank Dr. David Veesler, Dr. Young-Jun Park and Pooja Dnyaneshwar Bandawane for their discussions. The authors thank Brandon Lam for help in MSD data export.

