## Supporting Information for "Identification of Covalent Cyclic Peptide Inhibitors Targeting Protein-Protein Interactions Using Phage Display"

**Table of Contents** **Page**

1. **Supplementary Tables and Figures**

**Table S1**: Selection conditions for five rounds of selection S4

**Table S2**: Phage recovery and fold change using titer measurement S5

**Figure S1**: Phage infectivity and modification yield upon linker treatment S6

**Figure S2**: Phage recovery and fold changes in five rounds of selection S7

**Figure S3**: Evaluating binding of ACE2 and nanobody Nb70 (56FFY) to full length Omicron BA2 spike using MSD assay S8

**Figure S4**: Evaluating Omicron BA2 spike protein activity using biolayer interferometry assay S9

**Figure S5**: Quality control of peptide lengths and trimming to 12 amino acids S10

**Figure S6**: Abundance metrics and detected peptide counts of five rounds S11

**Figure S7**: Venn Diagrams of detected peptide distribution in round 5 samples S12

**Figure S8**: PCA analysis and enrichment of hits in counter selections over empty beads S13

**Figure S9**: Correlation matrix of positive over counter selections S14

**Figure S10**: Overlay of enriched hits in positive selection and positive/counter selection 1 S15

**Figure S11:** Time and dose-dependent activities of 37 compounds for Omicron BA2 spike using MSD plate S16

**Figure S12:** Full-dose inhibition activities of purified hits to Omicron BA2 spike using MSD assay S17

**Figure S13:** Direct protein labeling of all 17 hits to spike S18

**Figure S14:** Comparing protein labeling using purified hits or crude hits S19

**Figure S15:** Dose- and time-dependent labeling of purified hits S20

**Figure S16:** Spike labeling comparing CP-SW3A and negative control compound CP-SW3A-DS S21

**Figure S17:** Evaluating Kd of CP-SW3A and CP-SW3A-Bz using BLI assay S22

**Figure S18:** Inhibition activities of isomers, no warhead control, and alanine mutants of CP-SW3A in MSD assay S23

**Figure S19:** Evaluating compounds' activities in authentic antiviral assays S24

**2. Materials and Methods** **S25**

General reagents for phage display S25

Phage panning S27

Phage infectivity measurement (titers) S28

Post-translational chemical modification S28

Phage titers/recovery/fold change S29

Positive and counter selections S29

Library DNA Preparation for NGS S30

Analysis of Phage Display sequencing data S30

Bioinformatic pipeline for counter selection data analysis S30

Mesoscale Discovery (MSD) assay S58

Biolayer Interferometry S59

In-gel fluorescence protein labeling of cyclic peptides to spike S59

Determination of neutralization activity using authentic anti-viral assay S60

Antiviral cellular jump dilution assay S60

**3. Chemical Synthesis** **S62**

General methods S62

**Scheme S1**: Synthesis of DBr-FS linker (dibromo aryl fluorosulfate) S63

Synthesis of DBr-FS linker (dibromo aryl fluorosulfate) S63

Synthesis of macrocyclic hits S65

**4. Compound table** **S67**

**5. LCMS of linker DBr-FS and cyclic chemical probes** **S70**

**6. NMR Spectra** **S103**

**7. References** **S104**

| | Round 1 (+) | Round 2 (+) | Round 3 (+) | Round 4 (+/-) | Round 5 (+/-) |
|---|---|---|---|---|---|
| Spike Trimer used in selection | 5 ug | 2.5 ug | 1.25 ug | 1.25 ug | 0.5 ug |
| Spike Protein Conc. | 200 nM | 100 nM | 25 nM | 25 nM | 10 nM |
| Binding time | 2.5 h | 2.5 h | 2.5 h | 1.25 h | 1.25 h |
| Beads | Neutravidin | Streptavidin | Neutravidin | Streptavidin | Neutravidin |
| BSA | 0.1% | 0.1% | 0.1% | 0.5% | 1% |
| Washing<br>3x PBS (0.1% Tween, 1%BSA)<br>3x PBS (1% Tween)<br>3x 5M guanidinium | 5min (x9) | 5min (x9) | 5min (x9) | 10min (x9) | 20min (x9) |

**Table S1**: Selection conditions for five rounds of phage panning, including spike protein used, incubation time for the phage library with selection protein (Binding time), beads type, BSA amount, and washing conditions (buffers and washing cycles).

| Panning Rounds | Groups | pre-panning titer | after-panning titer | phage recovery (%) | recovery fold change (sample/ctrl) |
|---|---|---|---|---|---|
| 1 | Spike | 1.35E+12 | 6.50E+07 | 0.0048 | 25 |
| | Ctrl (Beads only) | 1.35E+12 | 2.60E+06 | 0.00019 | |
| 2 | Spike | 9.33E+11 | 6.60E+07 | 0.0071 | 21 |
| | Ctrl (Beads only) | 4.33E+11 | 1.47E+06 | 0.00034 | |
| 3 | Spike | 1.1E+12 | 1.93E+09 | 0.18 | 245 |
| | Ctrl (Beads only) | 1.133E+12 | 8.12E+06 | 0.00072 | |
| 4 | Spike | 1.67*10^11 | 7.8*10^7 | 0.047 | 2930 |
| | Ctrl (Beads only) | 1.2*10^12 | 2.66*10^5 | 0.000022 | |
| | Spike+ACE2 | 1.67*10^11 | 7.4*10^7 | 0.044 | 2780 |
| | Spike+Nb70 | 1.67*10^11 | 7.6*10^7 | 0.046 | 2082 |
| 5 | Spike | 0.47 *10^10 | 3.47*10^7 | 0.74 | 753 |
| | Spike (B) | 0.47 *10^10 | 2.66*10^5 | 0.00566 | 5.7 |
| | Ctrl (Beads only) | 4.7*10^11 | 4.6*10^6 | 0.00098 | |
| | Spike+ACE2 | 0.616*10^10 | 5.4*10^7 | 0.88 | 898 |
| | Spike+ACE2 (B) | 0.616*10^10 | 1.72*10^5 | 0.0028 | 2.9 |
| | Spike+Nb70 | 0.667*10^10 | 2.34*10^7 | 0.81 | 827 |
| | Spike+Nb70 (B) | 0.667*10^10 | 1.334*10^4 | 0.0026 | 2.7 |

Note: for round 5, Spike (B) means the spike output library from round 4 is incubated with only beads at round 5.      B: Beads

**Table S2**: Phage recovery and fold changes of five rounds of panning using titer measurement. Fold changes were calculated using the phage recovery of samples (positive and counter selections), normalized to control (beads only).

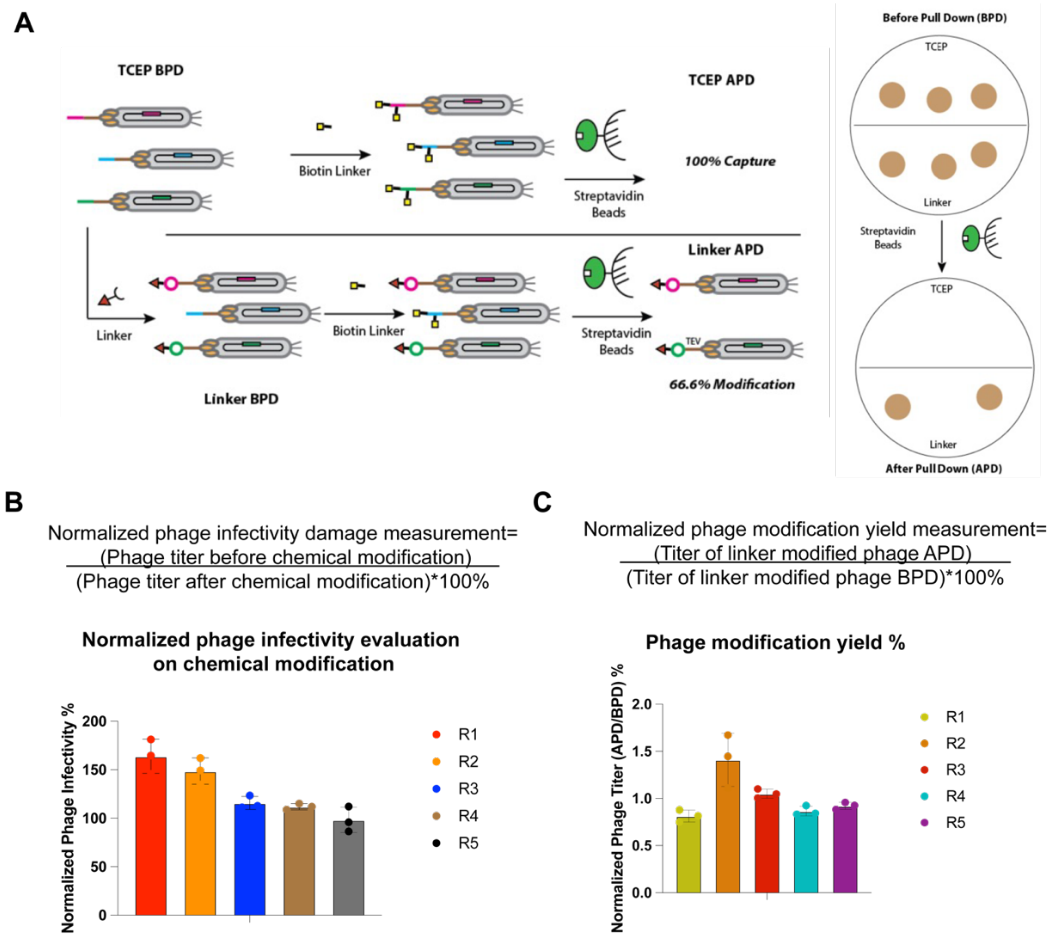


**Figure S1**: Phage infectivity and modification yield upon linker treatment (A). Representative scheme of phage titer measurement upon linker treatment and methods for measuring linker modification yield. Phage libraries were splitted, with one group treated with TCEP only ("TCEP BPD"), and another group treated with TCEP followed up with linker treatment ("Linker BPD") (details see methods). For two groups above, they were splitted again, with one subgroup remained untreated and another subgroup treated with biotin polyethyleneoxide iodoacetamide (IADB), followed up with pulldown using streptavidin beads (named as "TCEP APD", "Linker APD"). Phage titers of all four groups were measured counting colony numbers on plates (see methods for details). (B) Phage infectivity damage evaluation on chemical modification was quantified by phage titer after chemical modification (Linker BPD group) normalized to phage titer before chemical modification (TCEP BPD group). (C) Phage chemical modification yield was quantified by phage titer upon linker modification after pulldown (Linker APD) normalized to phage titer upon linker modification before pulldown (Linker BPD).

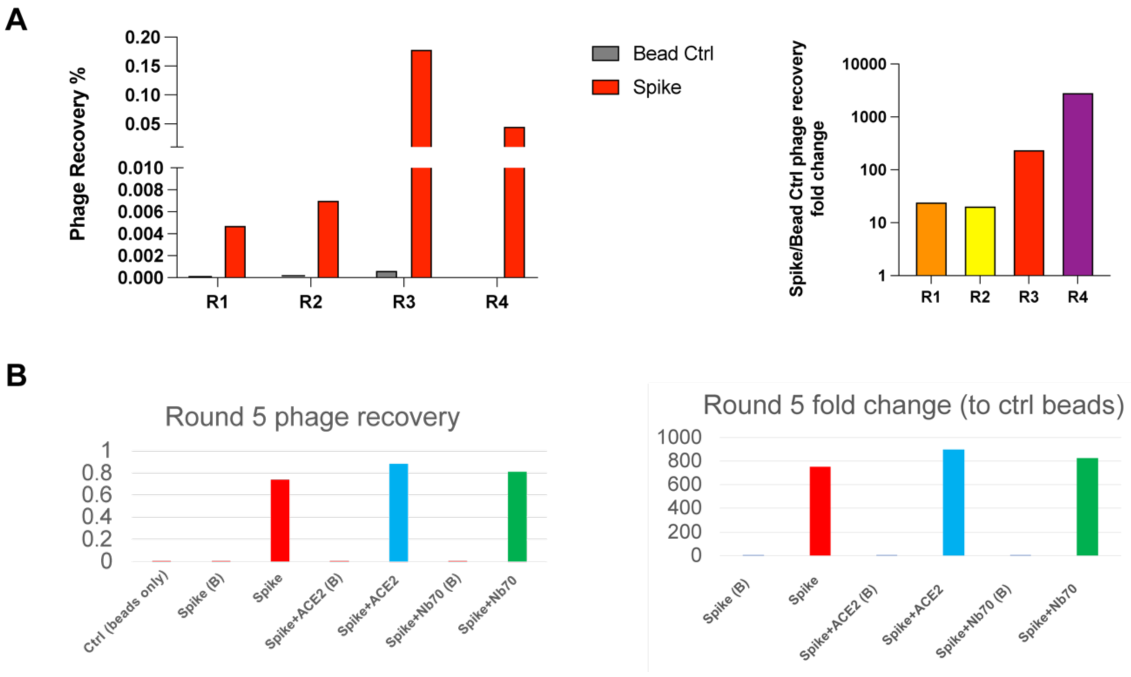


**Figure S2**: Phage recovery and fold changes in five rounds of selection (A) Recovery of Phage was quantified by the phage titers of post-panning samples (beads ctrl and spike) normalized to pre-panning library titers for each round from round 1 to round 4. Fold changes (spike/beads) were calculated by using the titers of spike and beads ctrl in that round. (B) Phage recovery and fold changes were plotted for 7 samples in round 5 panning (control (beads only throughout all five rounds), beads panning using the elute from spike panning in round 4 (Spike (B)), positive panning using the elute from spike panning in round 4 (Spike), beads panning using the elute from counter negative 2 panning using ACE2 in round 4 (Spike+ACE2 (B)), counter negative panning using ACE2 using the elute from counter negative 2 panning using ACE2 in round 4 (Spike+ACE2), beads panning using the elute from counter negative 1 panning using nanobody Nb70(56FFY) in round 4 (Spike+Nb70 (B)), counter negative panning using Nb70 using the elute from counter negative 1 panning using Nb70 in round 4 (Spike+Nb70).

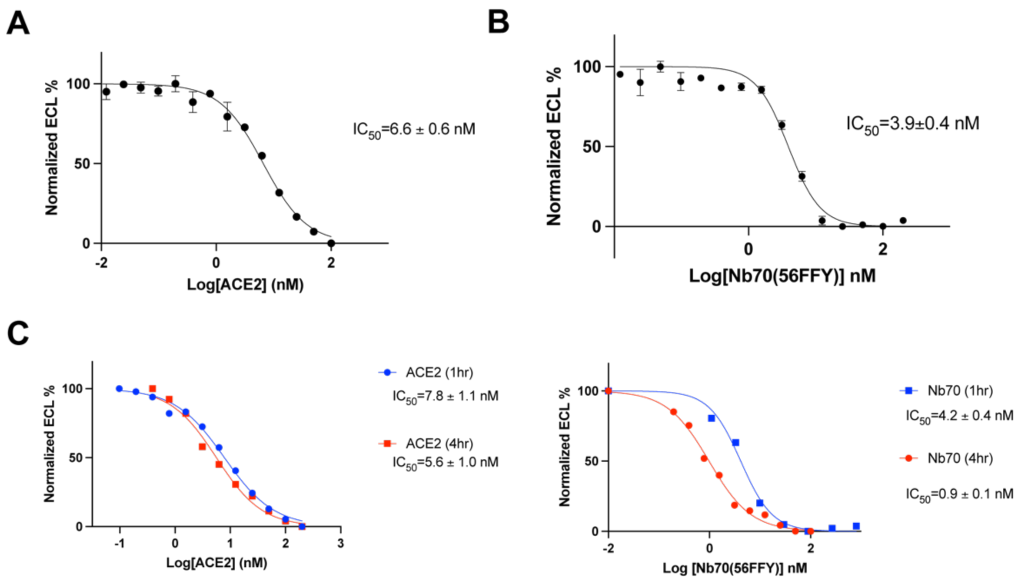


**Figure S3**: Evaluating binding of ACE2 and nanobody Nb70 (56FFY) to full length Omicron BA2 spike using MSD assay. Plots of ACE2 (A) and nanobody Nb70 (56FFY) (B) in full-dose binding to Omicron BA2 spike protein in competition against a Sulfo-tagged ACE2, using an MSD plate. Percent inhibition was calculated and normalized to a DMSO control, with points and bars representing mean ± standard deviation (n=3). $IC_{50}$ for each are shown at inset. (C) Time-dependent binding of ACE2 and Nb70 (56FFY) to Omicron BA2 spike. In MSD assay, ACE2 and Nb70 (56FFY) were added to pre-coated BA2 spike on the plate for either 1hr or 4hr incubation before adding a Sulfo-tag ACE2 in competition. Percent inhibition was calculated and normalized to a DMSO control, with points and bars representing mean ± standard deviation (n=3). $IC_{50}$ for each are shown at inset.

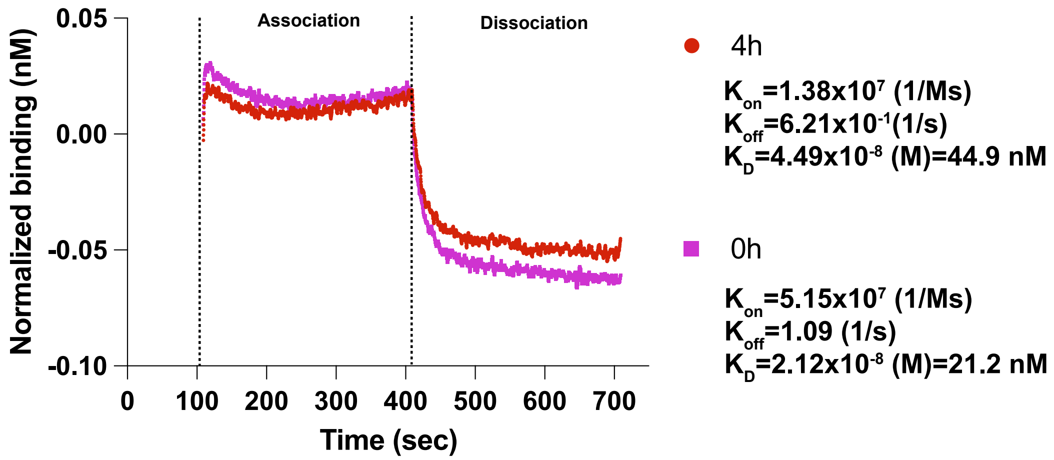


**Figure S4**: Evaluating Omicron BA2 spike protein activity using biolayer interferometry assay. Reactions were run on Octet RED96. Biotinylated Omicron BA2 spike (spike protein incubated in 37C for 4hours (4h:red) or 0hours (0h:purple) before BLI) was loaded onto the streptavidin biosensors. A buffer equilibrium step was added after that. ACE2 was then loaded for binding association step followed up with a dissociation step. For data presentation in plots, association and dissociation binding curves were normalized and substracted from the buffer equilibrium step before association. Binding curves were fit in Octet System Data Analysis Software version 9.0.0.15 using a 1:2 bivalent model for IgGs to determine apparent Kd using Kon and Koff. A control well with loaded antigen but that was associated in a well containing Octet buffer was used as a baseline subtraction for data analysis. Averages of Kd values from at least two independent experiments are reported.

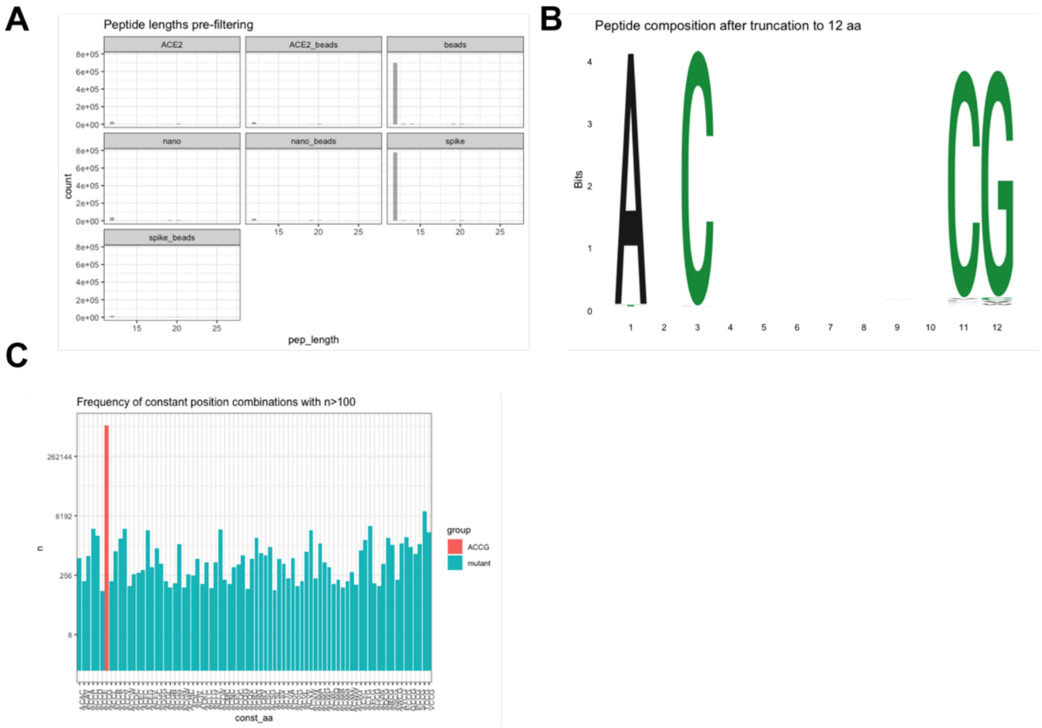


**Figure S5**: Quality control of peptide lengths and trimming to 12 amino acids. (A) Pre-filtering peptide lengths for 7 samples in round 5 elutes from NGS (B) Sequence logo showing expected aa composition AXCXXXXXXXCG after truncation. (C) Frequency of constant position combinations with n>100. The vast majority of peptides has the right amino acids at constant positions. There are ~542000 peptide species detected in total across all samples.

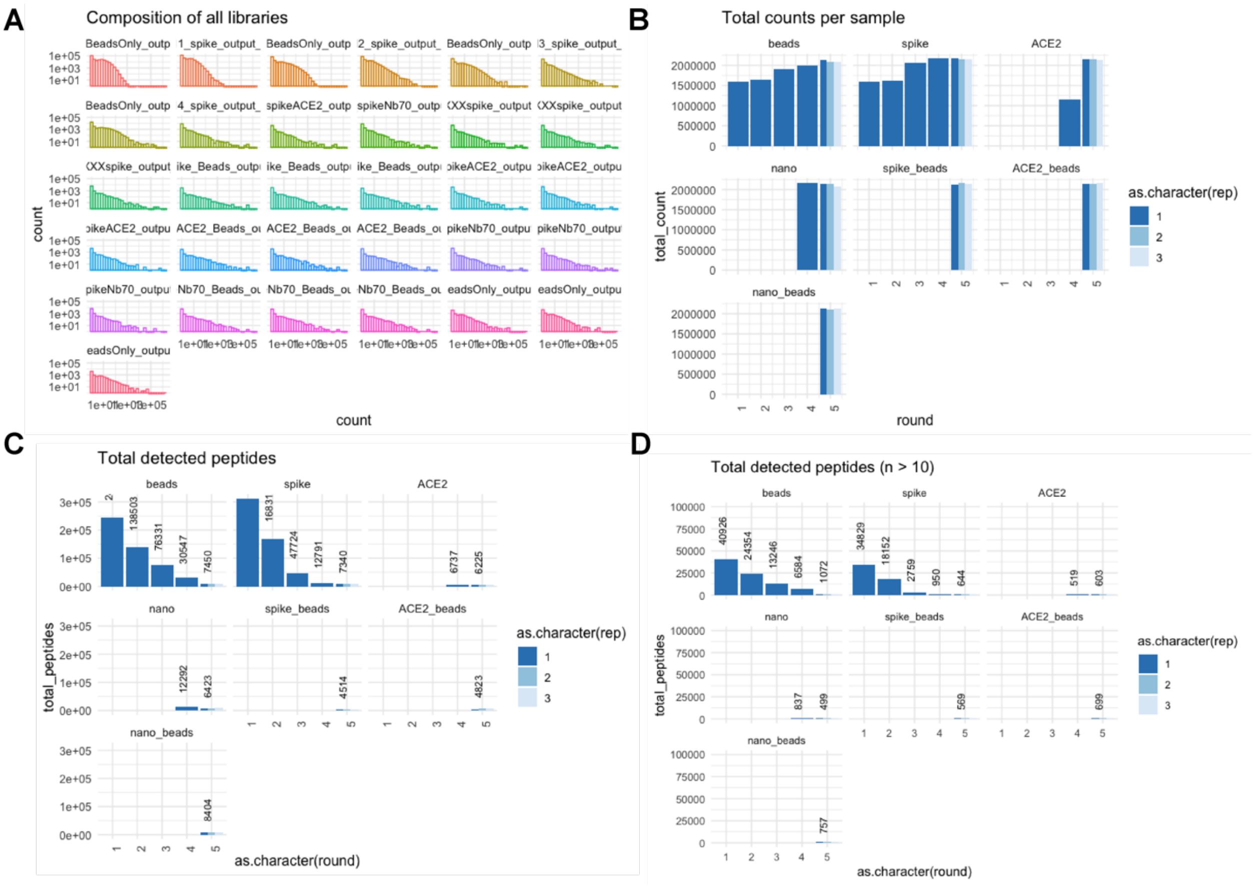


**Figure S6**: Abundance metrics and detected peptide counts of five rounds. (A) Composition of all samples in five panning rounds. X axis represents peptide counts. Y axis represents how many peptides had that count. (B) NGS total reads for each sample in five rounds. For all samples 2 million reads were assigned equally for each sample for sequencing. (C) Total detected peptides (n>=1) for each sample in five rounds. (D) Total detected peptides (n>10) for each sample in five rounds.

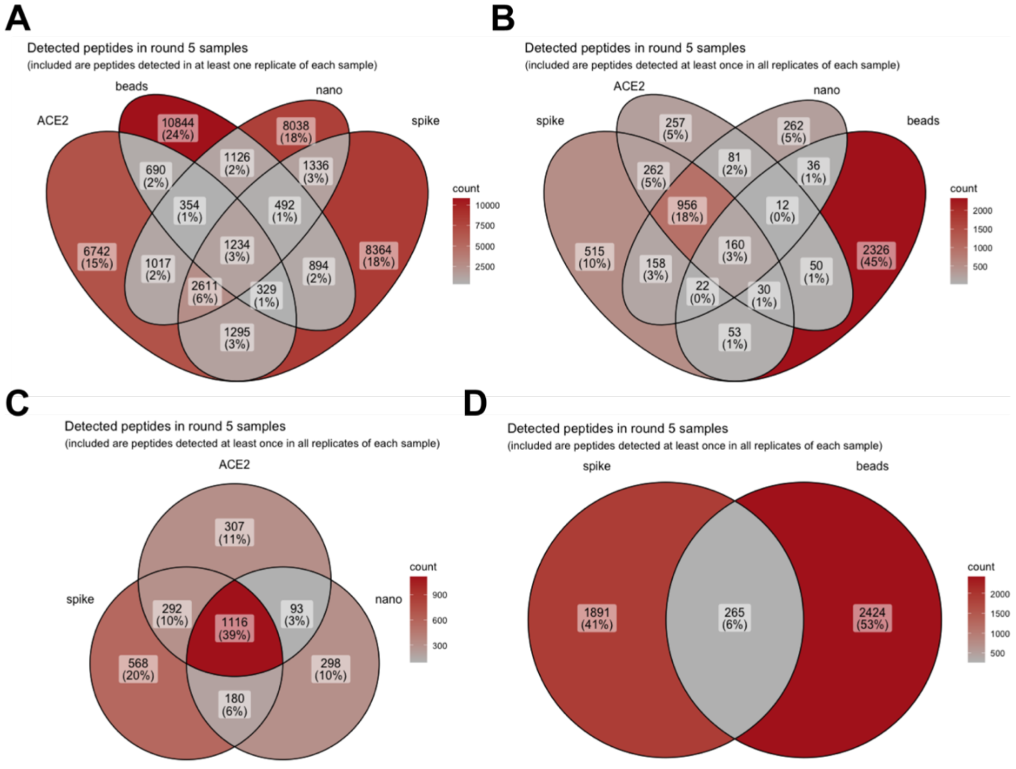


**Figure S7**: Venn Diagrams of detected peptide distribution in round 5 samples including spike (positive panning), nano (counter negative panning 1), ACE2 (counter negative panning 2), and beads (empty beads panning). (A) Detected peptides in at least one replicate of each sample, from the technical triplicates of each sample in round 5 (B) Detected peptides in at least once in all replicates of each sample, from the technical triplicates of each sample in round 5. (C) Detected peptides at least once in all replicates of each sample (spike/ACE2/nanobody) (D) Detected peptides at least once in all replicates of each sample (spike/beads)

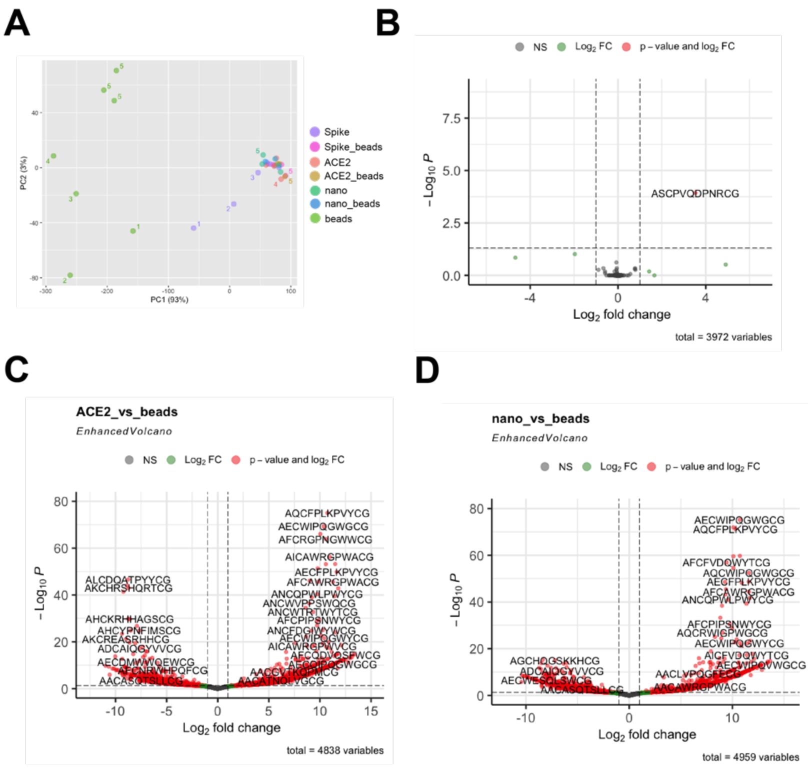


**Figure S8**: (A) Principal component analysis of selection samples, including beads selection, in five rounds. Samples are colored coded and selection rounds are coded by numbers. (B) Volcano plot of enriched hits comparing positive selection to counter negative selection using ACE2 in round 5 (C) enrichment of hits in counter selection 2 (ACE2 as blocker) over empty beads selection (D) enrichment of hits in counter selection 1 (Nb70(56FFY) as blocker) over empty beads selection.

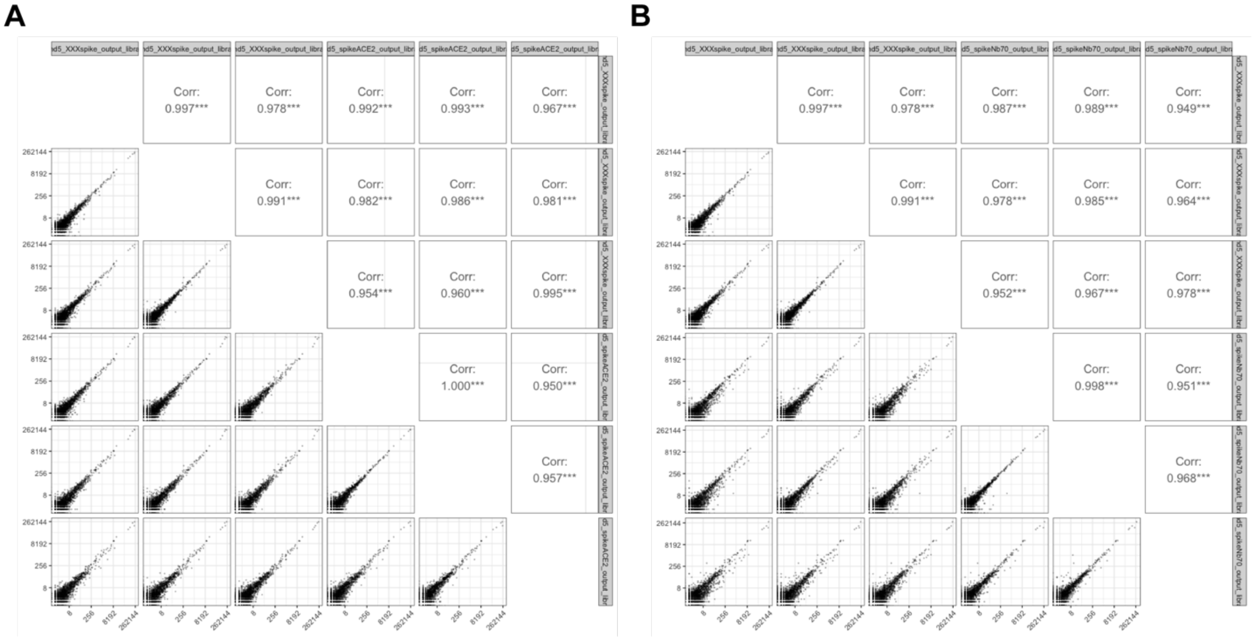

**Figure S9**: Correlation matrix of positive over counter selection 2 (A), and positive over counter selection 1 (B).

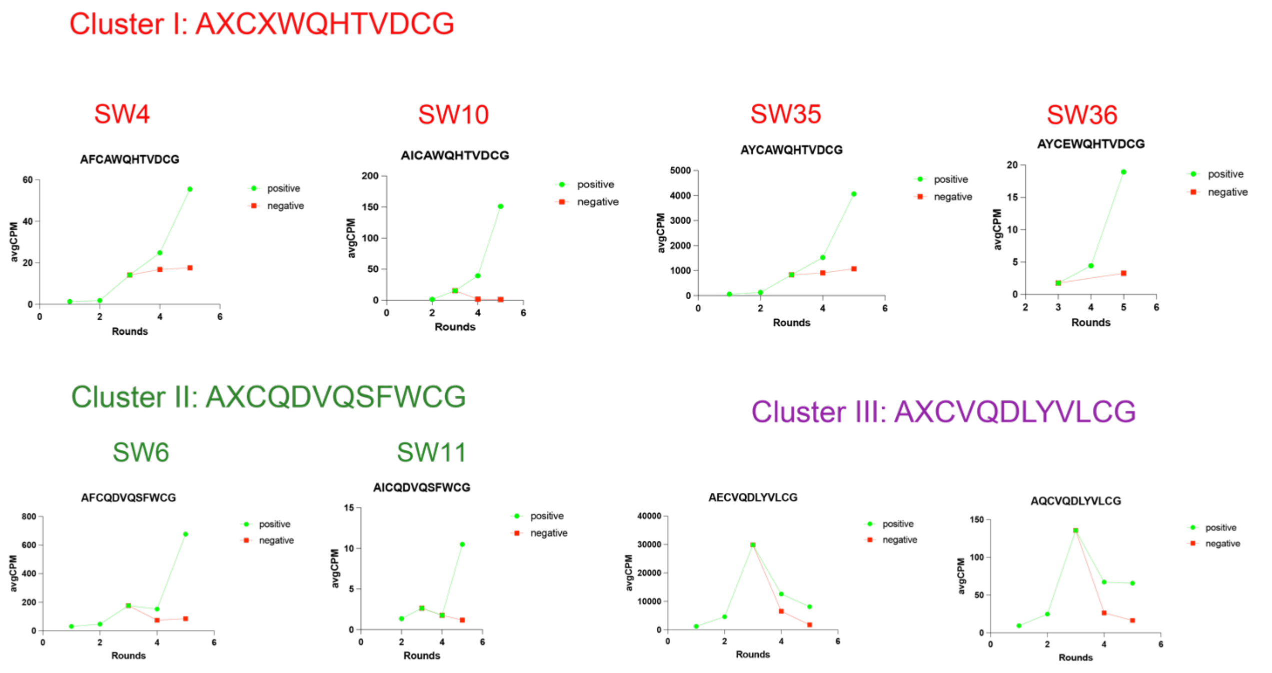


**Figure S10**: Enrichment profiles of hits within each cluster based on the DNA copy numbers (AvgCPM) for each round

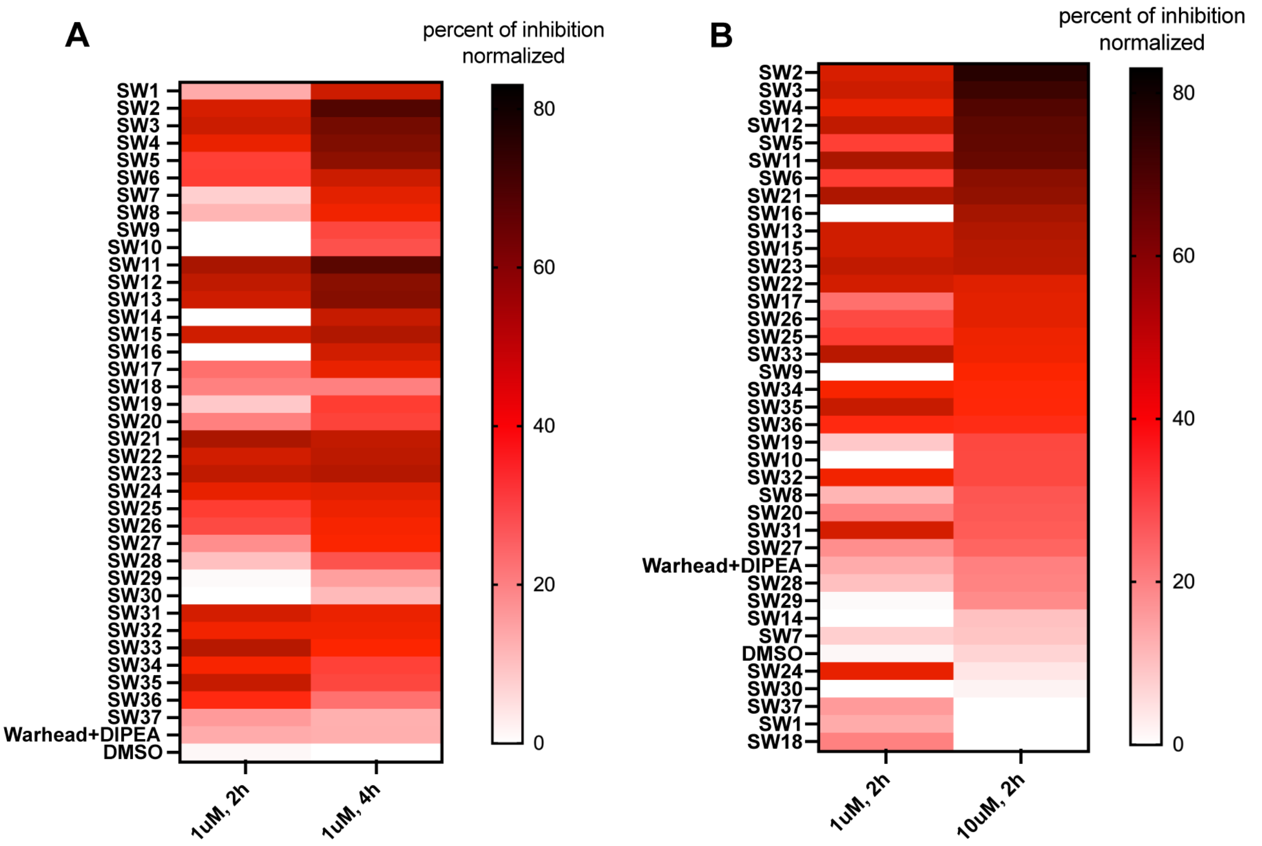


**Figure S11**: Time and dose-dependent activities of 37 compounds for Omicron BA2 spike using MSD plate. (A) 1 uM of stock compounds were pre-incubated with pre-coated 10 different SARS-CoV-2 full length spike variants with different incubation time length (t=2/4 hrs) on an MSD plate, before a competition binding against Sulfo-tagged ACE2. (B) 1 uM and 10uM of compounds were pre-incubated with pre-coated 10 different SARS-CoV-2 full length spike variants for 2 hrs on an MSD plate, before a competition binding against Sulfo-tagged ACE2. Warhead dibromomethyl aryl fluorosulfate warhead linker and DIPEA (used for cyclization) were doped in as negative controls, and DMSO was used as a solvent background control. Compound inhibition activity to Omicron BA2 spike was normalized using a calibration reagent provided with the MSD assay and plotted as a heatmap using Graphpad prism 10.

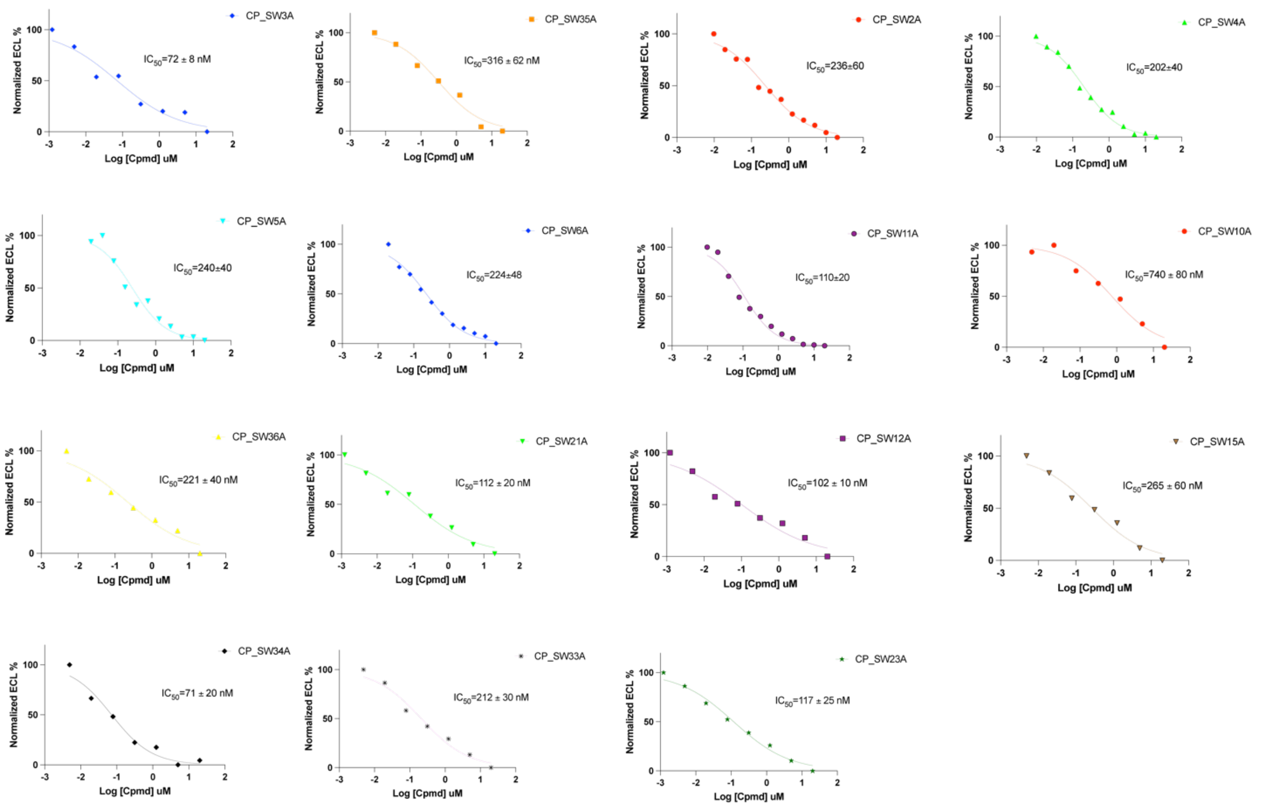


**Figure S12**: Full-dose inhibition activities of purified hits to Omicron BA2 spike using MSD assay. DMSO was used as a solvent background control. Compound inhibition activity to Omicron BA2 spike was normalized using a calibration reagent provided with the MSD assay and plotted as a heatmap using Graphpad prism 10.

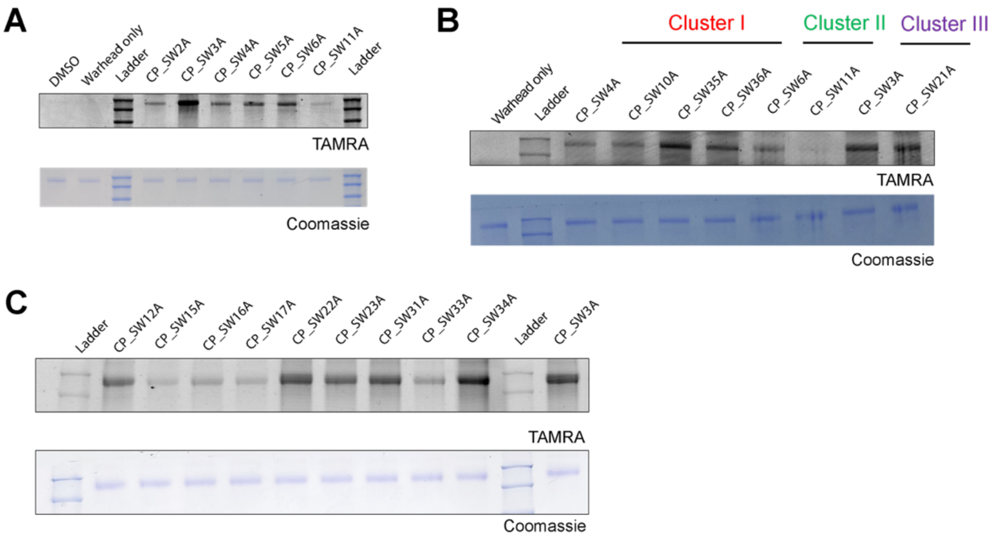


**Figure S13**: direct protein labeling of all 17 hits to spike. 10 uM compounds to full length spike Omicron BA2 variant were presented. In dose dependent labeling, 0.3 uM full length spike Omicron BA2 and 10 uM compounds were co-incubated in PBS for 2hr, in 37 °C. Labeled mixtures were clicked with an azide-TAMRA via CuAAC click reaction, followed by fluorescence scanning and Coomassie staining.

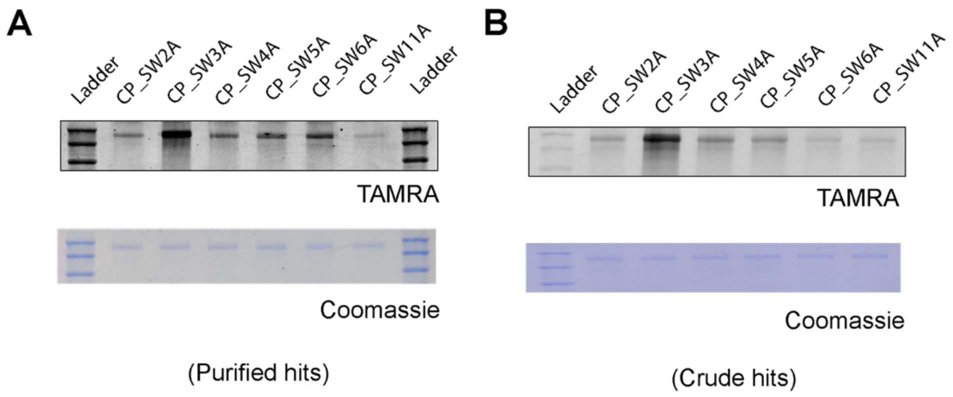


**Figure S14**: Comparing protein labeling using (A) purified hits or (B) crude hits. 0.3 uM full length spike Omicron BA2 and 10 uM compounds were co-incubated in PBS for 2hr, in 37 °C. Labeled mixtures were clicked with an azide-TAMRA via CuAAC click reaction, followed by fluorescence scanning and Coomassie staining.

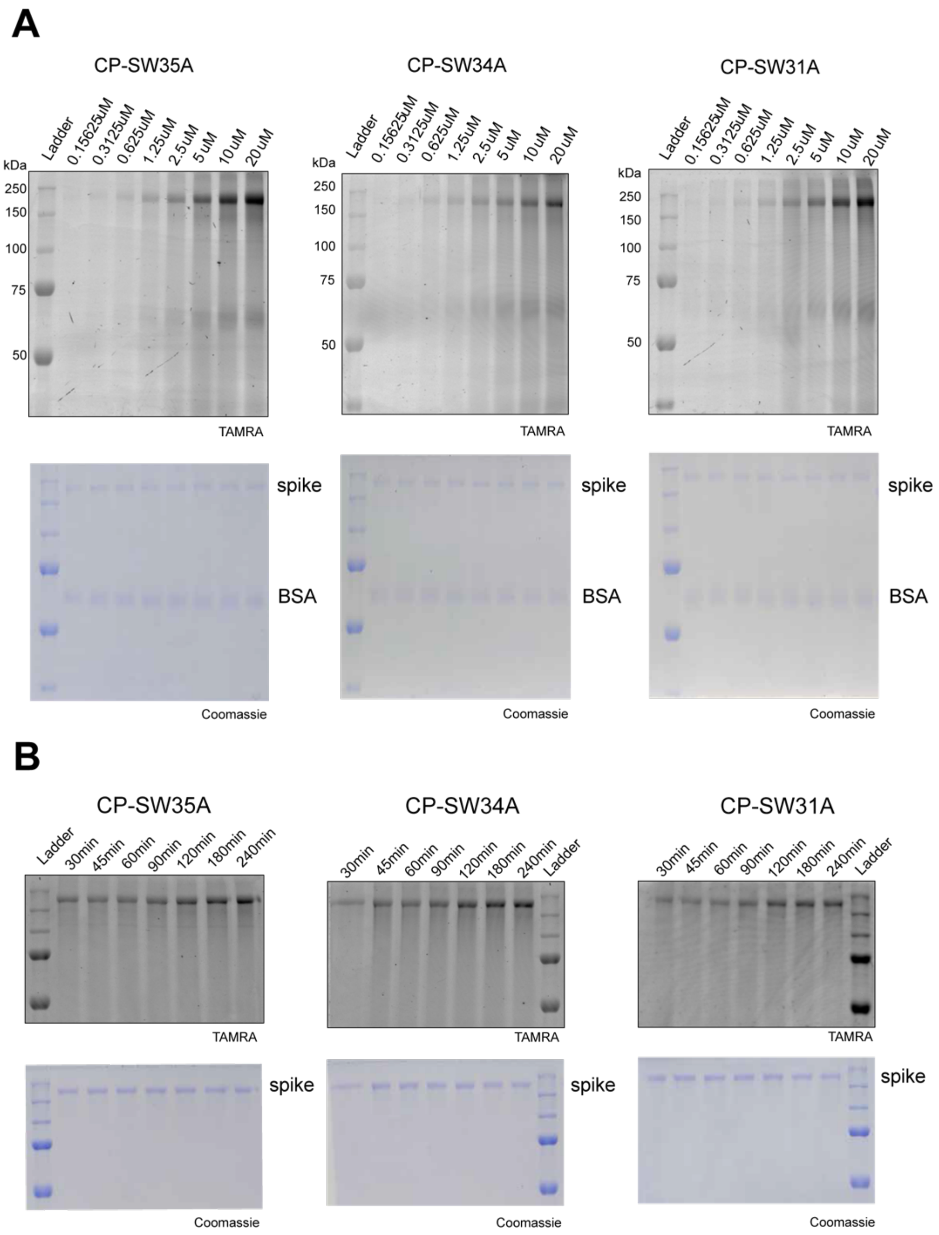


**Figure S15:** (A) Dose- and (B) time-dependent labeling of purified hits. In dose dependent labeling, 0.3 uM spike and 1.5 uM BSA were mixed and co-incubated with different annotated doses of CP-SW3A in PBS for 2hr, in 37C. Labeled mixtures were clicked with an azide-TAMRA via CuAAC click reaction, followed by fluorescence scanning and Coomassie staining. In time dependent labeling, 0.25 uM spike was incubated with 10 uM CP-SW3A in PBS, 37C, for different annotated time points. Labeled mixtures were clicked with a azide-TAMRA via CuAAC click reaction, followed by fluorescence scanning and Coomassie staining.

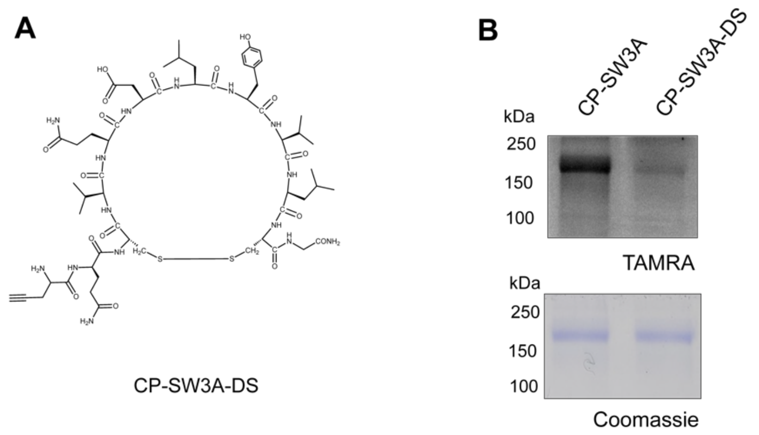


**Figure S16**: Spike labeling comparing CP-SW3A and negative control compound CP-SW3A-DS. (A) structure of compound CP-SW3A-DS (B) 0.3 uM spike is co-incubated with 10 uM CP-SW3A or 10 uM CP-SW3A-DS in PBS for 2hr, in 37 °C. Labeled mixtures were clicked with an azide-TAMRA via CuAAC click reaction, followed by fluorescence scanning and Coomassie staining.

**A**

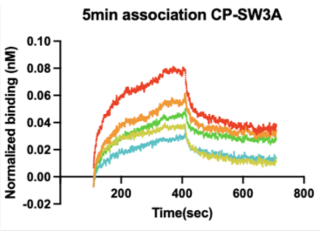


| KD (M) | KD Error | kon(1/Ms) | kon Error | kdis(1/s) | kdis Error |
|---|---|---|---|---|---|
| 6.78E-06 | 3.46E-07 | 4.82E+02 | 2.17E+01 | 3.27E-03 | 7.83E-05 |
| 3.68E-06 | 1.61E-07 | 5.70E+02 | 1.99E+01 | 2.10E-03 | 5.51E-05 |
| 3.93E-06 | 2.38E-07 | 1.55E+03 | 8.73E+01 | 6.09E-03 | 1.34E-04 |
| 1.00E-06 | 4.25E-08 | 1.89E+03 | 6.26E+01 | 1.89E-03 | 4.98E-05 |
| 1.07E-06 | 5.66E-08 | 3.04E+03 | 1.45E+02 | 3.25E-03 | 7.39E-05 |

Avg $k_{on}$ = 1.51 ± 0.067 E+03 (1/Ms) Avg $k_{off}$ = 3.32 ± 0.078 E-03 (1/s)

Avg KD = 2.2 ± 0.17 uM

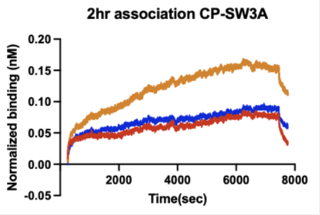


| KD (M) | KD Error | kon(1/Ms) | kon Error | kdis(1/s) | kdis Error | Full R^2 |
|---|---|---|---|---|---|---|
| 7.78E-07 | 3.35E-08 | 2.93E+03 | 1.15E+02 | 2.28E-03 | 3.99E-05 | 0.911 |
| 3.37E-07 | 1.16E-08 | 2.02E+03 | 4.67E+01 | 6.79E-04 | 1.72E-05 | 0.993 |
| 2.48E-07 | 9.92E-09 | 3.52E+03 | 9.31E+01 | 8.73E-04 | 2.62E-05 | 0.979 |

Avg $k_{on}$ = 2.82 ± 0.085 E+03 (1/Ms) Avg $k_{off}$ = 1.27 ± 0.028 E-03 (1/s)

Avg KD = 0.45 ± 0.018 uM

**B**

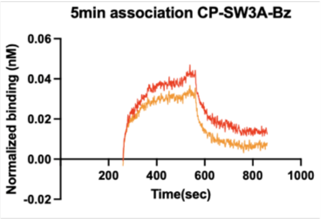


| KD (M) | KD Error | kon(1/Ms) | kon Error | kdis(1/s) | kdis Error | Full R^2 |
|---|---|---|---|---|---|---|
| 5.06E-06 | 2.61E-07 | 1.03E+03 | 4.88E+01 | 5.21E-03 | 1.04E-04 | 0.888 |
| 2.54E-06 | 3.43E-07 | 2.09E+03 | 1.43E+02 | 5.31E-03 | 1.79E-04 | 0.977 |

Avg $k_{on}$ = 1.56 ± 0.0959 E+03 (1/Ms) Avg $k_{off}$ = 5.26 ± 0.14 E-03 (1/s)

Avg KD = 3.89 ± 0.302 uM

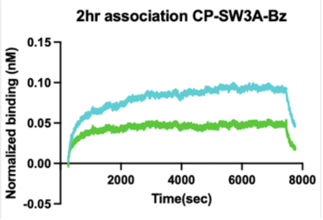


| KD (M) | KD Error | kon(1/Ms) | kon Error | kdis(1/s) | kdis Error | Full R^2 |
|---|---|---|---|---|---|---|
| 5.47E-06 | 1.53E-07 | 9.40E+02 | 1.32E+01 | 5.15E-03 | 2.27E-05 | 0.932 |
| 2.36E-06 | 9.85E-08 | 1.33E+03 | 5.05E+01 | 3.15E-03 | 5.52E-05 | 0.915 |

Avg $k_{on}$ = 1.14 ± 0.032 E+03 (1/Ms) Avg $k_{off}$ = 4.15 ± 0.039 E-03 (1/s)

Avg KD = 3.64 ± 0.12 uM

**Figure S17**: All reactions were run on an Octet RED96 at 30 °C, and samples were run in 1× PBS with 0.1% BSA and 0.05% Tween 20 (Octet buffer). IgGs were assessed for binding to biotinylated antigens using streptavidin biosensors (Sartorius/ForteBio). Biotinylated spike was loaded at a concentration of 200nM/100nM/50nM. Tips were then washed and baselined in wells containing only Octet buffer. A control well with loaded antigen but that was associated in a well containing only 200 µl of Octet buffer was used as a baseline subtraction for data analysis. Association and dissociation binding curves were fit in Octet System Data Analysis Software version 9.0.0.15 using a 1:2 bivalent model for IgGs to determine apparent Kd. Fold-change in apparent Kd were determined by computing the ratio of wildtype Kd to variant Kd. Averages of Kd fold-change values from at least two independent experiments are reported to two significant figures in tables. ± represents standard deviation errors.

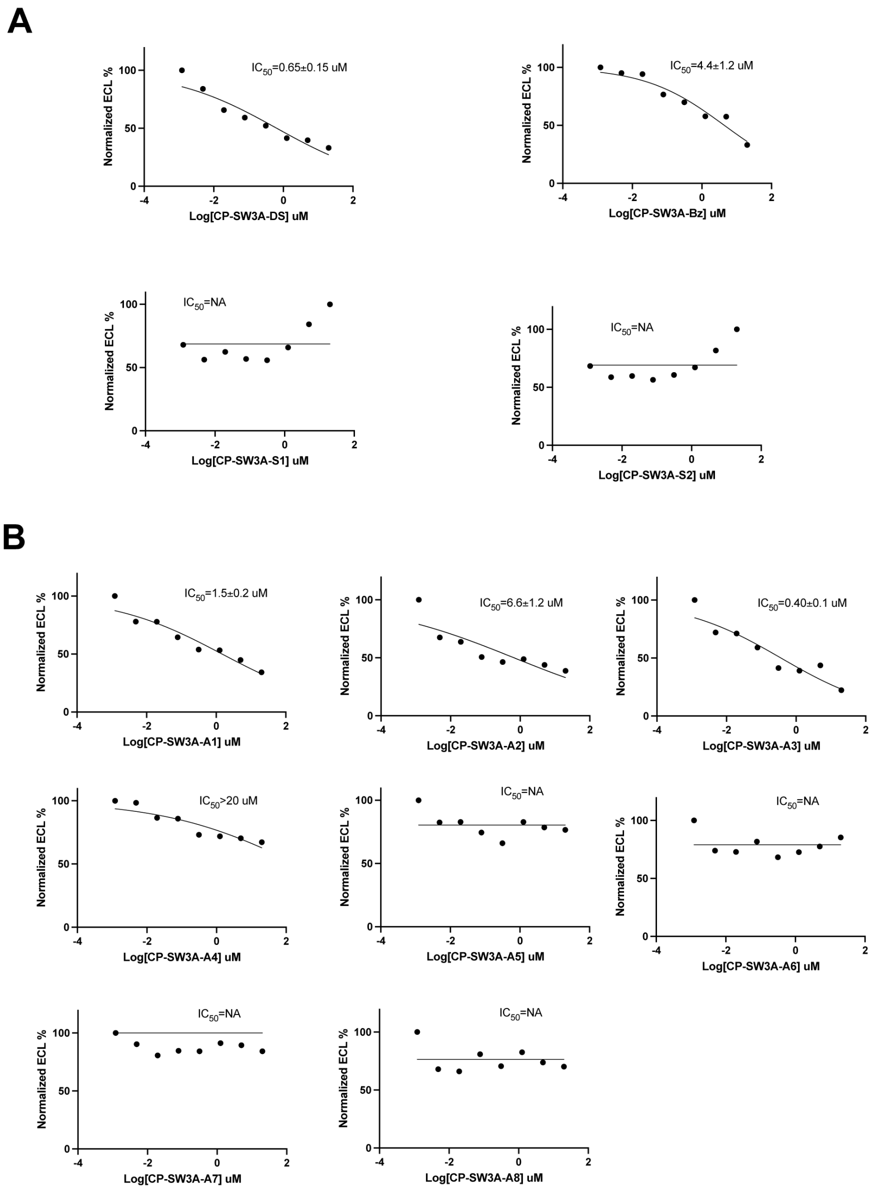


**Figure S18**: Inhibition activities of isomers, no warhead control (A) and alanine mutants (B) of CP-SW3A in MSD assay. DMSO was used as a solvent background control. Compound inhibition activity to Omicron BA2 spike was normalized using a calibration reagent provided with the MSD assay and plotted as a heatmap using Graphpad prism 10.

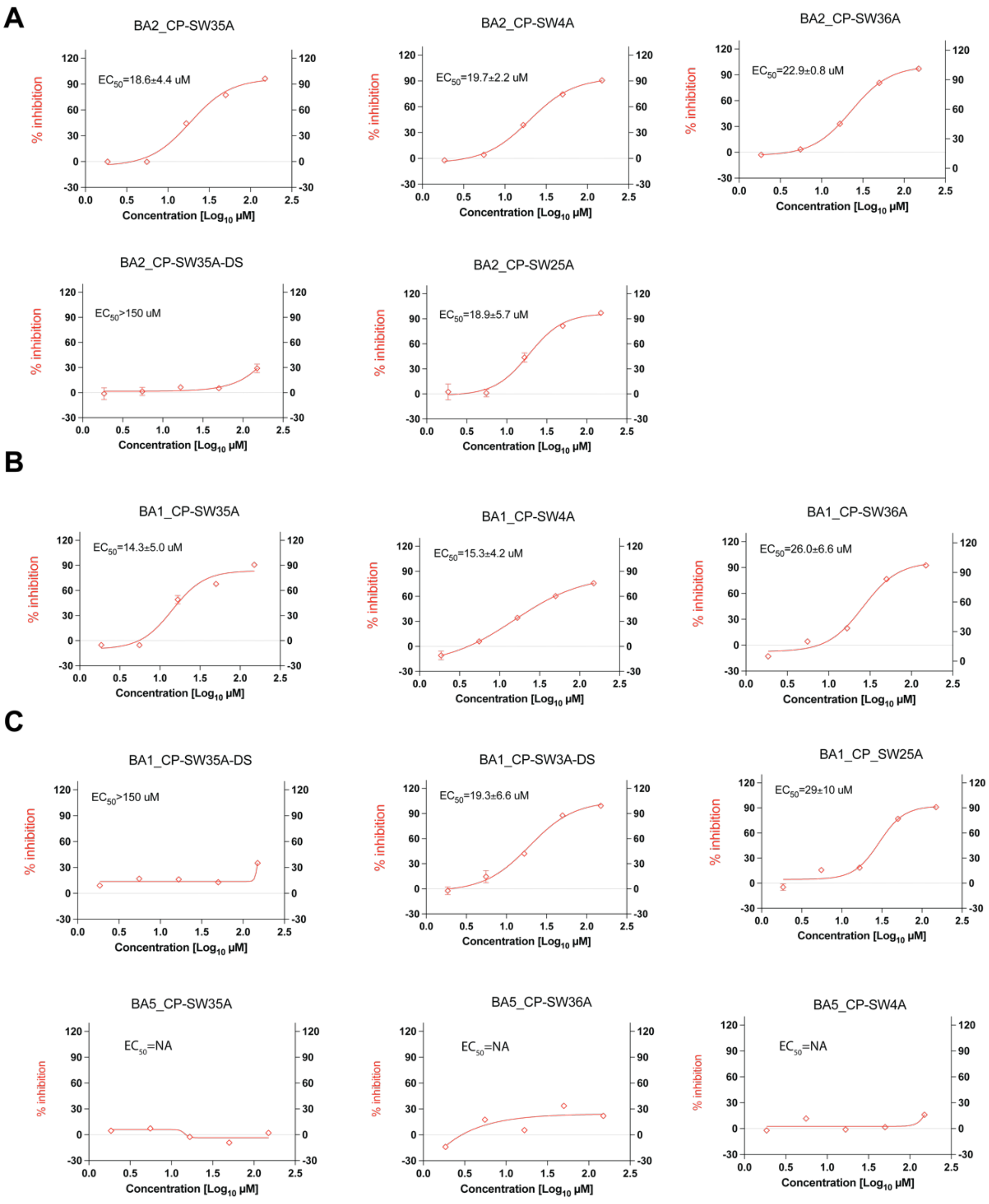


**Figure S19**: The neutralization capacity of compounds was assessed in titration experiments as previously described. SARS-CoV-2 viral particles, isolated from clinical samples, were incubated with compounds (or controls: sotrovimab as positive control, and DMSO as blank) in PBS for 1 hour in 37°C before infecting VeroE6 cells. Viral replication was determined at 24h post infection by immunostaining of intracellular viral nucleocapsid. Dose response neutralization activity of annotated compounds and positive control sotrovimab in antiviral assay using (A) Omicron BA5, (B) BA2 and (C) BA1 variants. Values were normalized to solvent control (1% DMSO) with infection (100% infection) and without infection (0% infection, assay background). EC50 values were calculated using non-linear regression dose response analysis by GraphPad Prism version 8.4.3 (GraphPad Software, San Diego CA, USA)

#### 2. Materials and Methods

##### General reagents for phage display:

*YT media with chloramphenicol (30ug/mL of chloramphenicol)*
Weight out 15.5 g of 2X YT broth (1.6% tryptone, 1.0% yeast extract, 0.5%sodium chloride) into 500mL of MilliQ water into a 2L flask. Add foil around it and autoclave it in the left autoclave on setting 1. Takes around 40 minutes. Remove the flask from the autoclave. Let it cool down to room temperature around 30 minutes. Make a final concentration of 30ug/mL of chloramphenicol into 2X YT broth. We have a 30mg/mL stock of chloramphenicol in ethanol. Add 500uL of this stock into 500mL of broth from autoclave.

*Agar plates with chloramphenicol (30ug/mL of chloramphenicol)*
Weight out 31 g of 2X YT broth (1.6% tryptone, 1.0% yeast extract, 0.5%sodium chloride) into a 2L flask. Weigh out 15 g of Agar into the vial. Then add 1000mL of MilliQ water. Add foil around it and autoclave it in the left autoclave on setting 1. Takes around 40 minutes. Remove the flask from the autoclave. Let it cool down and place int he water bath at 55C until ready. Make a final concentration of 30ug/mL of chloramphenicol into 2X YT broth. We have a 30mg/mL stock of chloramphenicol in ethanol. Add 1000uL of this stock into 500mL of broth from autoclave. Pour the plates by adding 25mL of media into a big circular plate. Let this cool overnight.

*20% PEG 8000 solution*
Weigh out 200 g of PEG 8000 and 145 g of NaCl. Place this into a large 4L glass container with lid. Autoclave on setting.

*Ammonium bicarbonate reaction buffer*

20mM Ammonium bicarbonate, 5mM EDTA in Milliq H2O at pH 8.75. For 50mL of reaction buffer 79 mg of AMBIC and 93 mg of EDTA(disodium, dihydrate ), pH to 8.75

*TCEP reducing buffer*

200mM of TCEP in 1mL Eppendorf. 57.3 mg of TCEP in 1mL of MilliQ water

*Washing buffers*

PBS (0.1% tween20), PBS (1% tween20), 5M guanidinium chloride

*Blocking buffer*

1mL of PBS (0.1% tween20) dissolved with 10mg of BSA

*TEV protease buffer*

TEV Protease Buffer is 50mM Tris Base, 0.5mM EDTA (Disodium) 1mM DTT(dithiothreitol). pH 8.0. Weigh out 302 mg of Tris Base, 73 mg EDTA, (Disodium), 7.7 mg of DTT. Dissolve in in 50mL of water and pH to 8.0.

*Biotinylated SARS-CoV-2 Spike Trimer Protein, His,Avitag™(BA.2/Omicron)*

Protein was purchased from ACROBiosystems (SPN-C82Er-200ug). This commercial protein is expressed from human 293 cells (HEK293) with T4 fibritin trimerization motif and a polyhistidine tag at the C-terminus. Mutations on this BA.2 spike protein: T19I, LPP24-26del, A27S, G142D, V213G, G339D, S371F, S373P, S375F, T376A, D405N, R408S, K417N, N440K, S477N, T478K, E484A, Q493R, Q498R, N501Y, Y505H, D614G, H655Y, N679K, P681H, N764K, D796Y, Q954H, N969K, R683A, R685A, F817P, A892P, A899P, A942P, K986P, V987P. Proline substitutions (F817P, A892P, A899P, A942P, K986P, V987P) and alanine substitutions (R683A and R685A) are introduced to stabilize the trimeric prefusion state of SARS-CoV-2 S protein.

*Human ACE2 / ACEH Protein, Fc Tag:*

ACE2 was purchased from ACROBiosystems (AC2-H5257-50ug). This commercial protein is glycosylated and expressed from human 293 cells.

*Nanobody Nb70 (56FFY):*

Nanobody is expressed and purified following the previous protocols[1], as a gift from Lei Wang lab in USCF.

#### Phage panning

Phage harboring the cysteine-rich peptide library ($CX_7C$) were used for selecting peptides against the Omicron BA2 Spike trimer. The randomized DNA sequences encoding the peptide library were inserted into the phage DNA between the pelB signal peptide and the disulfide-free pIII protein of the fdg3p0ss phage vector20. The phage peptide library bearing a TEV protease-cleavable site was used for selection of peptides. To overcome the low infectivity of the disulfide-free phage strain used in our screens, we used a large volume of 2YT rich medium (1 liter) for the production of phage. The phage peptide libraries in TG1 E. coli bacteria (Lucigen, 60502-1) were thawed from stock and used for inoculating 2YT medium containing 30 μg ml–1 chloramphenicol. Phage were generated by incubating at 30 °C with shaking at 250 r.p.m. for 16 h. On the second day, the medium rich with secreted phage was separated from the host bacteria by spinning at 8,500 r.p.m. for 30 min, and the supernatant was mixed with 250 ml of a solution containing 20% polyethylene glycol 8000 and 2.5 M NaCl. Following a 1-h incubation on ice, the precipitated phage were spun at 9,000 r.p.m. for 45 min. The resulting phage pellet was then dissolved in 1 ml buffer R (20 mM ammonium bicarbonate pH 8.0 and 5 mM EDTA) and reduced with 1 mM TCEP at R.T. for 30 h. After removing the excess TCEP by filtration using zeba column (7k, MWCO), 1.8 ml buffer R, 200ul acetonitrile, and 4ul of 20mM dibromo-OSO2F linker were added to modify the phage at 30 °C, 60rpm, for 1 h. Then, 0.5 ml of a solution with 20% polyethylene glycol 8000 and 2.5 M NaCl was added to precipitate the phage, and the recovered phage pellets were resuspended in selection buffer (50 mM Tris pH 8.0, 100 mM KCl, 1 mM EDTA and 1 mM DDT) for selections against TEV protease or FphF buffer (50 mM HEPES pH 7.5 and 100 mM NaCl) for selections against FphF hydrolase; both were supplemented with 1% BSA and 0.1% Triton X-100 for blocking at room temperature for 0.5 h. Biotin–TEV protease or FphF–biotin (10 μg) was incubated with 50 μl of hydrophilic streptavidin magnetic beads (New England Biolabs, S1421S) for 0.5 h, and beads were washed five times with TEV buffer supplemented with 1% BSA and 0.1%

Triton X-100. After being blocked in the same buffer for 0.5 h, the phage library was mixed with the magnetic beads and incubated at room temperature for different amounts of time depending on the round of screening being performed. After washing ten times with PBS, the magnetic beads were incubated with 100 µl guanidine chloride in PBS for 5 min, and then each was eluted with 10 µM TEV protease in 300 µl TEV buffer (50 mM Tris pH 8.0, 0.5 mM EDTA and 1 mM DTT) at 34 °C for 1 h to release the covalently bound phage from the solid support. The released phage were used to infect 45 ml of TG1 bacteria at an OD600 of 0.4 for 1.5 h in a 37 °C incubator without shaking. Afterwards, the TG1 cells were pelleted at 3,000 r.p.m. for 15 min and plated on two 15-cm 2YT agar plates with 30 µg ml–1 chloramphenicol antibiotic for each protease. The agar plate was then incubated at 37 °C overnight to allow the infected TG1 cells to expand. On the second day, TG1 cells were scraped off the agar plates with 5 ml 2YT medium, mixed with 5 ml 50% glycerol and stored at −80°C. This glycerol stock was then used for preparing plasmids for sequencing or for generating phage for the next round of phage panning.

**Phage infectivity measurement (phage tittering)**

Titering of phage was quantified by counting the number of infected TG1 colonies. Make proper serial dilutions of phage using PBS, and incubate 20ul of diluted phage with exponentially growing TG1 bacterial (OD600=0.4) for 90mins at 37C. After 90-min infection, add 10 ul of each tittering bacterial culture in triplicate on = 2YT agar plates containing 30 µg $ml^{-1}$ chloramphenicol overnight at 37C. Phage titers were calculated by colony counts and dilution factors.

**Post-translational chemical modification measurement**

From the TCEP treated phage (“CX7C TCEP”) and linker dibromo-OSO2F treated phage ((“CX7C+linker”) that were saved from the previous library preparations, take 25ul each into new tubes with 74uL ammonium bicarbonate buffer and 1uL of 100mM TCEP for 30 min reaction in R.T. Then, add 1uL of 100mM biotin polyethyleneoxide iodoacetamide (IADB) (Sigma-Aldrich, catalog# B2059) for 60 minutes reaction at R.T. Once the 60 min of reaction with IADB is complete. Label new tubes titled "TCEP BPD $10^2$" , "TCEP BPD $10^4$" , "TCEP BPD $10^5$ " and "Linker BPD $10^2$ " , "Linker BPD $10^4$ " , "Linker BPD $10^5$ ". (BPD: before pulldown) Add 990uL of PBS to 10uL from the the IADB treated “CX7C TCEP” and “CX7C linker” to make $10^2$ dilutions,  and once more to make $10^4$ dilutions. Then add 900uL of PBS to 100uL of $10^4$ dilution to make the $10^5$ dilution. Equilibrium and wash the streptavidin beads (use 10-20uL beads in excess of binding capacity) for pulldown in the next step. Take 950 uL of “TCEP BPD $10^5$” and “Linker BPD $10^5$” with the SA beads for 30min at R.T. After 30-min tumbling incubation, place the beads on magnetic holder and transfer the supernanent to new tubes labelled as “TCEP APD $10^5$” and “Linker APD $10^5$”. (APD:after pulldown). Generate an extra dilution of $10^6$ using the $10^5$ dilutions from samples before pulldown and after pulldown. Transfer 20uL of diluted phages to 180 exponentially growing TG1 bacterial cells for 90min infection at 37C. Add 10 ul of each tittering bacterial culture in triplicate ("TCEP BPD $10^5$”, "TCEP BPD $10^6$ ", "Linker BPD $10^5$ ", "Linker BPD $10^6$ ", "TCEP APD $10^5$ ", "TCEP APD $10^6$ ", "Linker APD $10^5$ ", "Linker APD $10^6$ ") on 2YT agar plates containing 30 µg $ml^{-1}$ chloramphenicol overnight at 37C. Phage titers were calculated by colony counts and dilution factors. Chemical modification efficiency can be calculated as yield%=(“Linker BPD”-“Linker APD”)/”Linker BPD”.

**Phage recovery/fold change for each panning**

For each library group, save 5-10uL of phage solutions for titer measurement before each panning and after panning. Phage recovery is calculated as the percentage of post-panning titer divided by pre-panning titer. For each panning cycle, no protein control panning (beads only) is used for calculating the fold change of phage recovery of sample (protein target) over control (beads only).

**Positive and counter selections**

For first three rounds of panning, positive selections indicate the panning of phage library to spike protein. For round 4 and 5, negative selections indicate the panning of phage library to the mixture of spike protein and ACE2 or nanobody Nb70 (56FFY). In negative selections, nanobody was incubated with spike for 4 hrs to form covalent interactions before panning against the phage library. Amount of ACE2 and Nb70 for each negative panning was determined using fraction bound equation to block >99% spike protein.

**Library DNA Preparation for NGS**

200 uL bacterial stocks containing amplified phage eluted from samples in all five rounds of panning were used for phage plasmid DNA extraction using DNA prep kit. 200 ng of phage plasmid for each sample from preparation was used for PCR amplification. Samples were evaluated by agarose gel electrophoresis to confirm the amplified products matching the calculated mobility shift. A second PCR amplification was performed to barcode samples directly from the first PCR procedure, using different combination pairs of forward and reverse primers. Amplified samples were checked using agarose gel electrophoresis to confirm the mobility shift of new products matching with the calculated molecular weight. Gel bands with expected molecular weight after amplification were separated and cut by clean razorblades for DNA extraction and purification using the commercial gel DNA recovery kit (Zymo Research, #D4008). Final qualified DNA samples after purifications were submitted for deep sequencing from NGS service provided by MedGenome, Inc. (Foster City, CA). Samples were pooled and read on the NovaSeq, with 2 million paired reads (paired end 150 reads) for each sample.

**Analysis of Phage Display sequencing data**

Paired reads were merged using the fastq_mergepairs command of the vsearch package. Next, adapters were clipped from the 5’ and 3’ end of merged reads using cutadapt. DNA sequences were translated into protein sequences using the translate function of Biostrings.
Using RStudio, data was filtered to exclude peptides with mutated constant amino acid positions or with cysteines in variable amino acid positions. We employed the DESeq2 package (Love et al, 2014, doi:10.1186/s13059-014-0550-8) for differential enrichment analysis, following standard procedures(https://bioconductor.org/packages/release/bioc/vignettes/DESeq2/inst/doc/DESeq2.html). This method employs the prior distribution of log fold changes (LFC) to calculate LFC and statistical significance for all peptides, even when they are not detected in all samples.

**Bioinformatic Pipeline for Counter Selection Data Analysis**

```
---
title: "Exploratory analysis of Phage Display Data"
```

````
date: '`r strftime(Sys.time(), format = "%B %d, %Y")`'
output:
  html_document:
    keep_md: yes
    code_folding: hide
    highlight: tango
    theme: united
    toc: yes
    toc_depth: 2
    toc_float: yes
  pdf_document:
    toc: yes
    toc_depth: '2'
  word_document:
    toc: yes
    toc_depth: '2'
editor_options:
  chunk_output_type: inline
---


## General information:

This pipeline performs exploratory analysis of Phage Display Data ...
It requires ....

# Set up

```{r setup, cache=FALSE, message=FALSE, warning=FALSE, comment = FALSE}

knitr::opts_chunk$set(echo = TRUE)
````
````

````
library(ggplot2)
library(readr)
# library(GGally)
library(stringr)
library(purrr)
library(RColorBrewer)
library(pheatmap)
library(tidyverse)
library(dplyr)
library(ggpubr)
library(tidyr)
library(DESeq2)
library(ggrepel)
library(EnhancedVolcano)
library(ggforce)
library(ggseqlogo)
library(ggVennDiagram)
library(GGally)


```

### Data import

```{r import_data, message=FALSE, warning = FALSE, echo=FALSE}

metadata <- read_csv("metadata.csv", show_col_types = FALSE)

df <- readRDS("merged_df_complete.rds") %>%
  dplyr::rename(sample_id_incorrect = sample) %>%
  full_join(metadata, by = "sample_id_incorrect") %>%
````

````
  select(-sample_id_incorrect, "Details for samples")

df %>% dplyr::rename(peptide = x) -> df

df$peptide <- str_replace_all(df$peptide, '\\*', 'Q')

head(df)

```

### QC of peptide lenghts and trimming to 12 aa

```{r peptide_lentgh, message=FALSE, warning = FALSE, echo=FALSE}

df %>%
  mutate(pep_length = str_length(peptide)) -> df

df %>%
  ggplot(aes(x=pep_length)) +
  geom_histogram(alpha=0.5, position = 'identity') +
  # scale_x_continuous(trans = "log10") +
  # facet_zoom(ylim = c(0, upper_ylim * 1.1), zoom.size = 1) +
  ggtitle("Peptide lengths pre-filtering") +
  theme_bw() +
  facet_wrap(~sample)

df %>%
  mutate(pep=substr(peptide, 1, 12)) %>%
  group_by(pep) %>%
  summarise(n=sum(n)) %>%
  ungroup() -> pep
````

```
ggseqlogo(unique(pep$pep)) +
  theme(legend.position = 'none') +
  ggtitle("Peptide composition after truncation to 12 aa")

# truncate all peptides to 12 aa length in all dataframes

df %>%
  mutate(pep=substr(peptide, 1, 12)) %>%
  group_by(sample_id, pep) %>%
  summarize(count=sum(n)) %>%
  full_join(metadata, by="sample_id") %>%
  mutate(CPM = ((count+1) / sum(count) * 1e6)) -> CPM_df

CPM_df$sample_id <- factor(CPM_df$sample_id, levels = metadata$sample_id)

#remove peptides that have mutated constant aa or cysteines in variable aa
test_mut <- function(x){
 const_aa <- paste(unlist(strsplit(x,""))[c(1, 3, 11, 12)], collapse = "")
 variable_aa <- paste(unlist(strsplit(x,""))[c(2, 4, 5, 6, 7, 8, 9, 10)], collapse = "")
 return(c(const_aa, variable_aa))
  }

CPM_df$test_mut <- lapply(CPM_df$pep, test_mut)

CPM_df %>%
  mutate(test_mut = purrr::map(test_mut, setNames, c("const_aa","variable_aa"))) %>%
  unnest_wider(test_mut) -> CPM_df

CPM_df %>%
  group_by(const_aa) %>%
```

```
  tally() %>%
  filter(n>100) %>%
  arrange(desc(n)) %>%
  mutate(group = ifelse(const_aa == "ACCG", "ACCG", "mutant"))%>%
  ggplot(aes(x=const_aa, y = n, fill=group)) +
  geom_bar(stat="identity") +
  theme_bw() +
  theme(axis.text.x = element_text(angle = 90), text = element_text(size = 10)) +
  scale_y_continuous(trans="log2") +
  ggtitle("Frequency of constant position combinations with n>100")


CPM_df %>%
  filter(const_aa == "ACCG") %>%
  filter(!grepl("C",variable_aa)) %>%
  select(-const_aa, -variable_aa)-> CPM_df

CPM_df %>%
  select(pep, sample_id, count) %>%
  spread(key = sample_id, value = count) %>%
  mutate_all(~replace(., is.na(.), 0)) %>%
  column_to_rownames('pep') -> large_df
```

- I truncated all peptides to 12 aa. Sequence logo shows the expected aa composition after truncation.
- The vast majority of peptides has the right amino acids at constant positions, peptides with mutated constant positions or with C at variable positions are discarded.
- There are ~542000 peptide species detected in total across all samples.

### Quality control of count and abundance metrics

```{r count_QC, message=FALSE, warning = FALSE, echo=FALSE}

CPM_df %>%
  ggplot(aes(x=count, color=sample_id)) +
  geom_histogram(fill="white", position="identity") +
  scale_x_continuous(trans = "log10") +
  scale_y_continuous(trans = "log10") +
  theme_minimal() +
  theme(text = element_text(size = 12), legend.position = "none")+
  ggtitle("Composition of all libraries") +
  facet_wrap(factor(CPM_df$sample_id, levels = unique(metadata$sample_id)))

CPM_df %>%
  group_by(sample_id) %>%
  tally(count) %>%
  dplyr::rename(total_count = n) %>%
  full_join(metadata) -> count_df

count_df %>%
  ggplot(aes(x=round, y=total_count, fill= as.character(rep))) +
  geom_bar(stat="identity", position = position_dodge()) +
  theme_minimal() +
  theme(axis.text.x = element_text(angle = 90), text = element_text(size = 12))+
  scale_fill_brewer(palette="Blues", direction = -1) +
  ggtitle("Total counts per sample") +
  facet_wrap(factor(count_df$sample, levels = unique(metadata$sample)))

# CPM_df %>%
#   ggplot(aes(x=as.character(round), y=CPM, fill= as.character(rep))) +
```

```
#   geom_boxplot()+
#   theme_minimal() +
#   theme(axis.text.x = element_text(angle = 90), text = element_text(size = 12)) +
#   scale_y_continuous(trans = "log10") +
#   xlab("Panning round") +
#   ggtitle("CPM distribution per sample") +
#   scale_fill_brewer(palette="Blues", direction = -1) +
#   facet_wrap(factor(CPM_df$sample, levels = unique(metadata$sample)))

# Total detected peptides per sample (with or without threshold)
CPM_df %>%
  group_by(sample_id) %>%
  tally() %>%
  dplyr::rename(total_peptides = n) %>%
  full_join(metadata) -> detected_peptides_df

detected_peptides_df %>%
  ggplot(aes(x=as.character(round), y=total_peptides, fill = as.character(rep))) +
  geom_bar(stat="identity", position = position_dodge()) +
  theme_minimal() +
  theme(axis.text.x = element_text(angle = 90), text = element_text(size = 12)) +
  ggtitle("Total detected peptides") +
  scale_fill_brewer(palette="Blues", direction = -1) +
  facet_wrap(factor(detected_peptides_df$sample, levels = unique(metadata$sample))) +
  geom_text(aes(label=total_peptides, angle = 90), vjust=0, hjust=-0.5, check_overlap = TRUE,
size=3)
  # scale_y_continuous(limits = c(NA, 100000))

CPM_df %>%
  filter(count > 10) %>%
  group_by(sample_id) %>%
```

````
  tally() %>%
  dplyr::rename(total_peptides = n) %>%
  full_join(metadata) -> detected_peptides_df10
detected_peptides_df10 %>%
  ggplot(aes(x=as.character(round), y=total_peptides, fill = as.character(rep))) +
  geom_bar(stat="identity", position = position_dodge()) +
  theme(text = element_text(size = 12)) +
  theme_minimal() +
  ggtitle("Total detected peptides (n > 10)") +
  scale_fill_brewer(palette="Blues", direction = -1) +
  facet_wrap(factor(detected_peptides_df10$sample, levels = unique(metadata$sample))) +
  geom_text(aes(label=total_peptides, angle = 90), vjust=0, hjust=-0.5, check_overlap = TRUE, size=3) +
  scale_y_continuous(limits = c(NA, 100000))

# ggsave("detected_peptides_10.png")

```
````

# How are detected peptides shared across samples?

I asked how many of the peptides that are detected in round5 samples are detected exclusively by one or multiple samples. This is important to understand if the peptides enriched in each sample make sense and to decide how to make downstream enrichment analysis with Deseq2.

````
```{r peptides_intersection, message=FALSE, warning = FALSE, echo=FALSE}

filter(metadata, `Sample ID` %in% c("SW_11", "SW_12", "SW_13", "SW_17", "SW_18", "SW_19", "SW_23", "SW_24", "SW_25", "SW_29", "SW_30", "SW_31")) %>%
  select(sample_id, sample, round, rep)-> coldata

row.names(coldata) <- coldata$sample_id
````

```
large_df %>%
  select(rownames(coldata)) -> large_df_filtered

# dataframe of peptides with at least one count by sample (merging peptides detected in at least one rep of each sample)
large_df_filtered %>%
  mutate(peptide=row.names(.)) %>%
  gather(key="sample_id", value="count", -peptide) %>%
  inner_join(metadata, by="sample_id") %>%
  select(sample, peptide, count) %>%
  group_by(sample, peptide) %>%
  summarise(count=sum(count)) %>%
  filter(count > 0) %>%
  select(sample, peptide) %>%
  group_by(sample) -> grouped_df

with(grouped_df, split(peptide,
     factor(sample, levels = unique(sample)))) -> pep_list

# dataframe of peptides with peptides detected at least once once in all reps of the same sample
large_df_filtered %>%
  mutate(peptide=row.names(.)) %>%
  gather(key="sample_id", value="count", -peptide) %>%
  full_join(metadata, by="sample_id") %>%
  group_by(peptide, sample) %>%
  filter(all(count>0)) %>%
  select(sample, peptide) %>%
  group_by(sample) -> grouped_df_nozero

with(grouped_df_nozero, split(peptide,
```

```
    factor(sample, levels = unique(sample)))) -> pep_list_mean
```

```{r Venn, message=FALSE, warning = FALSE, echo=FALSE}

ggVennDiagram(pep_list) +
  scale_fill_gradient(low="grey", high = "firebrick") +
  scale_color_manual(values=c(rep("black", 4))) +
  ggtitle("Detected peptides in round 5 samples", subtitle = "(included are peptides detected in at least one replicate of each sample)")

ggVennDiagram(pep_list_mean) +
  scale_fill_gradient(low="grey", high = "firebrick") +
  scale_color_manual(values=c(rep("black", 4))) +
  ggtitle("Detected peptides in round 5 samples", subtitle = "(included are peptides detected at least once in all replicates of each sample)")

ggVennDiagram(pep_list_mean[1:3]) +
  scale_fill_gradient(low="grey", high = "firebrick") +
  scale_color_manual(values=c(rep("black", 4))) +
  ggtitle("Detected peptides in round 5 samples", subtitle = "(included are peptides detected at least once in all replicates of each sample)")

ggVennDiagram(pep_list_mean[c(1, 4)]) +
  scale_fill_gradient(low="grey", high = "firebrick") +
  scale_color_manual(values=c(rep("black", 4))) +
  ggtitle("Detected peptides in round 5 samples", subtitle = "(included are peptides detected at least once in all replicates of each sample)")
names(pep_list_mean)
```

````
```
````

- When looking at all detected peptides, most are detected in only one sample and not in any other one (darker red areas are the external ones)
- When restricting analysis to only peptides detected repliably across replicates (>1 count in all reps of each sample), 55% are detected in beads only. This is the most minimal threshould one can apply.

Isn't it strange that the vast majority of peptides detected in the beads sample is not detected in any other sample? I would expect beads to enrich a general background which should be also enriched by other samples at least in part.

- Only 386 peptides are detected repliably in all 4 samples. Therefore DEseq analysis including all samples in the same design should be excluded, as too little peptides are shared across samples.

# Adding NL (from Franco's data) to the picture

````
```{r peptides_intersection_NL, message=FALSE, warning = FALSE, echo=FALSE}

readRDS("large_df_NL.rds") %>%
  rownames_to_column("peptide") %>%
  full_join(rownames_to_column(large_df_filtered, "peptide"), by='peptide') %>%
  mutate_all(~replace(., is.na(.), 0)) %>%
  column_to_rownames('peptide') %>%
  filter(rowSums(.)>0) -> large_df_filtered_NL

NL_rows <- data.frame(sample_id = c("NL_rep1", "NL_rep2", "NL_rep3"),
                      sample = c("NL", "NL", "NL"),
                      round = c(0, 0, 0),
                      rep = c(1, 2, 3))

coldata %>%
````

```
  bind_rows(NL_rows) -> coldata

# dataframe of peptides with peptides detected at least once once in all reps of the same sample
large_df_filtered_NL %>%
  rownames_to_column("peptide") %>%
  gather(key="sample_id", value="count", -peptide) %>%
  full_join(coldata, by="sample_id") %>%
  group_by(peptide, sample) %>%
  filter(all(count>0)) %>%
  select(sample, peptide) %>%
  group_by(sample) -> grouped_df_nozero_NL

with(grouped_df_nozero_NL, split(peptide,
     factor(sample, levels = unique(sample)))) -> pep_list_mean_NL

```

```
```{r Venn_NL, message=FALSE, warning = FALSE, echo=FALSE}

ggVennDiagram(pep_list_mean_NL[c(1, 2)]) +
  scale_fill_gradient(low="grey", high = "firebrick") +
  scale_color_manual(values=c(rep("black", 5))) +
  ggtitle("Detected peptides", subtitle = "(included are peptides detected at least once in all replicates of each sample)")

ggVennDiagram(pep_list_mean_NL[c(1, 5)]) +
  scale_fill_gradient(low="grey", high = "firebrick") +
  scale_color_manual(values=c(rep("black", 5))) +
  ggtitle("Detected peptides", subtitle = "(included are peptides detected at least once in all replicates of each sample)")
```
```

````
ggVennDiagram(pep_list_mean_NL[c(1, 3)]) +
  scale_fill_gradient(low="grey", high = "firebrick") +
  scale_color_manual(values=c(rep("black", 5))) +
  ggtitle("Detected peptides", subtitle = "(included are peptides detected at least once in all replicates of each sample)")

ggVennDiagram(pep_list_mean_NL[c(1, 4)]) +
  scale_fill_gradient(low="grey", high = "firebrick") +
  scale_color_manual(values=c(rep("black", 5))) +
  ggtitle("Detected peptides", subtitle = "(included are peptides detected at least once in all replicates of each sample)")

# names(pep_list_mean_NL)

```
- About 50% of peptides detected in any of the samples with protein on beads (spike, ACE2 and nano) are not detected in the native library
- Given that the overlap is not so much better with the NL library, it probably doesn't help to use this for differential enrichment analysis.

### Principal component analysis

```{r PCA, message=FALSE, warning = FALSE, echo=FALSE}

large_df %>%
  filter (rowSums(.) >= 5) -> large_df_5



#  check if there is any 0 in this large dataset
large_df_5[apply(large_df_5[,], 1, min) != 0, ] -> nonzero_large
#  only 90 peptides in the filtered dataset have no 0 in any sample
````

~~~
### Create a `DESeqDataSet` object to normalize and transform data
dds <- DESeqDataSetFromMatrix(
  countData = large_df_5, # the counts values for all samples in our dataset
  colData = metadata, # annotation data for the samples in the counts data frame (rows should
correspond to columns in large_df)
  design = ~1 # Here we are not specifying a model because we are not performing DE analysis
  # Replace with an appropriate design variable for your analysis
)

### Normalize and transform the data in the `DESeqDataSet` object
### using the `vst()` function from the `DESeq2` R package
dds_norm <- vst(dds)

### plotPCA(dds_norm, intgroup = c("sample")) +
  # geom_text_repel(aes(label = dds_norm$round), size = 3.5)

row.names(metadata) <- metadata$sample_id
plot_data <- plotPCA(dds_norm,intgroup="sample",returnData=TRUE)
plot_data <- bind_cols(plot_data, metadata)  %>%
  dplyr::rename(sample = "sample...4")

ggplot(plot_data, aes(x = PC1,y=PC2, col=sample)) +
  geom_point(size=3, alpha=0.5) +
  geom_text_repel(aes(label = plot_data$round), size = 3.5) +
  xlab("PC1 (93%)") + ylab("PC2 (3%)")


```
~~~

Most of the variance is explained by PC1. This suggests that panning against beads did not change substantially the population of peptides, while panning against the spike protein did have an impact mostly in the first 3 rounds.

# DEseq2 analysis

Even though there is very low overlap of detected peptides among samples, we can use DEseq to include all peptides, by removing any filtering for outliers.
Making pairwise comparisons, this means that enrichment for a peptide will be extrapolated based on the prior distribution of LFC, even if it is only detected in one of the two samples under comparison.

Explanation by the Deseq author of this extrapolation:
[regarding calculation of log fold changes when an element is not detected in one of the samples] While the maximum likelihood estimate (MLE) of DESeq goes to Inf, the use of a prior distribution on LFCs (log fold changes) in DESeq2 gives us a finite estimate. The way to interpret this is that: zeros might indicate absolute no fragments in samples of A, or more likely that the expected counts of fragments is some positive value below 1. If we were to increase the sequencing depth by 10 or 100, etc., we might start to observe some fragments in A. The prior distribution for LFCs is estimated by looking at the distribution of MLE fold changes observed, including other genes where the sequencing depth is higher, and using this range to give a finite estimate here. (See our paper for full details http://genomebiology.com/2014/15/12/550/abstract.) So the estimate here depends on: the dispersion for this gene, how large the counts are for B, and the distribution of log fold changes for other genes which had finite MLE LFCs.

````
```{r spike_vs_beads, message=FALSE, warning = FALSE, echo=FALSE, fig.height=7}

metadata %>%
  filter(sample %in% c("spike", "beads"), round == 5) %>%
  select(sample_id, sample, round, rep)-> coldata
````
````

```
row.names(coldata) <- coldata$sample_id

select(large_df_5, row.names(coldata)) %>%
  filter(rowSums(.)>0) -> large_df_spikebeads

dds_raw <- DESeqDataSetFromMatrix(large_df_spikebeads,
                                  colData = coldata,
                                  design = ~ sample)

### # pre-filtering: ensure that at least X samples with a count of 5 or more, where X can be chosen as the sample size of the smallest group of samples (in this case 6 samples are the minimum number of samples in one comparison)
keep <- rowSums(counts(dds_raw) >= 1) >= 3
dds <- dds_raw[keep,]
dds$sample<- relevel(dds$sample, ref = "beads")
### run Deseq
dds <- DESeq(dds)

res_splike_vs_beads <- results(dds, alpha=0.05, independentFiltering=FALSE)

resOrdered_splike_vs_beads <- res_splike_vs_beads[order(res_splike_vs_beads$padj),]
### resOrdered_splike_vs_beads
summary(res_splike_vs_beads)

### resultsNames(dds)
resLFC_spike <- lfcShrink(dds, res= res_splike_vs_beads, coef = "sample_spike_vs_beads", type="apeglm")

EnhancedVolcano(resLFC_spike,
    lab = rownames(resLFC_spike), x = 'log2FoldChange', pCutoff = 0.05, FCcutoff = 1,
```

```
    y = 'padj', title = "spike vs beads", subtitle = "(all peptides detected at least once in at least one of the replicates included)")

```
```

The "wing" shape of the vulcano plot reflects the low counts in at least one of the two samples under comparison, which forces the program to estimate fold changes and significance.

### Differential enrichment of ACE2 over beads

```{r ace2_vs_beads, message=FALSE, warning = FALSE, echo=FALSE, fig.height=7}

metadata %>%
  filter(sample %in% c("ACE2", "beads"), round == 5) %>%
  select(sample_id, sample, round, rep)-> coldata

row.names(coldata) <- coldata$sample_id

select(large_df_5, row.names(coldata)) %>%
  filter(rowSums(.)>0) -> large_df_volcano

dds_raw <- DESeqDataSetFromMatrix(large_df_volcano,
                                  colData = coldata,
                                  design = ~ sample)

# # pre-filtering: ensure that at least X samples with a count of 5 or more, where X can be chosen as the sample size of the smallest group of samples (in this case 6 samples are the minimum number of samples in one comparison)
keep <- rowSums(counts(dds_raw) >= 1) >= 3
dds <- dds_raw[keep,]
dds$sample<- relevel(dds$sample, ref = "beads")
```
```

~~~
# run Deseq
dds <- DESeq(dds)

res_ACE2_vs_beads <- results(dds, alpha=0.05, independentFiltering=FALSE)

# resOrdered_ACE2_vs_beads <- res_ACE2_vs_beads[order(res_ACE2_vs_beads$padj),]
# resOrdered_splike_vs_beads
summary(res_ACE2_vs_beads)

# resultsNames(dds)
resLFC <- lfcShrink(dds, res = res_ACE2_vs_beads, coef = "sample_ACE2_vs_beads",
type="apeglm")

EnhancedVolcano(resLFC,
   lab = rownames(resLFC), x = 'log2FoldChange', pCutoff = 0.05, FCcutoff = 1,
   y = 'padj', title = "ACE2_vs_beads")

```
~~~

### Differential enrichment of nano over beads

~~~
```{r nano_vs_beads, message=FALSE, warning = FALSE, echo=FALSE, fig.height=7}

metadata %>%
  filter(sample %in% c("nano", "beads"), round == 5) %>%
  select(sample_id, sample, round, rep)-> coldata

row.names(coldata) <- coldata$sample_id

select(large_df_5, row.names(coldata)) %>%
  filter(rowSums(.)>0) -> large_df_volcano
~~~

```
dds_raw <- DESeqDataSetFromMatrix(large_df_volcano,
                          colData = coldata,
                          design = ~ sample)

### # pre-filtering: ensure that at least X samples with a count of 5 or more, where X can be chosen as the sample size of the smallest group of samples (in this case 6 samples are the minimum number of samples in one comparison)
keep <- rowSums(counts(dds_raw) >= 1) >= 3
dds <- dds_raw[keep,]
dds$sample<- relevel(dds$sample, ref = "beads")
### run Deseq
dds <- DESeq(dds)

res_nano_vs_beads <- results(dds, alpha=0.05, independentFiltering=FALSE)

resOrdered_nano_vs_beads <- res_nano_vs_beads[order(res_nano_vs_beads$padj),]
### resOrdered_splike_vs_beads
summary(res_nano_vs_beads)

### resultsNames(dds)
resLFC <- lfcShrink(dds, res = res_nano_vs_beads, coef = "sample_nano_vs_beads", type="apeglm")

EnhancedVolcano(resLFC,
   lab = rownames(resLFC), x = 'log2FoldChange', pCutoff = 0.05, FCcutoff = 1,
   y = 'padj', title = "nano_vs_beads")
```

# Differential enrichment of spike over ACE2

```
```{r spike_vs_ace2, message=FALSE, warning = FALSE, echo=FALSE, fig.height=7}

metadata %>%
  filter(sample %in% c("spike", "ACE2"), round == 5) %>%
  select(sample_id, sample, round, rep)-> coldata

row.names(coldata) <- coldata$sample_id

select(large_df_5, row.names(coldata)) %>%
  filter(rowSums(.)>0) -> large_df_volcano

dds_raw <- DESeqDataSetFromMatrix(large_df_volcano,
                                  colData = coldata,
                                  design = ~ sample)

# # pre-filtering: ensure that at least X samples with a count of 5 or more, where X can be chosen as the sample size of the smallest group of samples (in this case 6 samples are the minimum number of samples in one comparison)
keep <- rowSums(counts(dds_raw) >= 1) >= 3
dds <- dds_raw[keep,]
dds$sample<- relevel(dds$sample, ref = "ACE2")
# run Deseq
dds <- DESeq(dds)

res_splike_vs_ACE2 <- results(dds, alpha=0.05, independentFiltering=FALSE)

resOrdered_splike_vs_ACE2 <- res_splike_vs_ACE2[order(res_splike_vs_ACE2$padj),]
# resOrdered_splike_vs_beads
summary(res_splike_vs_ACE2)
```
```

````
### resultsNames(dds)
resLFC <- lfcShrink(dds, res = res_splike_vs_ACE2, coef = "sample_spike_vs_ACE2",
type="apeglm")

EnhancedVolcano(resLFC,
    lab = rownames(resLFC), x = 'log2FoldChange', pCutoff = 0.05, FCcutoff = 1,
    y = 'padj', title = "spike vs ACE2", subtitle = "(all peptides detected at least once in at least one
of the replicates included)")

```

Almost all peptides grouped around zero, meaning they are not changed in spike vs ACE2.

# Correlation matrix of spike vs ACE2

```{r scatter_spike_vs_ace2, message=FALSE, warning = FALSE, fig.width=10,fig.height=10}

CPM_df %>%
  select(pep, sample_id, CPM) %>%
  spread(key=sample_id, value=CPM) -> large_CPMdf

select(large_CPMdf, rownames(coldata)) %>%
  mutate_all(~replace(., is.na(.), 0)) %>%
  filter(rowSums(.)>3) -> large_CPMdf

ggpairs(large_CPMdf, diag = list(continuous = "blankDiag"),
        aes(alpha = 0.5, size = 0.001),
        lower = list(continuous = wrap("points", alpha = 0.3, size=0.1)),
        upper = list(continuous = wrap("cor", size = 5)))  +
    theme_bw(base_size = 11) +
  theme(axis.text.x = element_text(angle = 45, vjust = 1, hjust=1)) +
````

```
  scale_y_continuous(trans = "log2") +
  scale_x_continuous(trans = "log2")

```
```

These samples are indeed highly correlated.

### Differential enrichment of spike over nano

```
```{r spike_vs_nano, message=FALSE, warning = FALSE, echo=FALSE, fig.height=7}

metadata %>%
  filter(sample %in% c("spike", "nano"), round == 5) %>%
  select(sample_id, sample, round, rep)-> coldata

row.names(coldata) <- coldata$sample_id

select(large_df_5, row.names(coldata)) %>%
  filter(rowSums(.)>0) -> large_df_volcano

dds_raw <- DESeqDataSetFromMatrix(large_df_volcano,
                                  colData = coldata,
                                  design = ~ sample)

### # pre-filtering: ensure that at least X samples with a count of 5 or more, where X can be chosen as the sample size of the smallest group of samples (in this case 6 samples are the minimum number of samples in one comparison)
keep <- rowSums(counts(dds_raw) >= 1) >= 3
dds <- dds_raw[keep,]
dds$sample<- relevel(dds$sample, ref = "nano")
### run Deseq
```

```
dds <- DESeq(dds)

res_splike_vs_nano <- results(dds, alpha=0.05, independentFiltering=FALSE)

resOrdered_splike_vs_nano <- res_splike_vs_nano[order(res_splike_vs_nano$padj),]
### resOrdered_splike_vs_beads
summary(res_splike_vs_nano)

### resultsNames(dds)
resLFC_nano <- lfcShrink(dds, res = res_splike_vs_nano, coef = "sample_spike_vs_nano", type="apeglm")

EnhancedVolcano(resLFC_nano,
    lab = rownames(resLFC_nano), x = 'log2FoldChange', pCutoff = 0.05, FCcutoff = 1,
    y = 'padj', title = "spike vs nano")
```

There are some enriched peptides in the spike vs nano comparison.

# Correlation matrix of spike vs nano

```{r scatter_matrix_spike_vs_nano, message=FALSE, warning = FALSE, fig.width=10,fig.height=10}

CPM_df %>%
  select(pep, sample_id, CPM) %>%
  spread(key=sample_id, value=CPM) -> large_CPMdf

select(large_CPMdf, rownames(coldata)) %>%
  mutate_all(~replace(., is.na(.), 0)) %>%
```

````
  filter(rowSums(.)>3) -> large_CPMdf

ggpairs(large_CPMdf, diag = list(continuous = "blankDiag"),
        aes(alpha = 0.5, size = 0.001),
        lower = list(continuous = wrap("points", alpha = 0.3, size=0.1)),
        upper = list(continuous = wrap("cor", size = 5)))  +
    theme_bw(base_size = 11) +
  theme(axis.text.x = element_text(angle = 45, vjust = 1, hjust=1)) +
  scale_y_continuous(trans = "log2") +
  scale_x_continuous(trans = "log2")

```
````

Samples are again highly correlated, but a few significantly enriched peptides could be found in the volcano plots.

### Identifying intersections of spike over beads and spike over nano

````
```{r intersect_res, message=FALSE, warning = FALSE, echo=FALSE}

as.data.frame(resLFC_spike) %>%
  filter(log2FoldChange > 1 & padj < 0.05) %>%
  mutate(peptide = rownames(.))-> enriched_spike_vs_beads

as.data.frame(resLFC_nano) %>%
  filter(log2FoldChange > 1 & padj < 0.05) %>%
  mutate(peptide = rownames(.)) -> enriched_spike_vs_nano

enriched_spike_vs_beads %>%
  inner_join(enriched_spike_vs_nano, by="peptide") -> intersection_pos

list(spike_vs_beads = enriched_spike_vs_beads$peptide,
````

spike_vs_nano = enriched_spike_vs_nano$peptide) -> venn_list

ggVennDiagram(venn_list) +
  scale_fill_gradient(low="grey", high = "firebrick") +
  scale_color_manual(values=c(rep("black", 2))) +
  ggtitle("Significantly enriched peptides", subtitle = "(FC > 2, FDR < 5%)")

write_csv(enriched_spike_vs_beads, "enriched_spike_vs_beads.csv")
write_csv(enriched_spike_vs_nano, "enriched_spike_vs_nano.csv")

write_csv(as.data.frame(intersection_pos$peptide), "intersection_peptides.csv")

```
Almost all peptides enriched by spike over nano are also enriched by spike over beads.

```{r volcano_intersection, message=FALSE, warning = FALSE, echo=FALSE, fig.width=8, fig.height=7}

keyvals <- ifelse(rownames(resLFC_spike) %in% intersection_pos$peptide, 'orange', 'grey')
names(keyvals)[keyvals == 'orange'] <- 'spike/beads & spike/nano (n=37)'
names(keyvals)[keyvals == 'grey'] <- 'not enriched'

EnhancedVolcano(resLFC_spike,
    lab = rownames(resLFC_spike), selectLab = intersection_pos$peptide, x = 'log2FoldChange',
    y = 'padj', pCutoff = 0.05, colCustom=keyvals, title = "Spike vs beads")

keyvals <- ifelse(rownames(resLFC_nano) %in% intersection_pos$peptide, 'orange', 'grey')
names(keyvals)[keyvals == 'orange'] <- 'spike/beads & spike/nano (n=37)'
names(keyvals)[keyvals == 'grey'] <- 'not enriched'

EnhancedVolcano(resLFC_nano,

~~~
   lab = rownames(resLFC_nano), selectLab = intersection_pos$peptide, x = 'log2FoldChange',
   y = 'padj', pCutoff = 0.05, colCustom=keyvals, title = "Spike vs nano")


```
~~~

### How do interesting peptides behave across panning rounds?

Here, I plot the 37 peptides that are enriched by spike over beads and over nano

~~~
```{r lineplot,  message=FALSE, warning = FALSE, echo=FALSE}

CPM_df %>%
  filter(pep %in% intersection_pos$peptide) %>%
  group_by(pep, sample, round) %>%
  summarise(avgCPM = mean(CPM)) -> avgCPM_df

avgCPM_df$sample_id <- paste(avgCPM_df$sample, avgCPM_df$round, sep = "_R")

avgCPM_df %>%
  filter(sample == "spike") %>%
  mutate(group = "spike_pos_panning") -> avgCPM_pos

avgCPM_df %>%
  filter(sample == "beads") %>%
  mutate(group = "beads_panning")-> avgCPM_beads

avgCPM_df %>%
  filter(sample == "nano" | sample_id =="spike_R3") %>%
  mutate(group = "spike_nano_panning") %>%
  mutate(sample = "nano") %>%
  mutate(wrap="spike_pos_panning")-> avgCPM_neg
~~~

```
long_df <- bind_rows(avgCPM_pos, avgCPM_beads) %>%
  mutate(wrap = if_else(group == "beads_panning", "beads_panning", "spike_pos_panning"))
long_df$sample_id <- factor(long_df$sample_id, levels = unique(avgCPM_df$sample_id))

ggplot(avgCPM_neg, aes(x=round, y=avgCPM, group = pep, color = sample)) +
  geom_point(alpha=0.2) +
  geom_line(alpha = 0.2) +
  geom_point(data=long_df, alpha=0.2) +
  geom_line(data=long_df, alpha = 0.2) +
  facet_wrap(~wrap, ncol = 1) +
  scale_y_continuous(trans = "log2") +
  guides(colour = guide_legend(override.aes = list(alpha=1))) +
  theme_bw() +
  theme(text = element_text(size = 14))


ggplot(avgCPM_neg, aes(x=round, y=avgCPM, group = pep, color = sample)) +
  geom_point(alpha=0.2) +
  geom_line(alpha = 0.2) +
  geom_point(data=long_df, alpha=0.2) +
  geom_line(data=long_df, alpha = 0.2) +
  facet_wrap(~wrap, ncol = 1) +
  # scale_y_continuous(trans = "log2") +
  guides(colour = guide_legend(override.aes = list(alpha=1))) +
  theme_bw() +
  theme(text = element_text(size = 14))

### write.csv(long_df, "lineplot_data.csv")
### ggsave("lineplot_linear.pdf")
```

**Mesoscale Discovery (MSD) assay**

In general, the V-PLEX COVID-19 ACE2 Neutralization Kits quantitatively measure chemical/biological samples that block the binding of ACE2 to its cognate ligands. Plates are provided with antigens on spots in the wells of a 96-well plate. Blocking samples bind to antigens on the spots, and human ACE2 protein conjugated with MSD SULFO-TAG™ is used for detection. The plate is read on an MSD® instrument, which measures the light emitted from the MSD SULFO-TAG. To prepare blocker A solution, follow the preparation procedure in the product insert provided with the Blocker A Kit to prepare the Blocker A solution. To prepare wash buffer, MSD provides 100 mL of Wash Buffer as a 20X stock solution. Dilute the stock solution before use. PBS + 0.05% Tween-20 can be used as an alternative to MSD Wash Buffer. For one plate, combine 15 mL of MSD Wash Buffer (20X) and 285 mL of deionized water. Use Diluent 100 as assay diluent. To prepare plate, remove the plate from its packaging. Add 150 µL/well of Blocker A solution to the plate. Seal the plate with an adhesive plate seal and incubate at room temperature with shaking (~700 rpm) for 30 minutes. During this time, prepare samples and calibrators. To prepare compound samples, prepare the samples by diluting with Diluent 100. To make an intermediate 1:4 dilution in a 96-well plate, combine 10 µL of sample, and 30 µL of Diluent 100. For compound solubility, final DMSO concentration is 1%. To prepare calibrator, the kits include a calibration reagent, which is used to establish a calibration curve in the assay. After the Blocker A incubation step, wash the plate 3 times with at least 150 µL/well of 1X MSD Wash buffer. Add 25 µL/well of diluted samples and calibrator to the plate. Seal the plate with an adhesive plate seal and incubate at 37 °C with shaking (~700 rpm) for annotated time. Do not aspirate or wash the plate prior to addition of detection solution. During this time, prepare the ACE2 detection solution. To prepare a 1X solution of SULFO-TAG Human ACE2 Protein, combine 2,985 µL of Diluent 100 and 15 µL of 200X SULFO-TAG Human ACE2 Protein. After the sample and calibrator incubation, do not aspirate or wash the plate prior to addition of detection solution. Add 25 µL/well of 1X SULFO-TAG Human ACE2 Protein detection solution to the plate. Seal the plate with an adhesive plate seal and incubate at room temperature with shaking (~700 rpm) for 1 hour. After the detect incubation step, wash the plate 3 times with at least 150 µL/well of 1X MSD Wash buffer. MSD provides MSD GOLD Read Buffer B ready for use. Do not dilute. Add 150 µL/well of the MSD GOLD Read Buffer B. Read the plate on the MSD instrument. No incubation in read buffer is required before reading the plate. Read plate immediately after adding read buffer. Do not shake the plate after adding read buffer. Results can be reported as percent inhibition, calculated using the equation below. Highly positive samples show high percent inhibition whereas negative or low samples show low percent inhibition.

$$\%\,Inhibition = 1 - \frac{Average\ Sample\ ECL\ Signal}{Average\ ECL\ signal\ of\ Calibrator\ 8\ (Diluent\ only)}\ x\ 100$$

**Biolayer Interferometry**

All reactions were run on an Octet RED96 at 30 °C, and samples were run in 1× PBS with 0.1% BSA and 0.05% Tween 20 (Octet buffer). Macrocycle compounds, ACE2 and Nb70, were assessed for binding to biotinylated antigens using streptavidin biosensors (Sartorius/ForteBio). Antigen was loaded at annotated concentrations. Tips were then washed and baselined in wells containing only Octet buffer. Samples were then loaded for binding association step followed up with a dissociation step. A control well with loaded antigen but that was associated in a well containing only 200 µl of Octet buffer was used as a baseline subtraction for data analysis. Association and dissociation binding curves were fit in Octet System Data Analysis Software version 9.0.0.15 using a 1:2 bivalent model to determine apparent Kd. Averages of Kd fold-change values from at least two independent experiments are reported to two significant. To estimate measurement error, we computed the standard deviation for each sample.

**In-gel fluorescence protein labeling of cyclic peptides to spike**

For direct labeling, Omicron BA2 spike protein (0.25 uM) was incubated with the indicated concentrations of each chemical probe for annotated time at 37 °C. For competition labeling, ACE2 or BSA was doped in with cyclic peptides for binding to spike at the indicated concentration. After protein labeling, 20 µL of resulting mixtures were added 2.16 µL of freshly prepared click mix (0.5 µL of 50 mM CuSO4 in $H_2O$, 1.16 µL of 100 mM BTTAA in DMSO, and 0.5 µL of 1 mM N3-TAMRA in DMSO) and 1.16 µL of 300 mM sodium ascorbate solution in $H_2O$. After incubation for 30 min at 37 ºC, 8 µL of 4X SDS loading buffer was added and samples were boiled at 100 ºC for 5 min. The samples treated with fluorescent probe was directly subjected by 4X SDS loading buffer without click reaction and boiled at the same condition with alkyne probes. Then, samples were analyzed by SDS-PAGE (12%) running at 120 V in an electrophoresis chamber under the ambient temperature. In-gel fluorescence was visualized using a GE Typhoon FLA 9000 (GE Healthcare, Pittsburgh, PA) followed by staining using Coomassie.

**Determination of neutralization activity using authentic anti-viral assays**

The neutralization capacity of compounds was assessed in titration experiments as described in previous studies.[2–4] The SARS-CoV-2 variants B.1.1.529.1 (omicron BA.1) and B.1.1.529.5 (omicron BA.5) were isolated from nasopharyngeal swabs of RT-PCR- and sequencing-confirmed SARS-CoV-2-positive patients. B.1.1.529.2 (omicron BA.2) was kindly provided by Dr. Marie-Anne Rameix-Welti from Institut Pasteur via European Virus Archive Global (EVAg) platform. Each variant was amplified in Calu-3, lung epithelial cells and viral titers of stocks were determined using the Median Tissue Culture Infectious Dose ($TCID_{50}$) assay.

To evaluate neutralization capacity, three-fold serial dilutions of the test compounds were prepared in PBS containing 3% DMSO and incubated with $5x10^4$ $TCID_{50}$ of SARS-CoV-2 omicron BA.1, BA.2 or BA.5 for 1 hour at 37°C. 1/3 of the mixture was inoculated to VeroE6 cells (1.5x $10^4$ cells/well in 96 well plate, MOI=1.1), and viral replication was determined at 24h post infection using immunostaining of viral nucleocapsid. Cells were fixed with 5% formaldehyde and permeablized with 0.2% Triton X-100. After blocking with 2% skim milk, cells were incubated with rabbit monoclonal anti-SARS-CoV-2 nucleocapsid antibody (Sinobiological, Beijing, China) followed by a secondary incubation with anti-rabbit HRP-conjugated antibody (Merck, Darmstadt, Germany) for 1 hour at 37°C. The signal was developed using the KPL SureBlueTM 3,3',5,5'-tetramethylbenzidine peroxidase substrate (Seracare, Milford, MA, USA) after 5 min incubation, and the reaction was stopped by the addition of 0.5 M sulfuric acid. Absorbance was measured at 450 nm with reference wavelength 620 nm on a Tecan Sunrise plate reader (Tecan, Männedorf, Switzerland). Values were normalized to solvent control (1% DMSO) with infection (100% infection) and without infection (0% infection, assay background). $EC_{50}$ values were calculated using non-linear regression dose response analysis by GraphPad Prism version 8.4.3 (GraphPad Software, San Diego CA, USA).

**Antiviral cellular jump dilution assay**

Three-fold serial dilutions of the test compounds were prepared in PBS containing 3% DMSO and incubated with $5x10^5$ $TCID_{50}$ of SARS-CoV-2 omicron BA.1 for 1 hour at 37°C. The mixture was then diluted 10-fold with 3% DMSO containing PBS, afterward inoculated to VeroE6 cells. The virus replication was determined at 24h post infection using immunostaining of viral nucleocapsid as described above.

## 3. Chemical Synthesis

**General Synthesis Methods:** Chromatographic separations were performed by manual flash chromatography unless otherwise specified. Silica gel 60 (70–230 mesh, Merck) was used for manual column chromatography. Commercial plates (F254, 0.25-mm thickness; Merck) were used for analytical thin-layer chromatography to follow the progress of reactions. Unless otherwise specified, $^{1}H$ NMR spectra and $^{13}C$ NMR spectra were obtained on a Varian Mercury 400-MHz console connected with an Oxford NMR AS400 actively shielded magnet or a 500-MHz Varian Inova spectrometer at room temperature. Chemical shifts for $^{1}H$ or $^{13}C$ are given in ppm ($\delta$) relative to tetramethylsilane as an internal standard. Mass spectra (m/z) of chemical compounds were recorded on a Finnigan LTQ mass spectrometer (Thermo Scientific), an ACQUITY UPLC SQ Detector 2 system (Waters) or a 1260 Infinity LC (Agilent) connected with an InfinityLab MS detector. Reverse-phase HPLC (RP-HPLC) purifications were performed using a 1260 Infinity HPLC equipped with a semi-prep C18 column (5 µm, C18, 100 Å, 250 × 10mm liquid chromatography column; Phenomenex) eluted over a linear gradient from 95% solvent A (water and 0.1% trifluoroacetic acid) to 100% solvent B (acetonitrile and 0.1% trifluoroacetic acid).

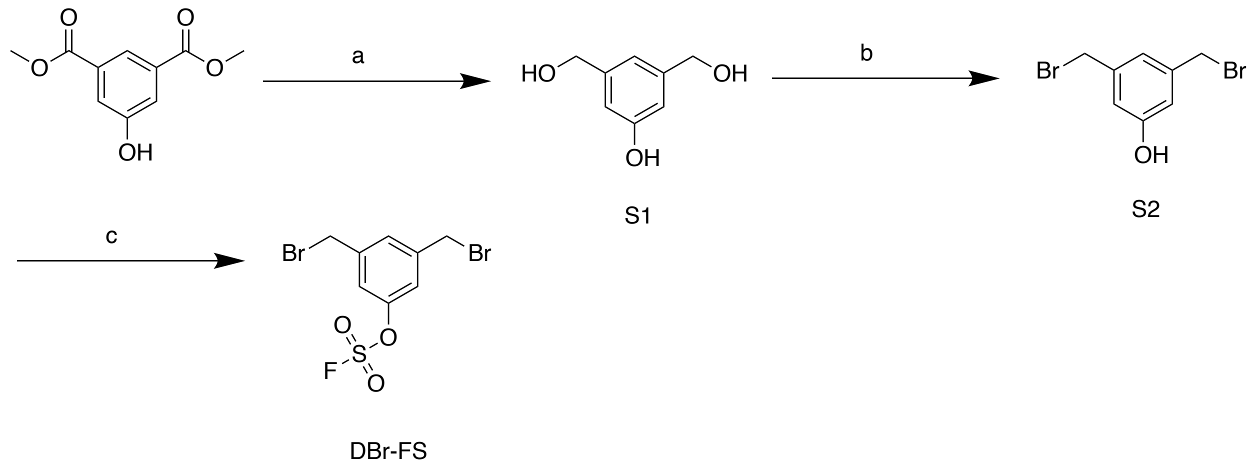


**Scheme S1:** Synthesis of DBr-FS (dibromo-aryl fluorosulfate). Reagent and conditions: (a) aluminum (III) lithium hydride, THF; (b) tribromophosphane, THF; (c) (((4-acetamidophenyl)azanediyl)bis(oxy))disulfonyl fluoride, 2,3,4,6,7,8,9,10-octahydropyrimido[1,2-a]azepine, THF

**Synthesis of DBr-FS**

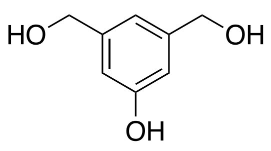

S1

(5-hydroxy-1,3-phenylene)dimethanol (**S1**)

To a stirred solution of aluminum(III) lithium hydride (3.61 g, 95.2 mmol) in THF (95 mL) was added dimethyl 5-hydroxyisophthalate (5 g, 23.8 mmol) at 0 °C under argon. After being stirred for 3 hrs at room temperature, the reaction mixture was acidified with 100mL 10% $H_2SO_4$. Solid was filtered. The aqueous mixture was extracted with ethyl acetate, and washed with brine. The organic layer was dried over $Na_2SO_4$ and concentrated in vacuo. The resulting residue was purified by flash column chromatography on silica gel (EtOAc/*n*-hexane = 1:1.5) to afford 3.146g (85.8%) **S1**

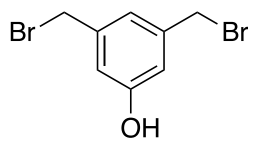

S2

3,5-bis(bromomethyl)phenol (**S2**)

To a stirred solution of **S1** (3.14 g, 20.4 mmol) in THF (18 mL) was added tribromophosphane (17.9 g, 66.2 mmol) in 2.63 mL of THF over 30 minutes. After being stirred for 72 h at 40 °C, the reaction mixture was washed with saturated sodium bicarbonate, and extracted with ethyl acetate three times. The organic layer was washed with brine and dried over $Na_2SO_4$ and concentrated in vacuo. The resulting residue was purified by flash column chromatography on silica gel using a gradient EtOAc (0-30%) in *n*-hexane to afford 1.22 g (21.4%) **S2**

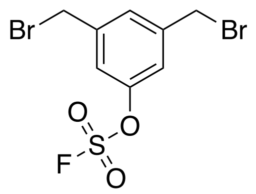

DBr-FS

3,5-bis(bromomethyl)phenyl sulfurofluoridate (**DBr-FS**)

To a one-dram vial containing **S2** (1.096 g, 3.915 mmol) and AISF ((((4-acetamidophenyl)azanediyl)bis(oxy))disulfonyl fluoride) (1.627 g, 4.698 mmol, 1.2 equiv.) was added tetrahydrofuran (20 mL) followed by 1,8 diazabicyclo [5.4.0]undec-7-ene (760 μL, 5.089 mmol, 1.3 equiv.) over a period of 30 seconds. The reaction mixture was stirred at room temperature for 10 minutes and then diluted with ethyl acetate or ether and washed with either 0.5 N KHSO4 or 0.5 N HCl (2x) and brine (1x). The combined organic fraction was dried with anhydrous sodium sulfate and concentrated under reduced pressure. The crude residue was purified by silica gel flash chromatography to afford 751 mg (53%) **DBr-FS** (3,5-bis(bromomethyl)phenyl sulfurofluoridate) as a white solid: $^1$H NMR (500 MHz, $CDCl_3$) δ 7.47 – 7.42 (s, 1H), 7.31 – 7.26 (s, 2H), 4.44-4.42 (d, 4H); $^{19}$F NMR (470 MHz, $CDCl_3$) δ 38.3 (s, 1F). M.W. calc. 360.85/362.85 $[M+1]^+$, found 360.9/362.9 $[M+1]^+$

**Synthesis of macrocyclic hits**

*Solid-Phase Peptide Synthesis (SPPS)*

Peptides were prepared using traditional SPPS methods. Peptides were synthesized on rink amide resin (30 mg, loading 100-200 mesh, 0.64 meq/g, 1% DVB) (Chem-Impex Int'L INC, Cat. # 02900) using standard "coupling-deprotection-coupling" Fmoc chemistry. A Syro II (Biotage) semi-automated parallel peptide synthesizer with standard reactor block with 2 mL reaction vessel (PP-Reactor, 2 mL, with PE Frit, Cat. # V020PE051) (2mL plunger, Cat. # V020ST020) were used for synthesis. General procedures for linear peptide synthesis follow the general procedure A (Fmoc deprotection), general procedure B (amide coupling), and general procedure C (washing steps). For the last step chloroacetic acid coupling, 5eq chloroacetic acid, 5 eq HBTU, and 5eq collidine were pre-mixed for activation before manually loading onto the peptide resin for reactions. The procedure was repeated twice. After the last step coupling, 6 times washing using DMF, followed by with 6 times washing using DCM, was performed before cleavage. Peptides were cleaved from resin using a mixture of 95% trifluoroacetic acid (Chem-Impex Int'L INC, Cat. # 00289), 2.5 % triisopropylsilane (Sigma Aldrich, Cat # 233781), 2.5% MilliQ water for 2 hours at RT, with occasional shaking. The cleavage mixture was drained and collected. The resin was then washed with additional cleavage mixture, drained, and collected. TFA was concentrated through evaporation with air stream in a ventilated hood. The residual cleavage mixture was precipitated in diethyl ether and allowed to cool at -20°C before centrifuge. The ether was then decanted, and this process was completed three times. After the third ether wash, the residual ether was allowed to evaporate, and the compounds were dissolved in DMSO into 2mM stocks for next step cyclization.

*General Procedure A*: Fmoc Deprotection. Peptides were deprotected using a 20% piperidine (TCI, Cat # 203-642-1) in DMF solution. Peptides were deprotected for 15 min at RT two times.

*General Procedure B*: Amide bond coupling for amino acids. The coupling reagent (2-(1Hbenzotriazol-1-yl)-1,1,3,3-tetramethyluronium hexafluorophosphate (HBTU) (Peptides International, Cat # KHB-1065-PI) was pre-dissolved in DMF. The coupling reagent 2,4,6-collidine (Alfa Aesar, Cat # A11058) was pre-dissolved in DMF. All coupling reagent amino acids were predissolved in DMF. 5 equivalents of amino acid and 5 equivalents HBTU, and 10.0 equivalents of 2,4,6-collidine were preactivated and added to the reaction vessel using minimal DMF. The coupling reaction was allowed to react for 2x20 min at RT with contiguous shaking using the peptide synthesizer.

*General Procedure C*: Washing Step. The reaction vessel was drained followed by addition of DMF for washing, this process was repeated five more times before next "deprotection-coupling-washing" cycle.

*Peptide Cyclization*

For 37 hit peptides selected based on sequencing analysis, ~6 mg of lyophilized linear peptides was dissolved in 500 uL DMSO. Based on the approximate molecular weight of the peptides (1000-1500 Da), an average M.W. of 1200 Da was used to estimate the total amount (1eq.) of crude peptides in 500 uL DMSO. 1.5 eq. of dibromo-$OSO_2F$ linker in DMSO was doped into the 500 uL DMSO dissolved peptide, with 2.5 eq. of DIPEA as the base, in the reaction mixture for macrocyclization of linear peptides into cyclic peptides. Reaction was incubated at 37 C° for 1 hour, shaking at 300 rpm. Cyclization of compounds was monitored and product yield was quantified by the LCMS.

**4. Compound Table**

37 Crude hits

| Compound name | Calc. MW $[M+1]^+$ | Found MW $[M+1]^+$ |
|---|---|---|
| CP_SW1 | 1364.52 | 1364.8 |
| CP_SW2 | 1419.5 | 1418.8 |
| CP_SW3 | 1511.59 | 1510.9 |
| CP_SW4 | 1536.54 | 1536.8 |
| CP_SW5 | 1337.51 | 1337.8 |
| CP_SW6 | 1589.56 | 1589.9 |
| CP_SW7 | 1502.54 | 1501.8 |
| CP_SW8 | 1498.59 | 1498.8 |
| CP_SW9 | 1550.51 | 1550.2 |
| CP_SW10 | 1502.56 | 1501.8 |
| CP_SW11 | 1555.57 | 1554.8 |
| CP_SW12 | 1515.58 | 1514.8 |
| CP_SW13 | 1416.52 | 1415.9 |
| CP_SW14 | 1392.58 | 1391.9 |
| CP_SW15 | 1416.52 | 1415.8 |
| CP_SW16 | 1463.59 | 1462.9 |
| CP_SW17 | 1355.48 | 1354.8 |
| CP_SW18 | 1590.59 | 1589.8 |
| CP_SW19 | 1390.51 | 1389.8 |
| CP_SW20 | 1507.58 | 1506.8 |
| CP_SW21 | 1510.61 | 1509.9 |
| CP_SW22 | 1463.58 | 1462.9 |
| CP_SW23 | 1526.55 | 1525.8 |
| CP_SW24 | 1444.55 | 1443.8 |
| CP_SW25 | 1445.53 | 1444.8 |
| CP_SW26 | 1347.45 | 1346.8 |

| | | |
|---|---|---|
| CP_SW27 | 1522.50 | 1521.9 |
| CP_SW28 | 1509.57 | 1508.9 |
| CP_SW29 | 1426.55 | 1425.9 |
| CP_SW30 | 1426.51 | 1425.9 |
| CP_SW31 | 1515.54 | 1514.8 |
| CP_SW32 | 1663.60 | 1662.9 |
| CP_SW33 | 1443.46 | 1442.8 |
| CP_SW34 | 1640.65 | 1640.9 |
| CP_SW35 | 1552.54 | 1551.8 |
| CP_SW36 | 1610.54 | 1609.9 |
| CP_SW37 | 1487.56 | 1486.9 |

17 purified hits (alkyne versions)

| Compound name | Calc. MW $[M+1]^+$ | Found MW $[M+1]^+$ |
|---|---|---|
| CP_SW2A | 1443.49 | 1443.0 |
| CP_SW3A | 1534.61 | 1534.8 |
| CP_SW4A | 1560.54 | 1560.8 |
| CP_SW5A | 1361.51 | 1360.9 |
| CP_SW6A | 1613.56 | 1612.8 |
| CP_SW10A | 1526.56 | 1526.8 |
| CP_SW11A | 1579.57 | 1580.3 |
| CP_SW12A | 1539.58 | 1539.8 |
| CP_SW15A | 1440.52 | 1440.8 |
| CP_SW16A | 1487.59 | 1487.9 |
| CP_SW17A | 1379.48 | 1378.8 |
| CP_SW21A | 1534.61 | 1533.9 |
| CP_SW22A | 1487.58 | 1487.9 |
| CP_SW23A | 1550.55 | 1549.8 |
| CP_SW31A | 1539.54 | 1538.9 |

| | | |
|---|---|---|
| CP_SW33A | 1467.46 | 1466.8 |
| CP_SW34A | 1664.65 | 1663.9 |
| CP_SW35A | 1576.54 | 1576.8 |
| CP_SW36A | 1634.54 | 1633.8 |
| CP_SW25A | 1469.53 | 1468.9 |

Other compounds (alanine mutants, negative control compounds, linker)

| Compound name | Calc. MW $[M+1]^+$ | Found MW $[M+1]^+$ |
|---|---|---|
| CP_SW3A_A1 | 1477.59 | 1476.9 |
| CP_SW3A_A2 | 1506.58 | 1506.9 |
| CP_SW3A_A3 | 1477.59 | 1477.9 |
| CP_SW3A_A4 | 1490.62 | 1490.9 |
| CP_SW3A_A5 | 1492.56 | 1492.9 |
| CP_SW3A_A6 | 1442.58 | 1442.9 |
| CP_SW3A_A7 | 1506.58 | 1506.9 |
| CP_SW3A_A8 | 1492.56 | 1492.8 |
| CP_SW3A_Bz | 1452.66 | 1453.0 |
| CP_SW3A_DS | 1332.6 | 1332.9 |
| CP_SW3A_S1 | 1534.61 | 1534.8 |
| CP_SW3A_S2 | 1534.61 | 1534.8 |
| DBr-FS (linker) | 360.85/362.85 | 360.9/362.9 |

## 5. LCMS of linker DBr-FS and cyclic chemical probes

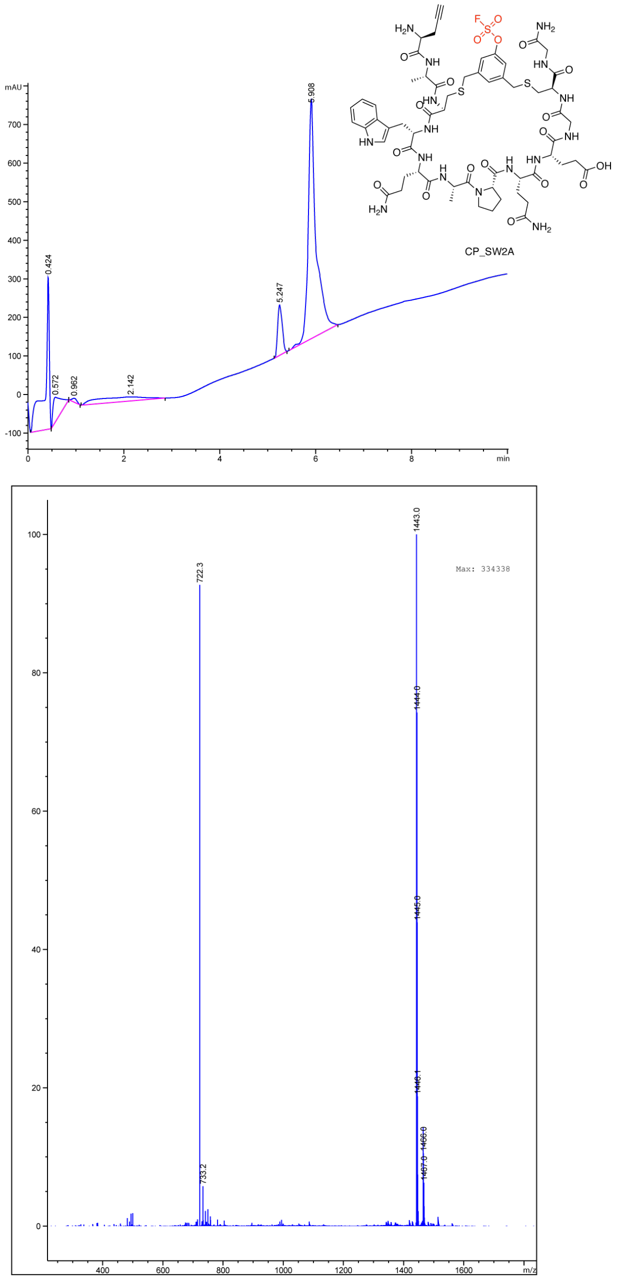


LCMS traces of CP-SW2A (A) LC trace (B) MS

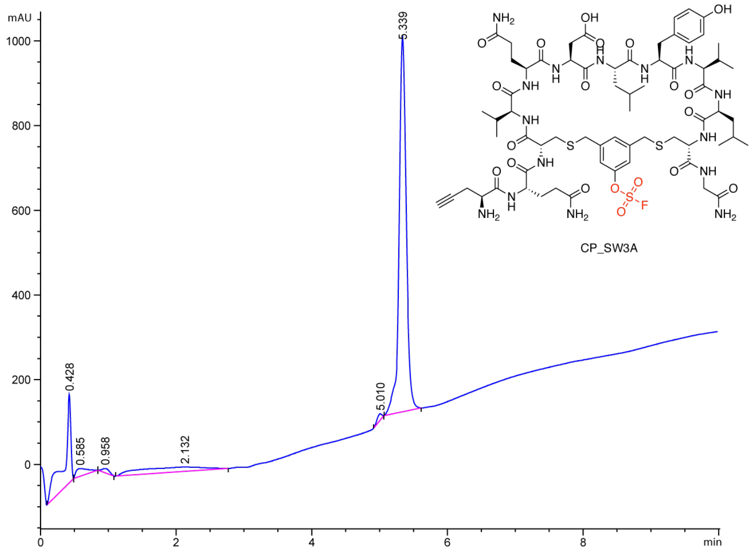


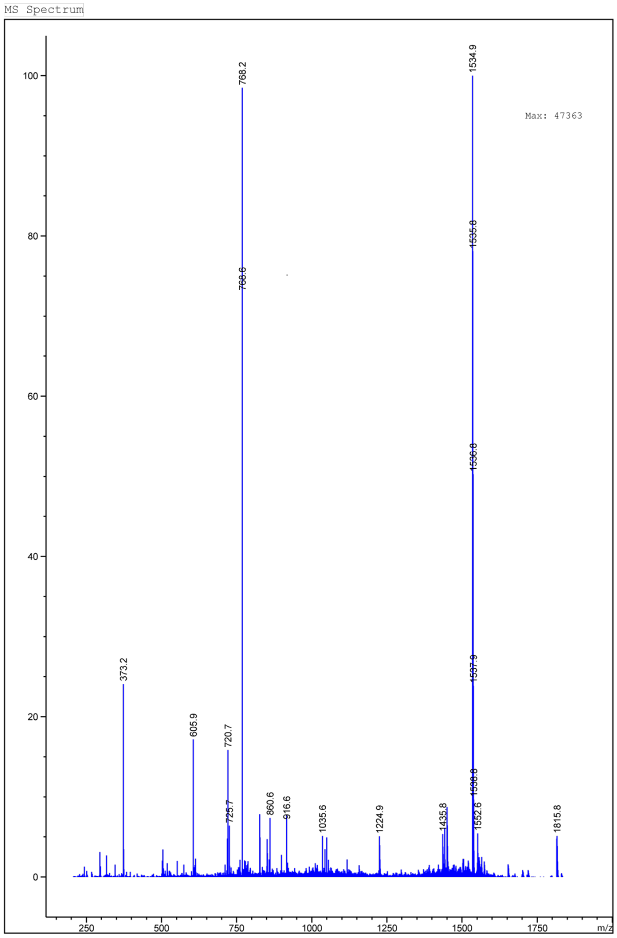


LCMS traces of CP-SW3A. (A) LC trace (B) MS

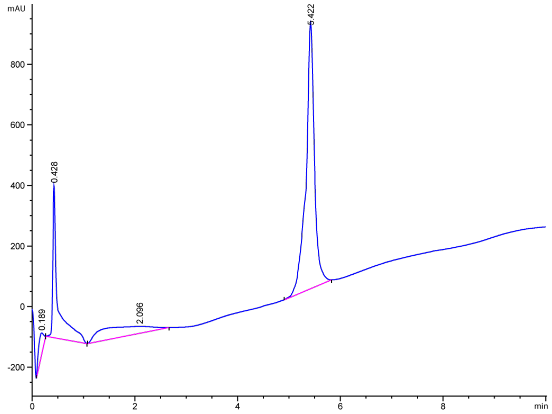


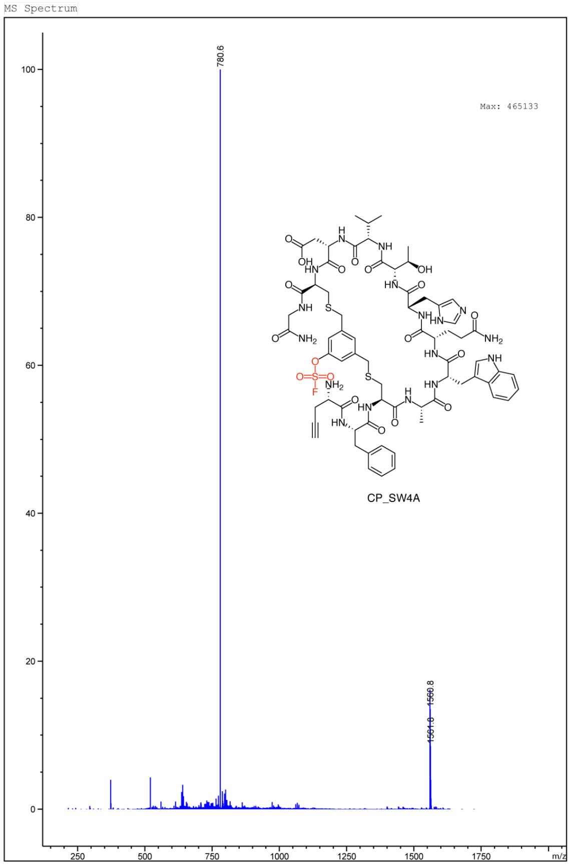


LCMS traces of CP-SW4A. (A) LC trace (B) MS

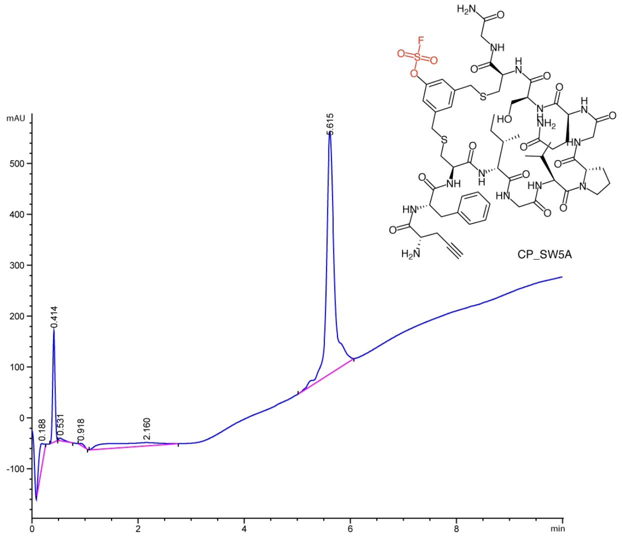


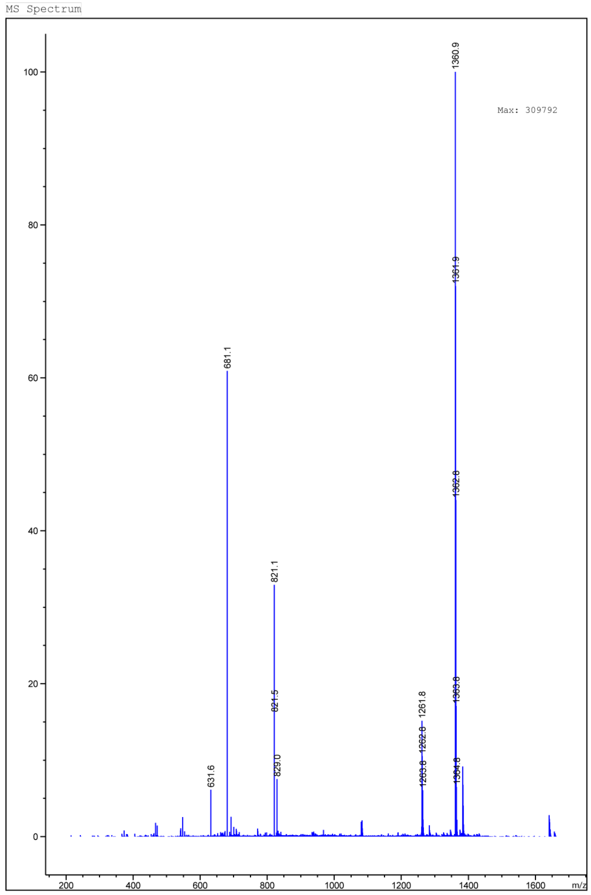


LCMS traces of CP-SW5A. (A) LC trace (B) MS

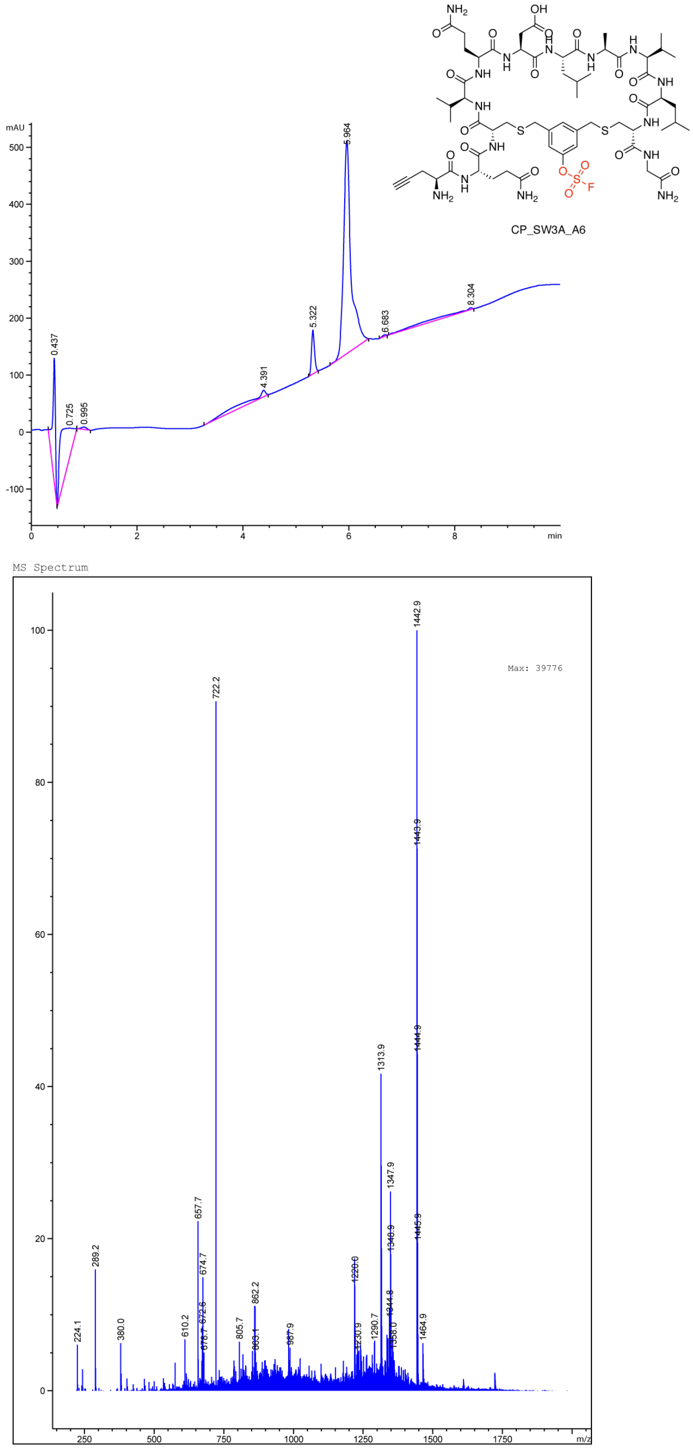


LCMS traces of CP-SW6A. (A) LC trace (B) MS

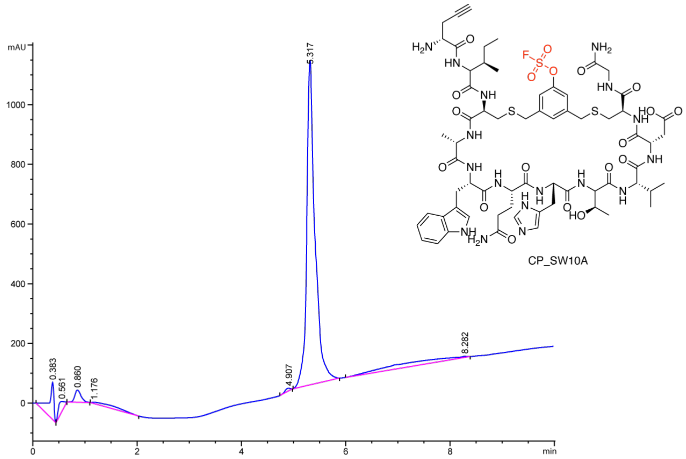


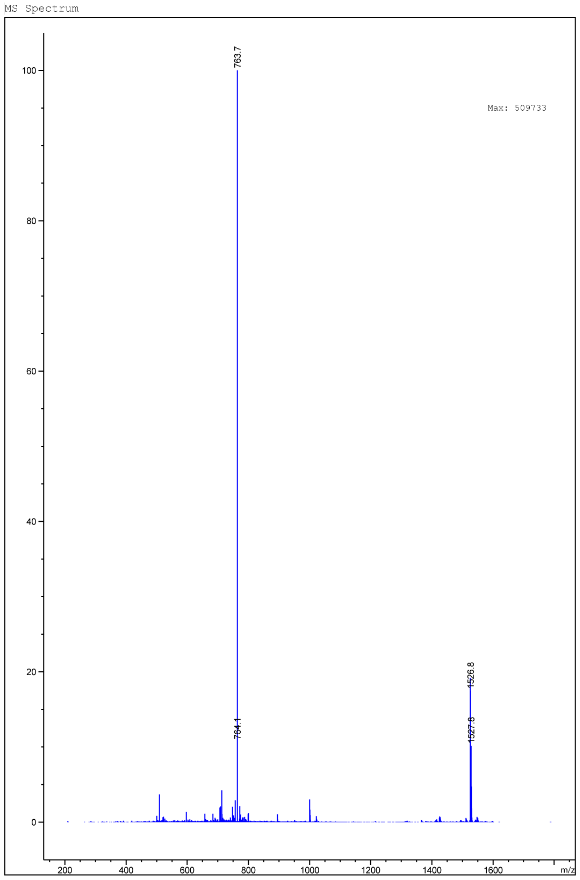


LCMS traces of CP-SW10A. (A) LC trace (B) MS

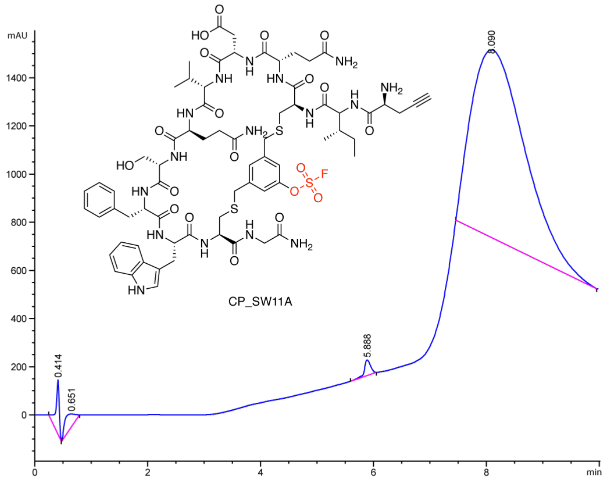


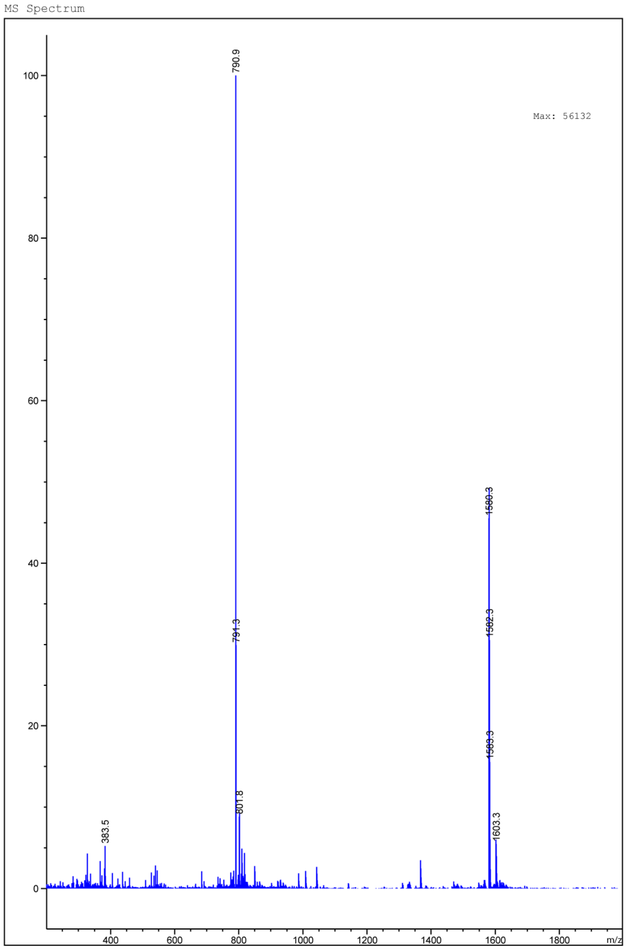


LCMS traces of CP-SW11A. (A) LC trace (B) MS

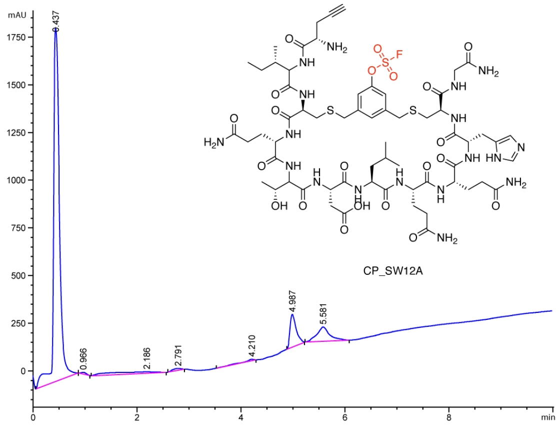


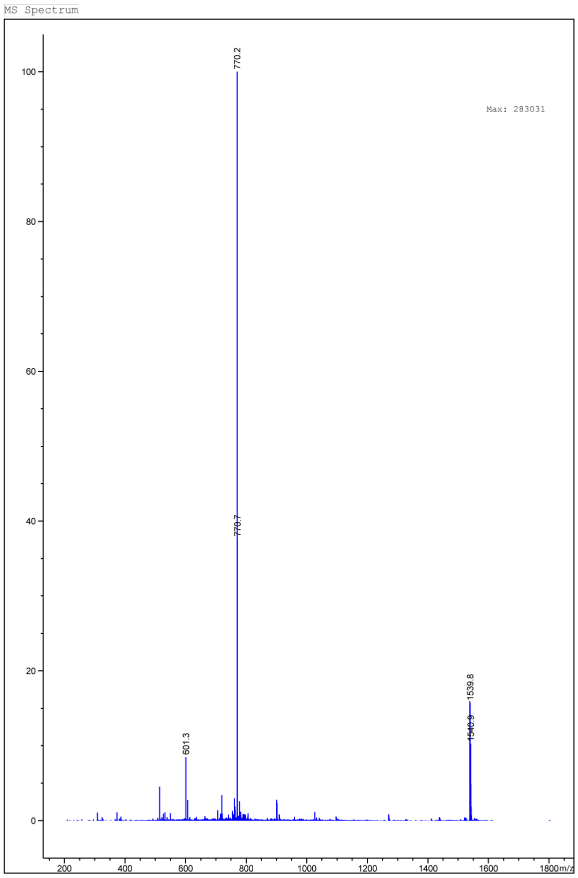


LCMS traces of CP-SW12A. (A) LC trace (B) MS

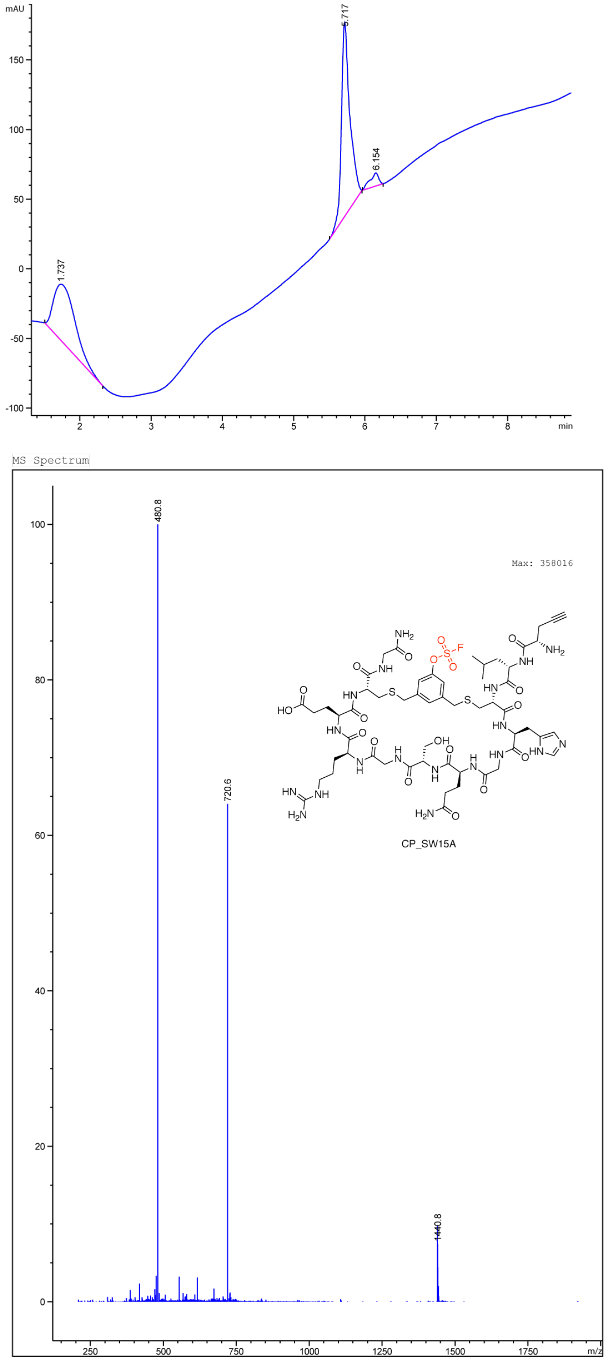


LCMS traces of CP-SW15A. (A) LC trace (B) MS

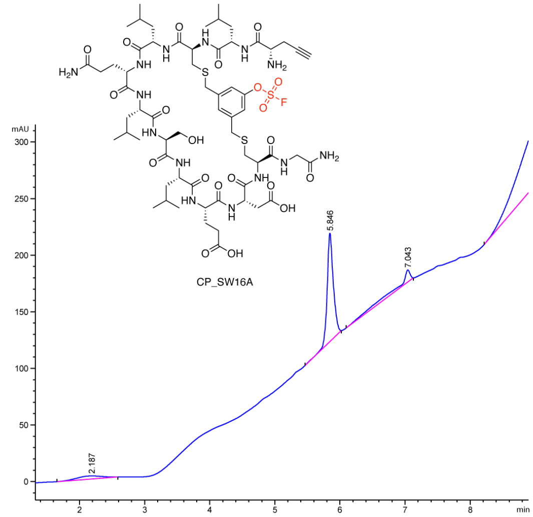


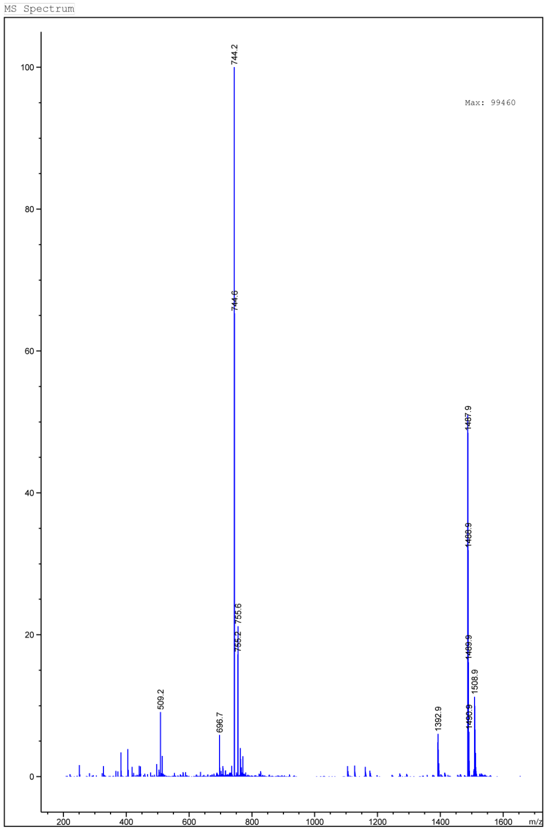


LCMS traces of CP-SW16A. (A) LC trace (B) MS

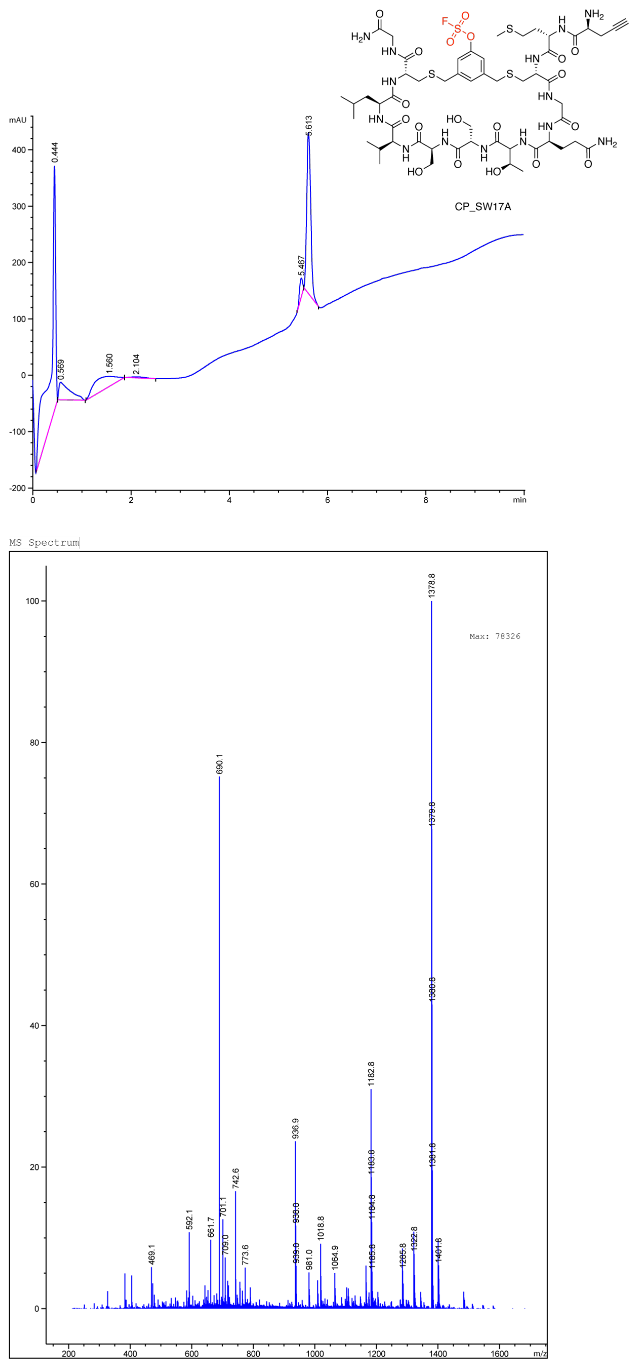


LCMS traces of CP-SW17A. (A) LC trace (B) MS

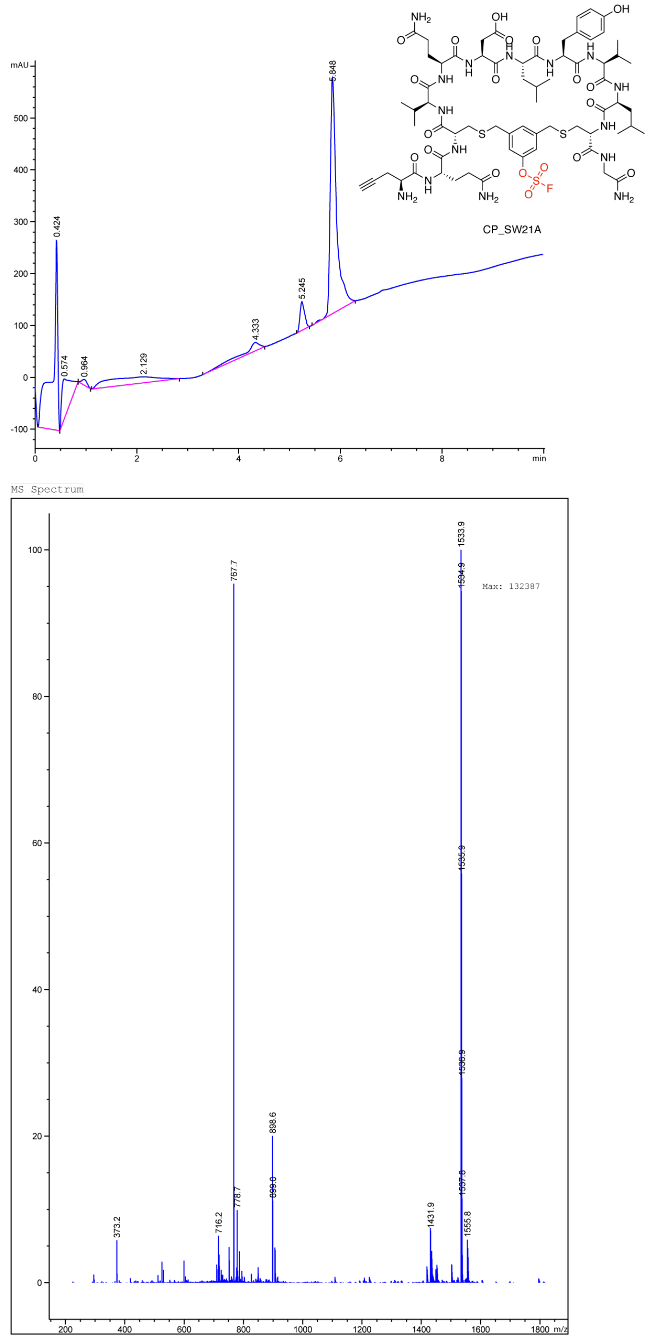


LCMS traces of CP-SW21A. (A) LC trace (B) MS

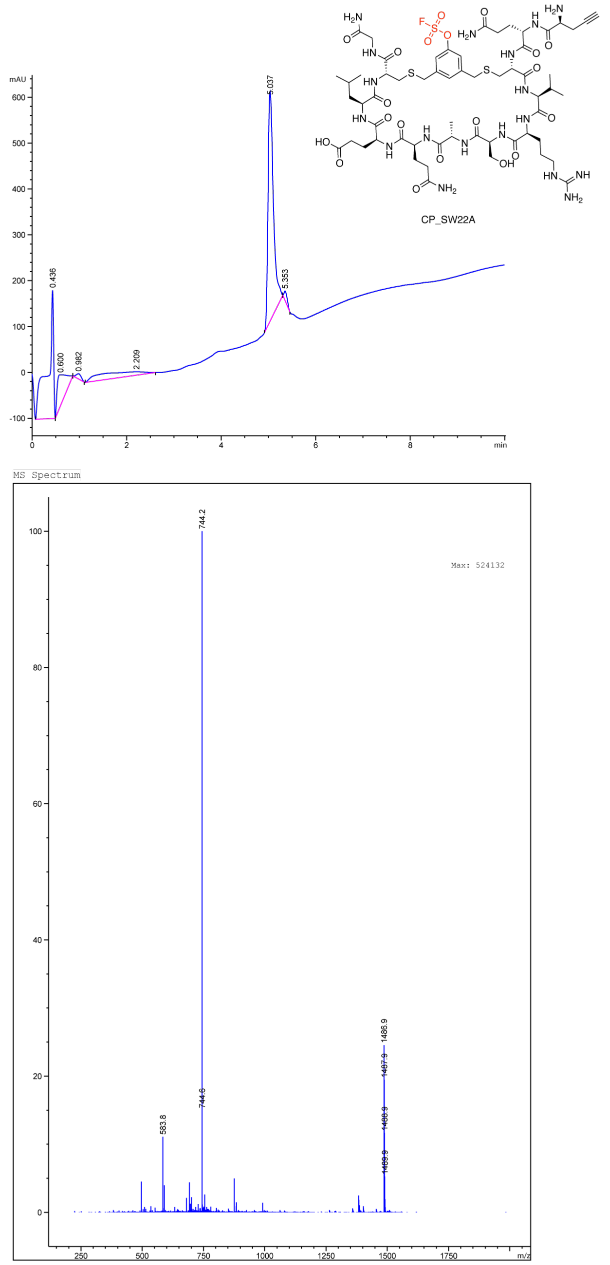


LCMS traces of CP-SW22A. (A) LC trace (B) MS

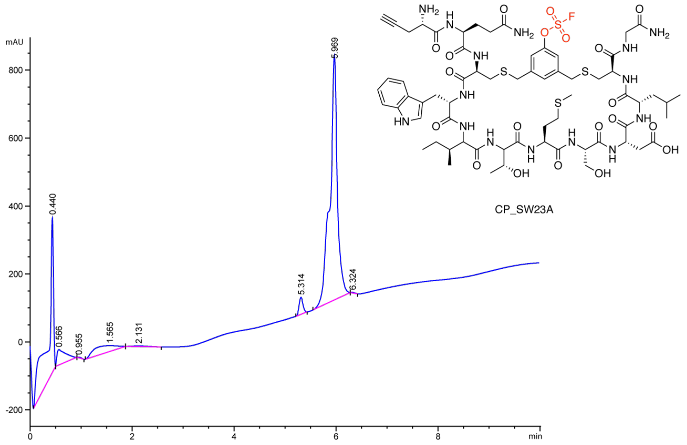


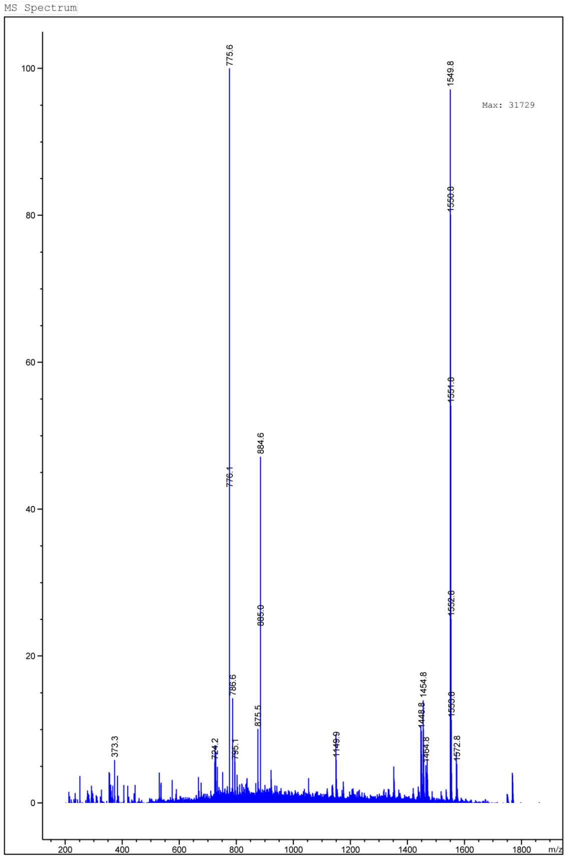


LCMS traces of CP-SW23A. (A) LC trace (B) MS

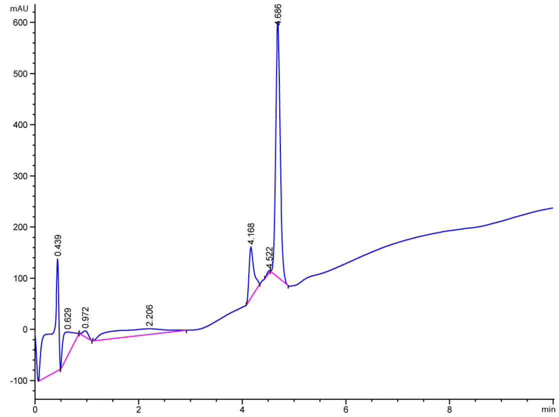

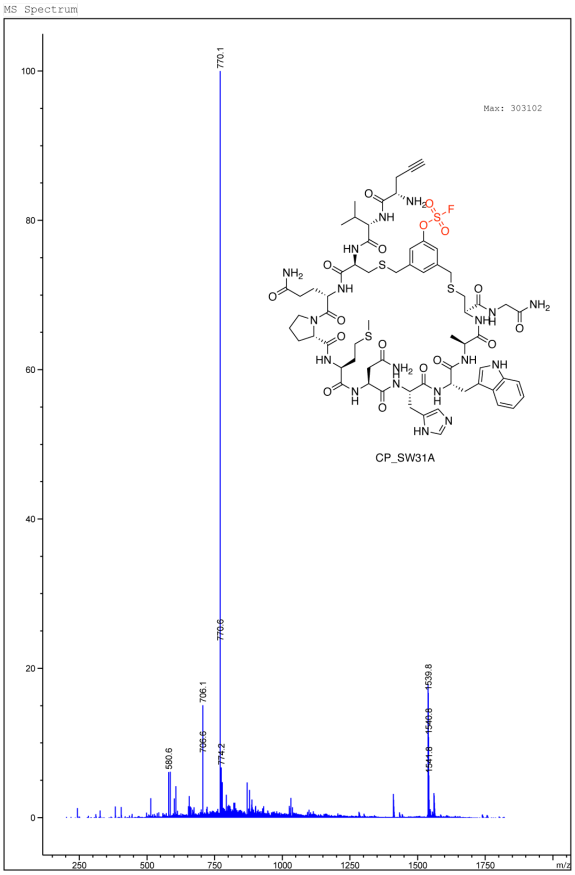


LCMS traces of CP-SW31A. (A) LC trace (B) MS

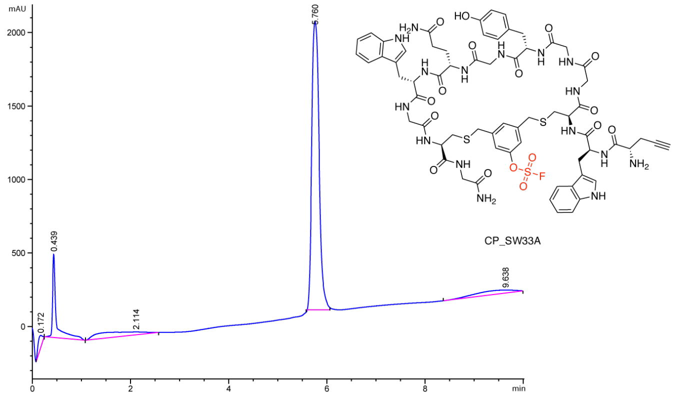


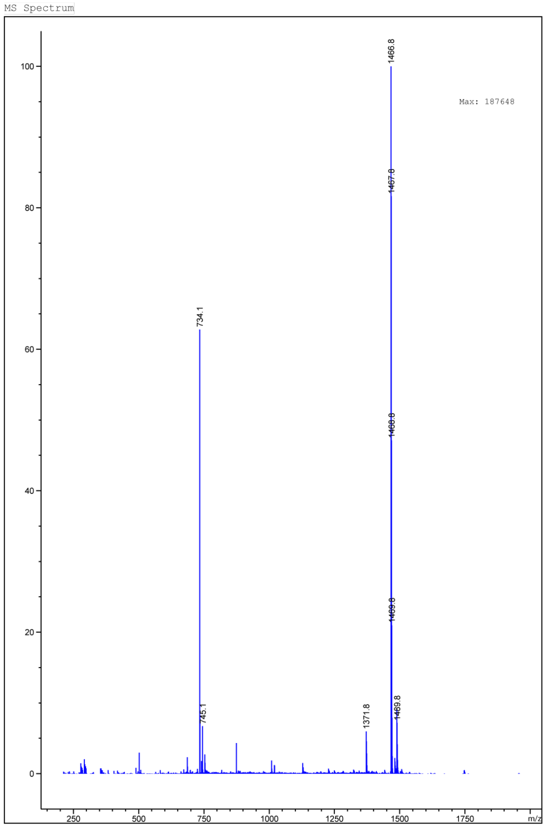


LCMS traces of CP-SW33A. (A) LC trace (B) MS

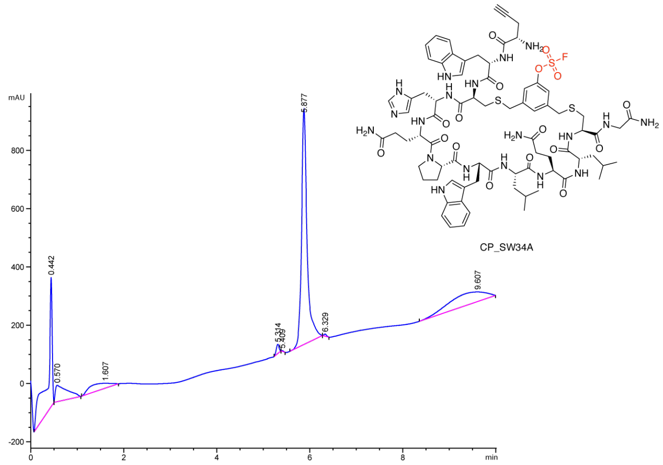


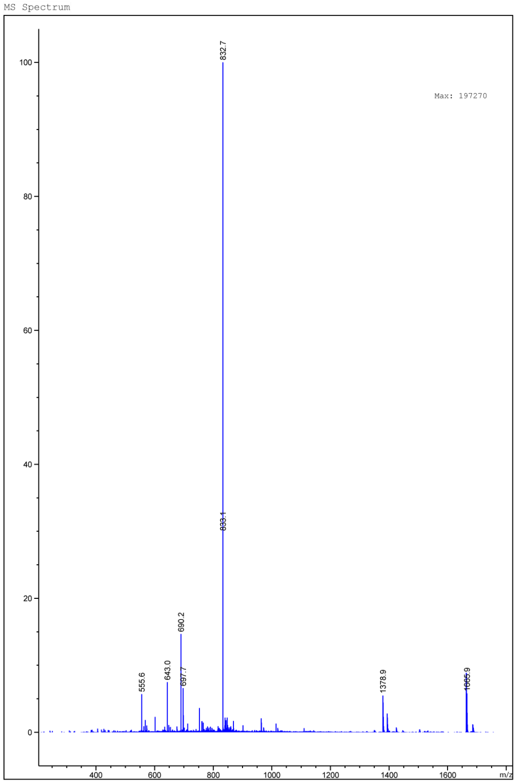


LCMS traces of CP-SW34A. (A) LC trace (B) MS

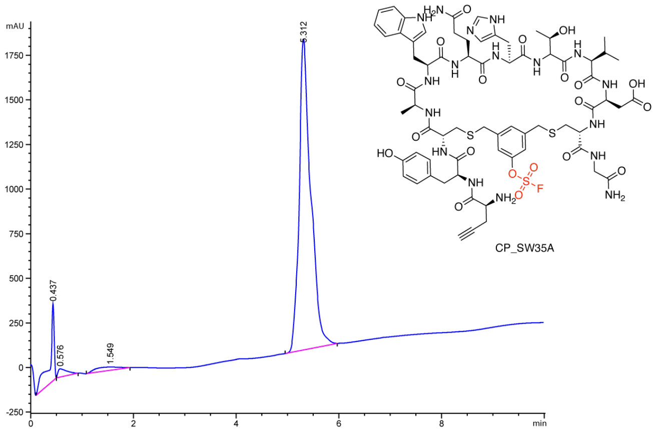


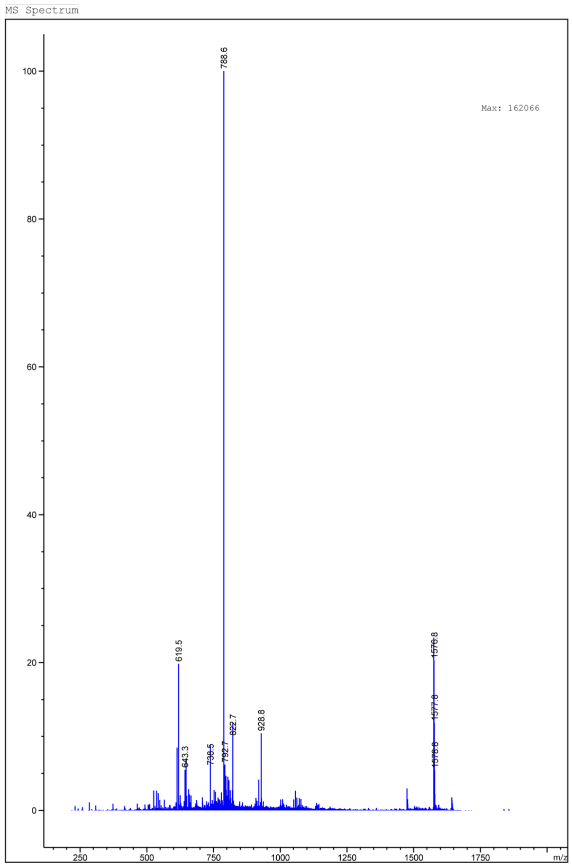


LCMS traces of CP-SW35A. (A) LC trace (B) MS

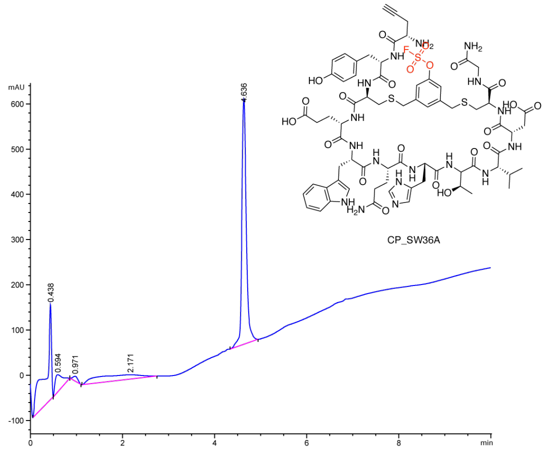


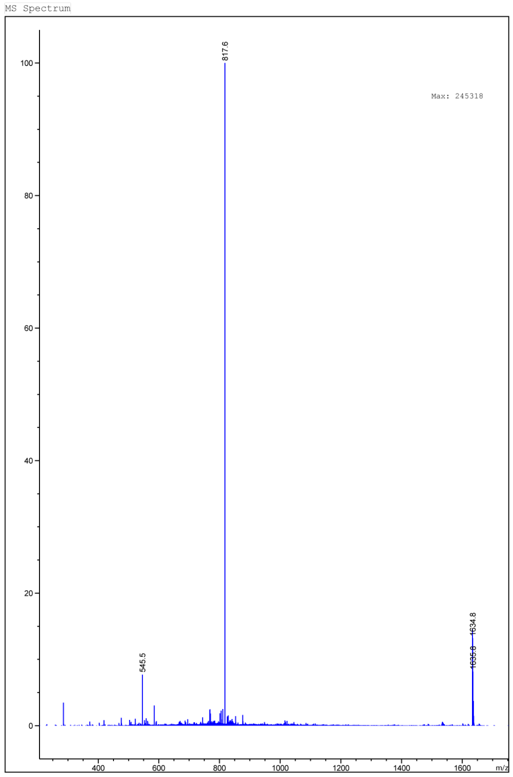


LCMS traces of CP-SW36A. (A) LC trace (B) MS

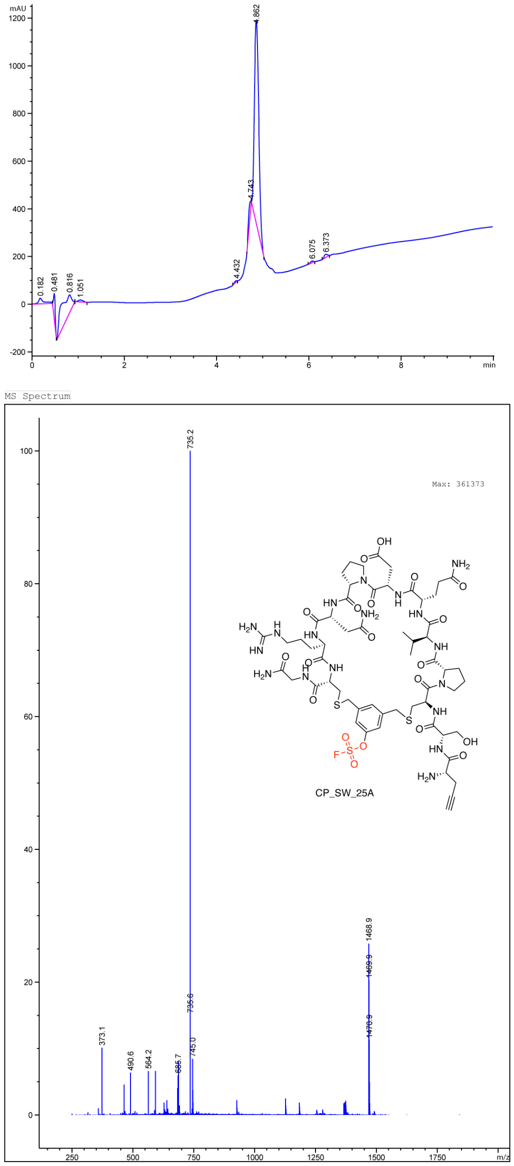


LCMS traces of CP-SW25A. (A) LC trace (B) MS

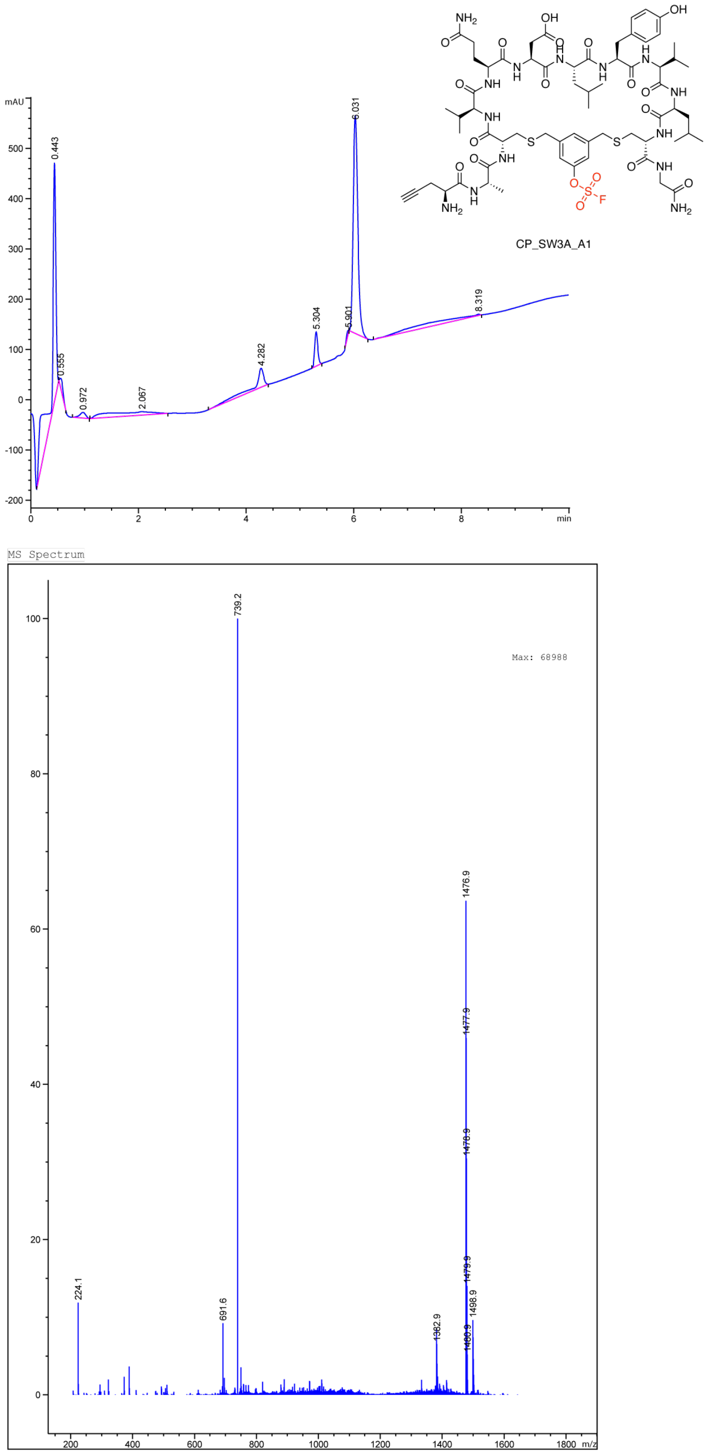


LCMS traces of CP_SW3A_A1. (A) LC trace (B) MS

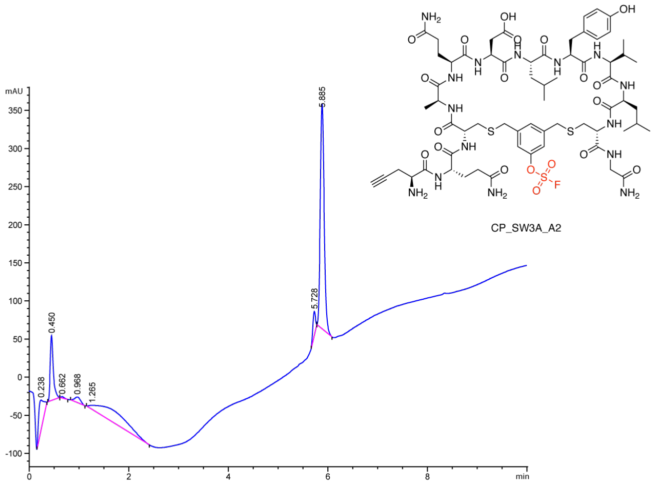


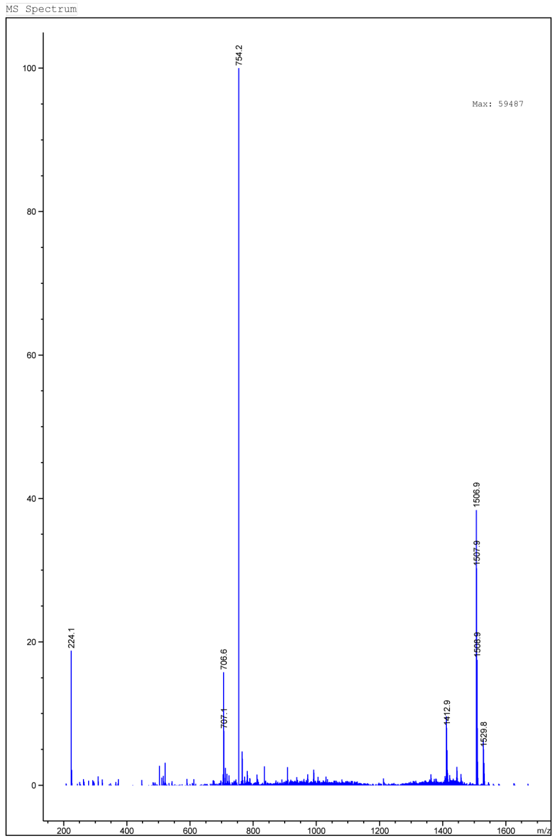


LCMS traces of CP_SW3A_A2. (A) LC trace (B) MS

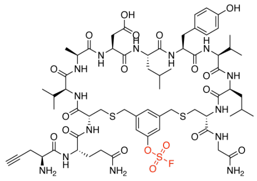

CP_SW3A_A3

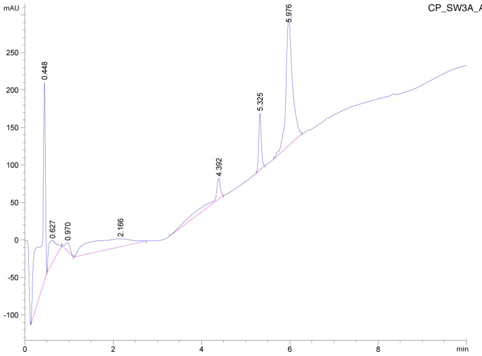


MS Spectrum

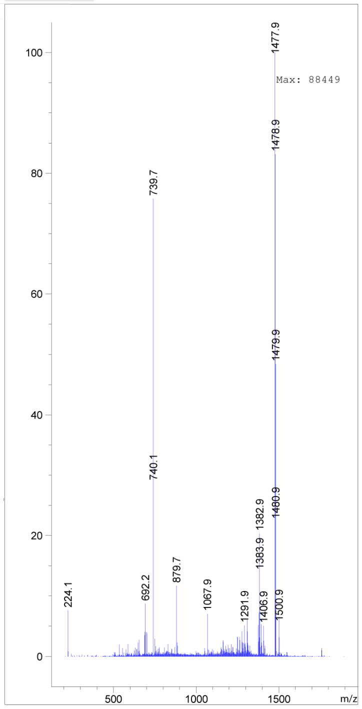


LCMS traces of CP_SW3A_A3. (A) LC trace (B) MS

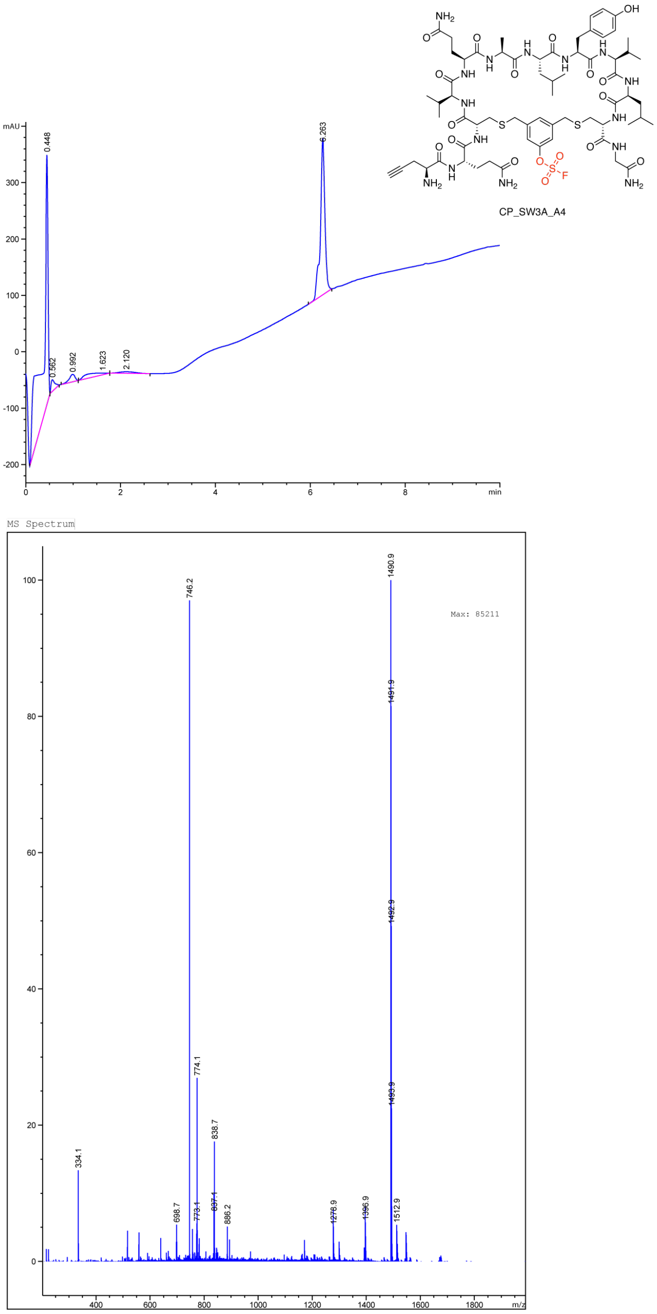


LCMS traces of CP_SW3A_A4. (A) LC trace (B) MS

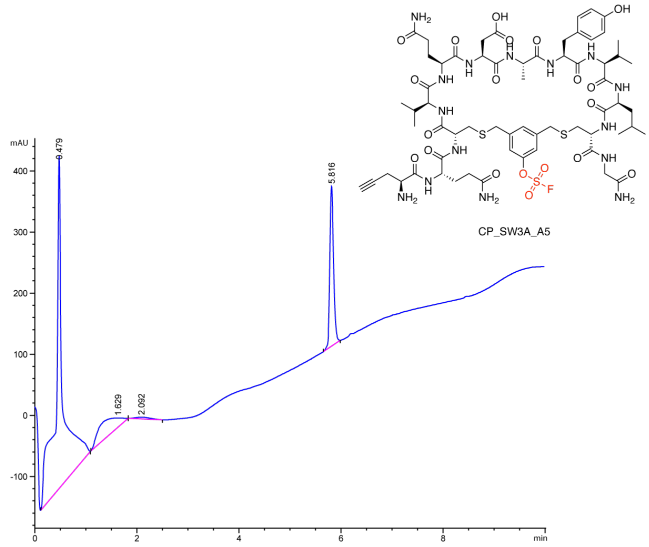


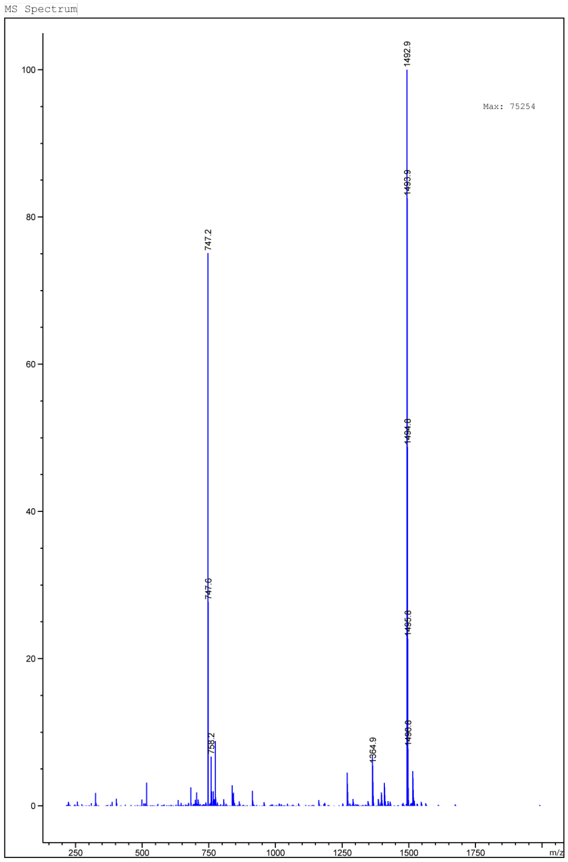


LCMS traces of CP_SW3A_A5. (A) LC trace (B) MS

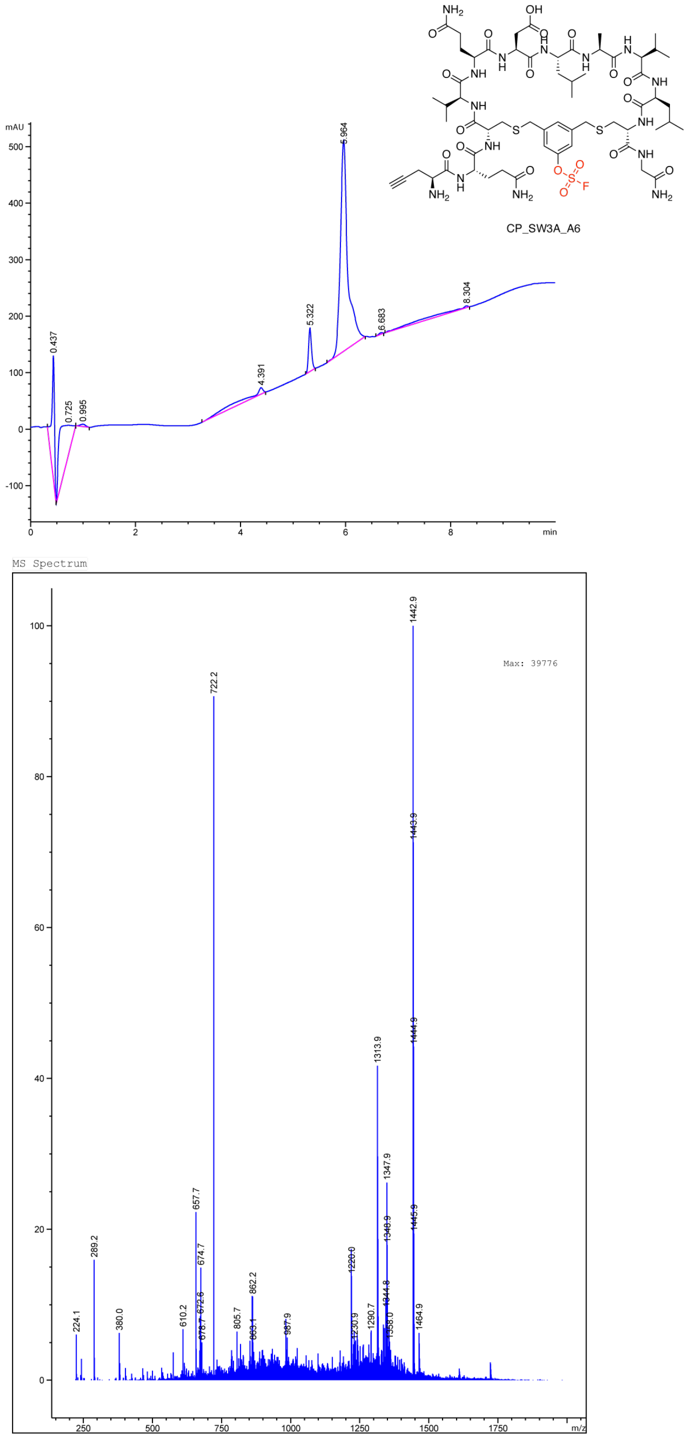


LCMS traces of CP_SW3A_A6. (A) LC trace (B) MS

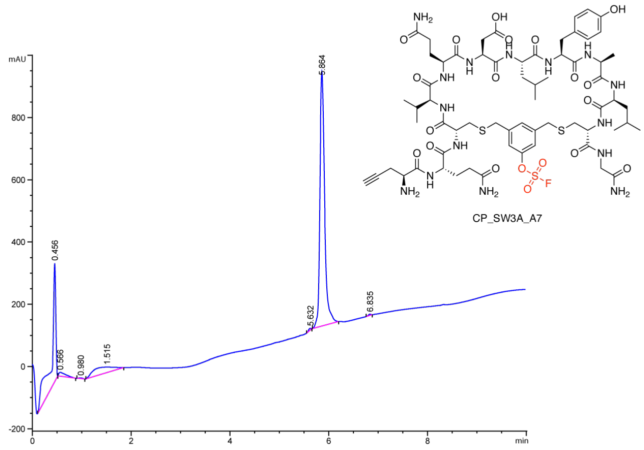

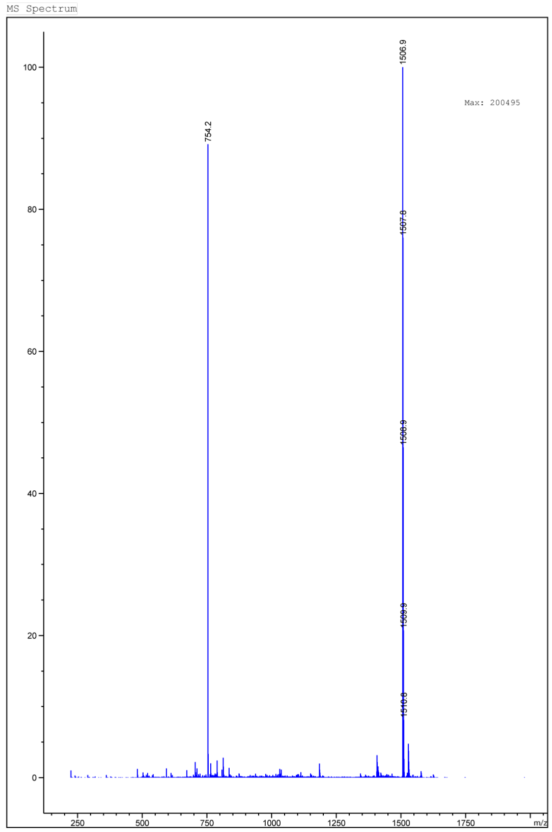

LCMS traces of CP_SW3A_A7. (A) LC trace (B) MS

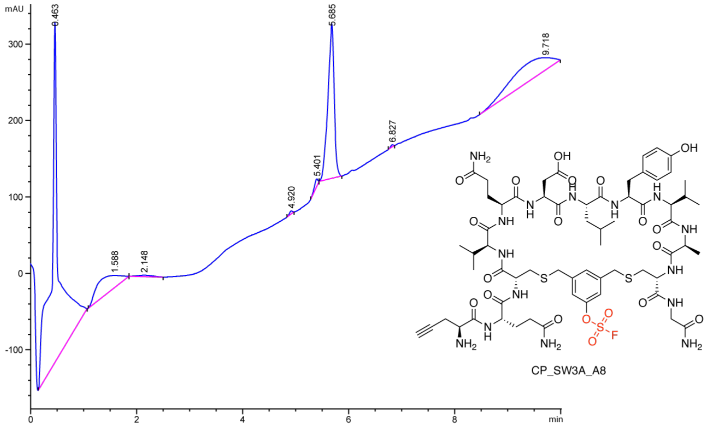


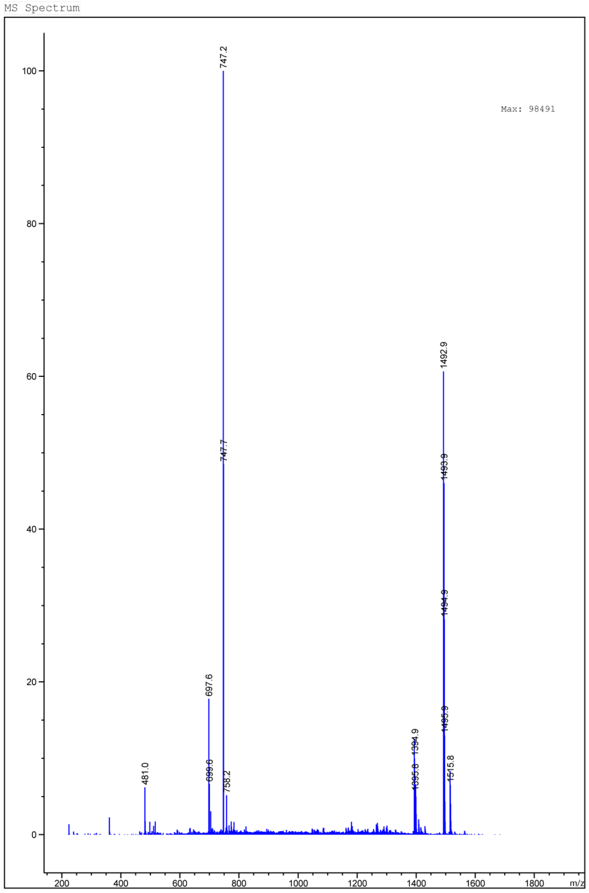


LCMS traces of CP_SW3A_A8. (A) LC trace (B) MS

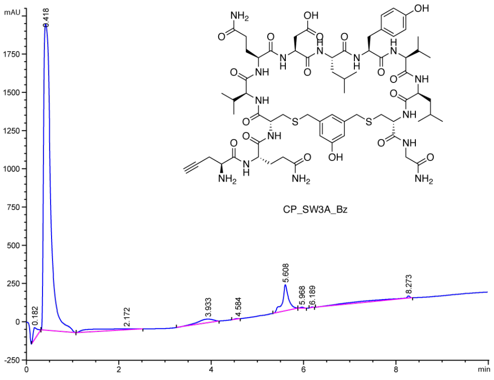


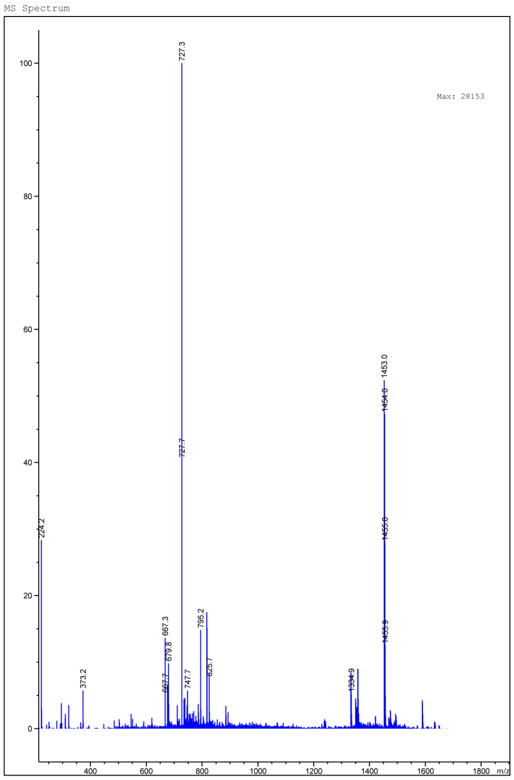


LCMS traces of CP_SW3A_Bz. (A) LC trace (B) MS

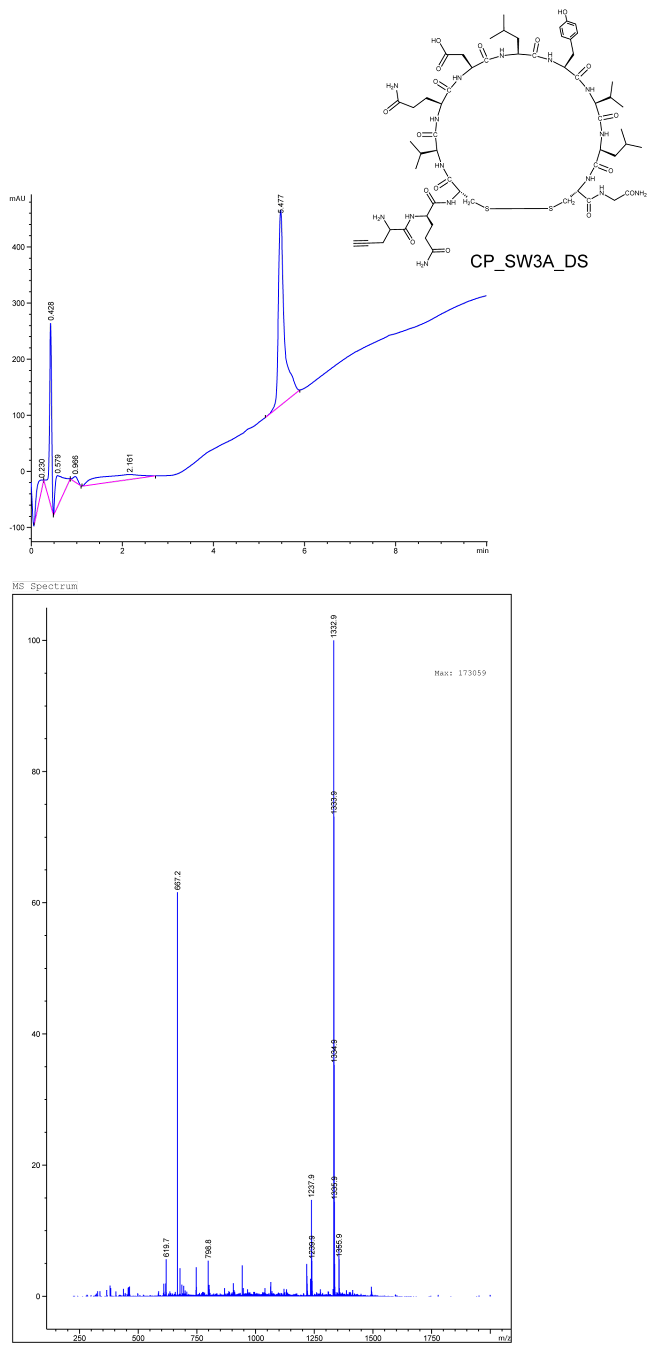


LCMS traces of CP_SW3A_DS. (A) LC trace (B) MS

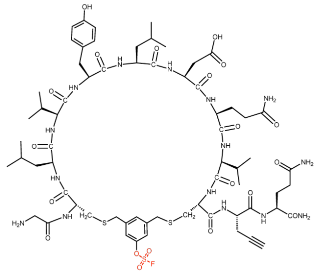

CP_SW3A_S1

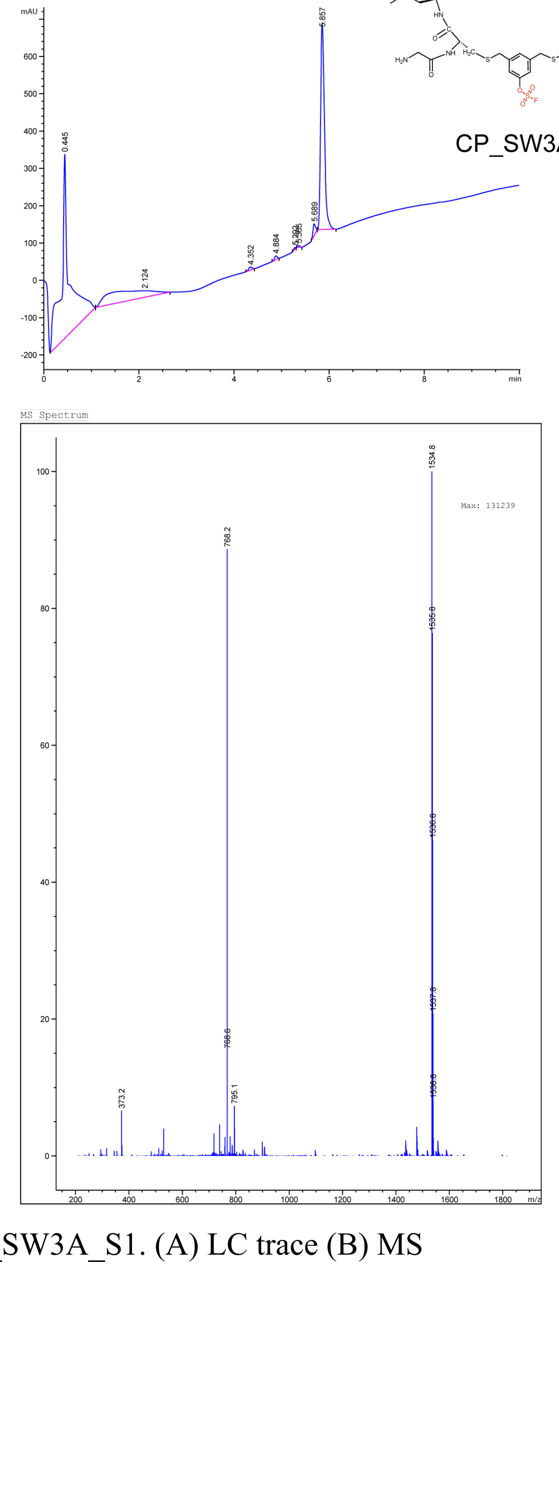


LCMS traces of CP_SW3A_S1. (A) LC trace (B) MS

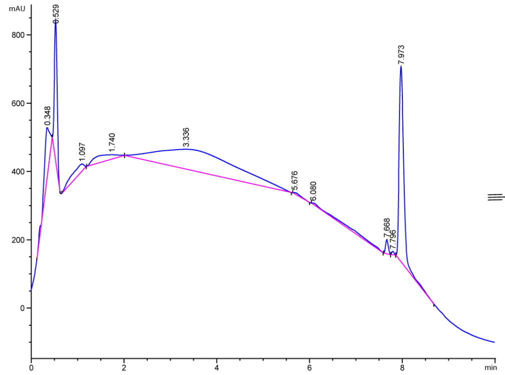


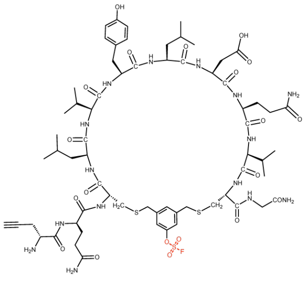

CP_SW3A_S2

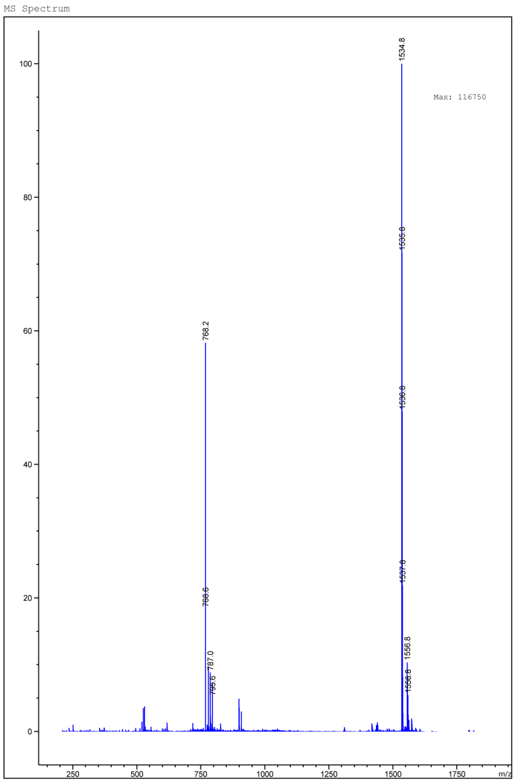


LCMS traces of CP_SW3A_S2. (A) LC trace (B) MS

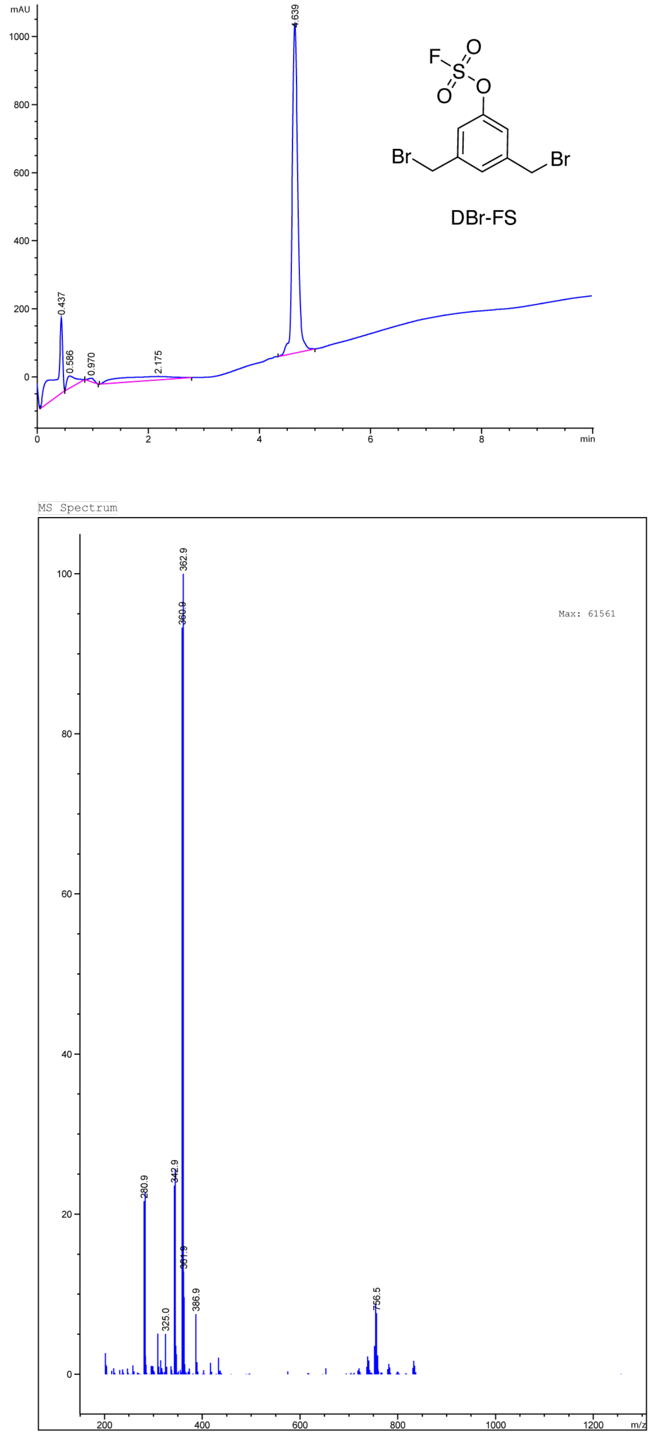


LCMS traces of DBr-FS linker. (A) LC trace (B) MS

## 6. NMR Spectra

$^{1}H$ NMR

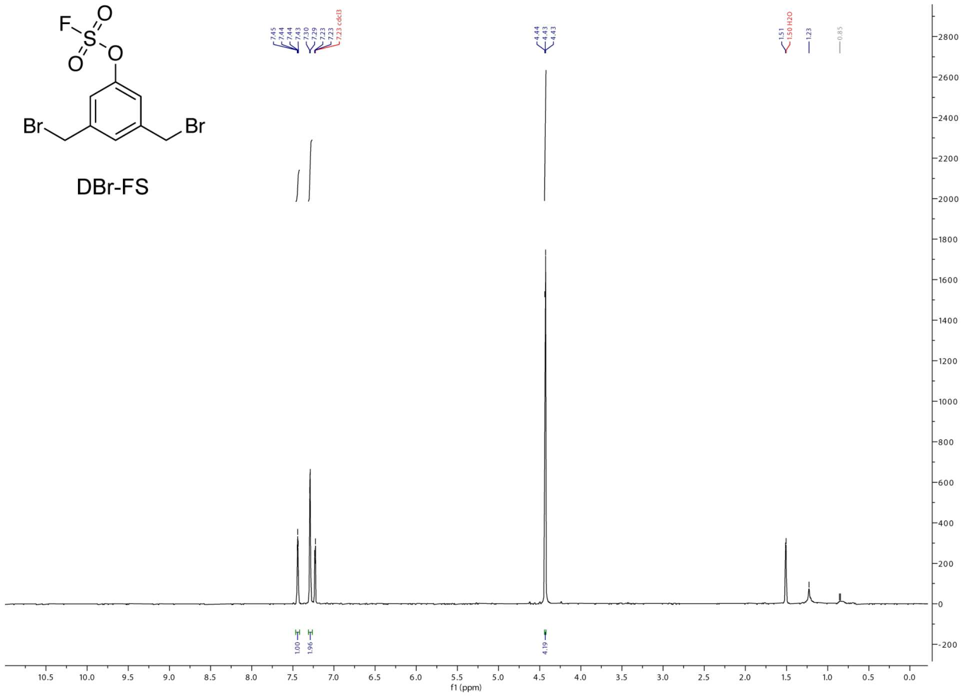

$^{19}F$ NMR

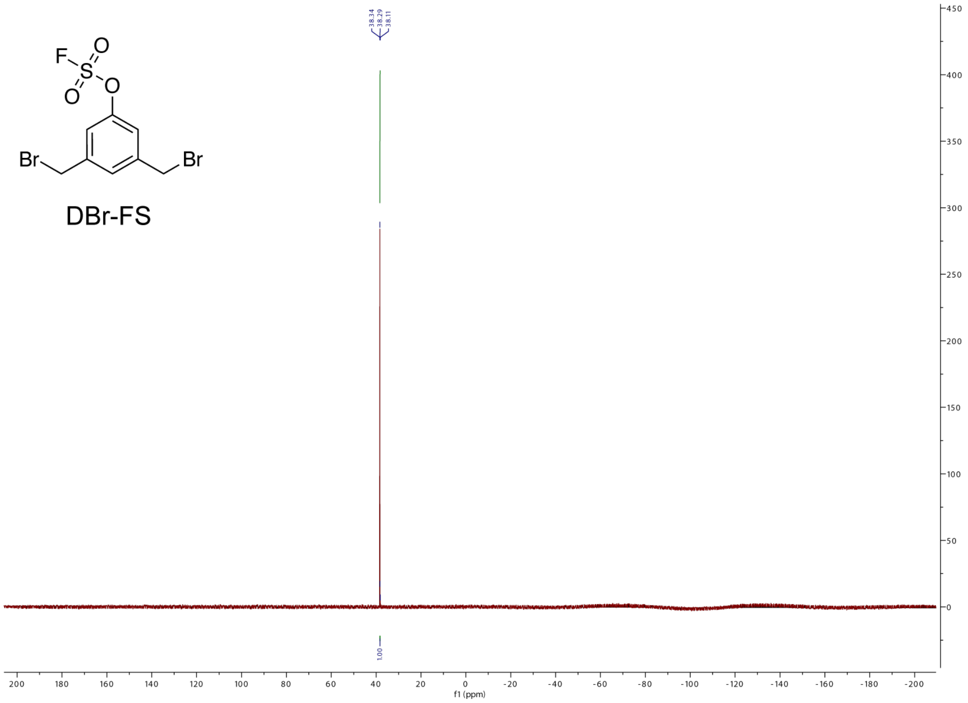
